# Additive Channels in Curved Fitness Landscapes

**DOI:** 10.64898/2026.03.21.713332

**Authors:** Daniel Ortiz-Barrientos, Mark Cooper

## Abstract

Fitness variation within a population depends both on how genetic differences affect reproduction and on how common those differences are. We examine this dependence using breeding values, which sum the average effects of an individual’s alleles on traits in that population. We average reproductive output at each breeding value before taking its logarithm, then ask how much of the resulting variation a straight line can describe. For normally distributed breeding values, a quadratic approximation separates variation due to slope from variation due to curvature. Holding the mean and fitness surface fixed, reducing all variances and covariances proportionally reduces the curved contribution faster. The linear share then rises when both contributions are positive. A large share defines an additive channel. That closer fit might suggest that curvature can also be ignored when predicting selection. We show why this inference can fail: selection weights breeding values by expected reproductive output as well as by their frequency. Curvature can still matter for selection even when it accounts for only a small share of variation in log expected fitness. Within the quadratic model, we bound the relative error from omitting curvature in the predicted shift of mean breeding value among selected parents. The same information about genetic variation and fitness can explain an additive channel and assess the accuracy of a selection prediction.

**Article summary:** Seed production may rise with plant height and then fall, yet a line can describe log expected fitness closely across breeding values in a population. These values sum average allelic effects on height in that population. The authors show how genetic variation changes the linear share, calling a large share an additive channel. A closer fit might seem to justify ignoring curvature when predicting selection. But reproductive output gives parents different weights, so ignoring curvature can mispredict the shift in mean breeding value among selected parents. They bound this error, allowing the prediction to be assessed separately from the fit.

## 1 Introduction

Seed production can rise with plant height and then fall. Yet a population may contain mostly plants from a range over which a line describes differences in expected reproduction closely. Another population, with the same mean but a wider range of heights, may show a larger contribution from curvature. What does the closer linear fit tell us about the first population? It describes the fitness differences among the individuals present, but its value for predicting their reproductive contributions needs a separate assessment. We develop that comparison for breeding values, making the genetic contribution to trait variation explicit.

For height, a plant’s breeding value sums the average effects of its alleles on height in the population (Fisher, 1918, 1941). A one-centimeter difference in breeding value means that the additive genetic predictions of height differ by one centimeter. Average effects can be defined even when alleles interact, so a breeding value describes the additive contribution to a trait in the present population without assuming additive gene action. Predicting change in offspring also requires a model of inheritance.

An additive genetic description of a trait does not require a linear relationship between that trait and reproduction. A related distinction arises in breeding: Technow et al. (2015) modeled how traits with additive genetic effects combine during maize growth to determine grain yield in different environments. Their study addressed yield prediction. Here the outcome is reproductive fitness, and the question is how the breeding values present affect its linear description.

The same breeding value can occur in individuals that differ in genotype, phenotype, growing conditions and reproductive success. To describe the relationship with breeding value, we therefore average over that variation. Let *W* denote nonnegative reproductive output during one selection episode, and **A** the random vector of breeding values for the traits of interest. At a particular value **a**, expected fitness is

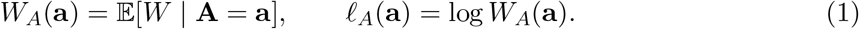

Taking the logarithm expresses proportional differences in this mean output: an increase from one expected offspring to two gives the same difference as an increase from ten to twenty. We call *l*_*A*_ *log expected fitness*. The logarithm gives the scale on which we describe fitness variation. Reproductive contributions themselves are weighted by output, a distinction that becomes important when we use the description to predict selection.

The expected output at a breeding value does not tell us how common that value is. Both quantities are needed to describe a population. For two traits, place one point per individual at its pair of breeding values. Fitness contours join pairs with equal expected output, whereas population density contours show where individuals occur. A combination with high expected fitness can still be rare. The density determines how frequently each fitness value enters a description of variation.

This population weighting can be read directly from a plot. Fit a straight line to log expected fitness against breeding value, giving each individual equal weight. A common breeding value receives more weight because more individuals carry it. Under the Gaussian, quadratic assumptions used below, the variation in this fitted line is the linear component of the total variation across the population. Appendix S7 illustrates the comparison with three one-trait populations on one curve before extending it to two traits.

We use *fitness landscape* for log expected fitness as a function of breeding value (Equation (1)). Each point averages over the genotypes, phenotypes and environments represented at that breeding value. The fraction of variance in a local quadratic approximation contributed by its linear term is the *first-order variance share*. A population and fitness surface with a large share form an *additive channel*; no universal cutoff is required. This definition concerns fitness across breeding values, not interactions among alleles. Two populations can have different conditional fitness relationships as well as different breeding-value frequencies. Holding the surface fixed lets us isolate the contribution of the frequencies.

This use of population frequencies follows classical selection theory. Lande averaged fitness over environmental deviations to obtain a function of breeding values (Lande, 1979). Lande and Arnold fitted linear and quadratic descriptions of relative fitness across the observed phenotype distribution and separated their contributions to fitness variance (Lande and Arnold, 1983). Subsequent work developed these variance comparisons (Moorad and Wade, 2013) and related log-link fitness models to selection gradients (Morrissey and Goudie, 2022). We extend the comparison to log expected fitness conditional on breeding value. This makes it possible to ask how a change in population covariance alters the linear share on a fixed surface, and whether that share also indicates the accuracy of a selection prediction.

A proportional change in covariance provides a controlled comparison: the mean and fitness relationship can stay fixed while the range of breeding values changes. If the linear fit improves, the next question is whether curvature can also be omitted when calculating parental reproductive contributions. That calculation requires expected-output weights in addition to population frequencies. To follow the resulting covariance into offspring, we then introduce a separate inheritance model, using established results on selection-generated linkage disequilibrium and its reduction by recombination (Bulmer, 1971; Barton and Turelli, 1991).

Each comparison requires its own assumptions, collected in Table 1. The appendices give the proofs and address interpretation and estimation (Appendix S6); Supplementary File 1 develops the one-trait calculations before introducing several traits.

**Table 1.**
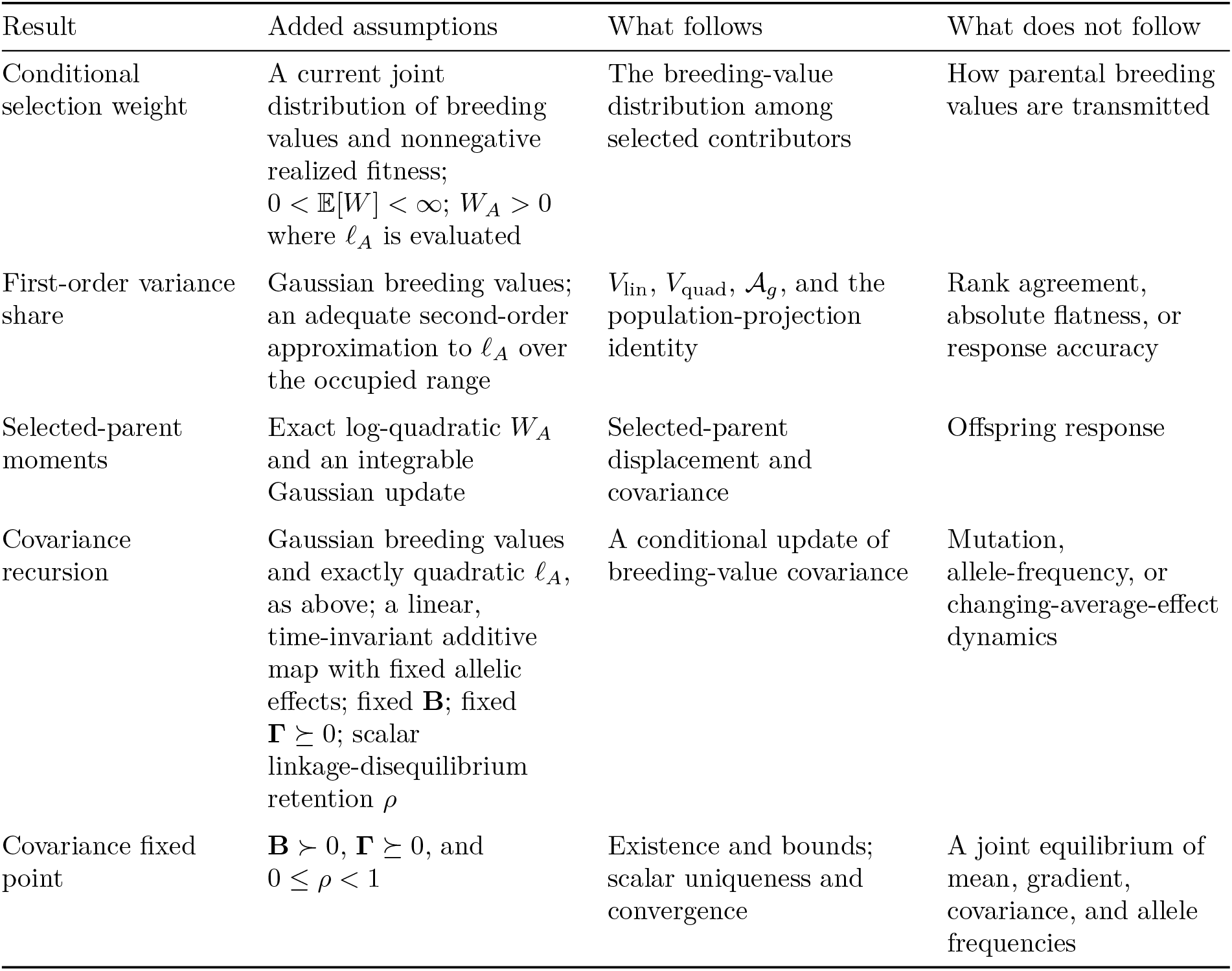
Assumptions and conclusions. Each row adds assumptions to those above it only where stated. The table separates current-state selection from transmission to offspring. Symbols are defined where the text first uses them: *l*_*A*_ in Equation (1), ***b*** and **H** in Equation (5), *V*_lin_, *V*_quad_, and *A*_*g*_ in Equations (7)–(9), *W*_*A*_ in Equation (1).

| Result | Added assumptions | What follows | What does not follow |
|---|---|---|---|
| Conditional selection weight | A current joint distribution of breeding values and nonnegative realized fitness; 0 < 𝔼[*W*] < ∞; *W*<sub>*A*</sub> > 0 where *ℓ*<sub>*A*</sub> is evaluated | The breeding-value distribution among selected contributors | How parental breeding values are transmitted |
| First-order variance share | Gaussian breeding values; an adequate second-order approximation to *ℓ*<sub>*A*</sub> over the occupied range | *V*<sub>lin</sub>, *V*<sub>quad</sub>, 𝒜<sub>*g*</sub>, and the population-projection identity | Rank agreement, absolute flatness, or response accuracy |
| Selected-parent moments | Exact log-quadratic *W*<sub>*A*</sub> and an integrable Gaussian update | Selected-parent displacement and covariance | Offspring response |
| Covariance recursion | Gaussian breeding values and exactly quadratic *ℓ*<sub>*A*</sub>, as above; a linear, time-invariant additive map with fixed allelic effects; fixed **B**; fixed **Γ** ⪰ 0; scalar linkage-disequilibrium retention *ρ* | A conditional update of breeding-value covariance | Mutation, allele-frequency, or changing-average-effect dynamics |
| Covariance fixed point | **B** ≻ 0, **Γ** ⪰ 0, and 0 ≤ *ρ* < 1 | Existence and bounds; scalar uniqueness and convergence | A joint equilibrium of mean, gradient, covariance, and allele frequencies |

## 2 Conditional expected fitness

The conditional mean in Equation (1) is the expected reproductive contribution of individuals with a given breeding value. Combining it with the frequency of that value identifies the distribution among reproductive contributors. This establishes the biological role of *W*_*A*_ before we use its logarithm to describe fitness differences. No assumption about the shape of the breeding-value distribution is needed at this stage (Table 1, first row).

### 2.1 Average first, then take the logarithm

How much a breeding value contributes to reproduction depends on both its frequency and the expected output of its carriers. Let *p*_*A*_(***a***) denote its density before selection. Multiplying that density by the conditional expected output *W*_*A*_(***a***) gives its reproductive contribution; dividing by population mean fitness gives the selected-parent density:

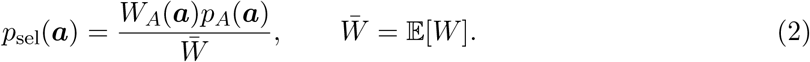

In this selected distribution, breeding values associated with greater expected output receive more weight. Equation (2) describes the parental breeding values represented by reproductive contributions. It does not yet describe the breeding values of offspring, because that also depends on what the parents transmit.

For the fitness description, we compare proportional differences in expected output. Where *W*_*A*_(***a***) *>* 0, the log difference between two breeding values is

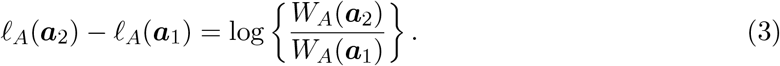

The gradient of *l*_*A*_ is therefore the local proportional change in expected output per unit change in breeding value. Multiplying every fitness value by the same positive constant adds a constant to *l*_*A*_ and leaves its derivatives and variances unchanged. Exponentiating a quadratic log mean also keeps expected output positive, as in a log-link model (Morrissey and Goudie, 2022). A line fitted on this scale describes ratios of expected output; its explained variance need not equal the explained variance on the original scale.

The order of the operations matters. We first average reproductive output at a breeding value and then take the logarithm. Reversing that order generally gives a different quantity, by Jensen’s inequality:

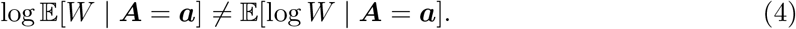

For example, individuals with the same breeding value might leave zero and two offspring. Their mean output is one and its logarithm is zero, but their mean log output is undefined. Expected log growth has a different purpose when population growth factors multiply across generations (Lewontin and Cohen, 1969). Our target concerns reproductive output averaged within one selection episode, including individuals that leave no offspring.

### 2.2 Slope and curvature near the population mean

To ask how closely a line describes this log mean across the population, we need both its slope and the way that slope changes. Let ***a***_0_ = E[***A***] and ***X*** = ***A****−* ***a***_0_, so E[***X***] = **0** and Var(***X***) = **G**. If *ω*_*A*_ is twice continuously differentiable near ***a***_0_, its second-order approximation is

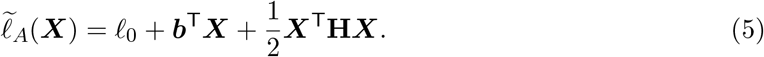

Here *l*_0_ = *l*_*A*_(***a***_0_) is log expected fitness at the mean. The gradient ***b*** = ∇*l*_*A*_(***a***_0_) gives the slope for each trait, and the Hessian **H** = ∇^2^*l*_*A*_(***a***_0_) describes how those slopes change. The linear term assigns a constant slope across breeding values; the quadratic term accounts for changes in slope. Both derivatives refer to the current mean and may change when the mean or environment changes.

The range represented by the population determines whether this approximation is useful. Twice differentiability gives a local expansion, but does not ensure that higher-order terms are small over all breeding values that occur. An empirical application must therefore check the quadratic fit across the population’s realized distribution, rather than only near its mean.

Figure 1 shows the two averages involved. In panel a, individuals with the same breeding value have different phenotype-dependent fitness values. Each open circle is a sample mean, and the curve is the corresponding conditional expectation *W*_*A*_. The groups are placed at evenly spaced breeding values to illustrate averaging; their horizontal frequency is not the population distribution. The shaded density in panel b supplies that distribution and hence the weights used to calculate variation along the log-fitness curve.

**Figure 1:**
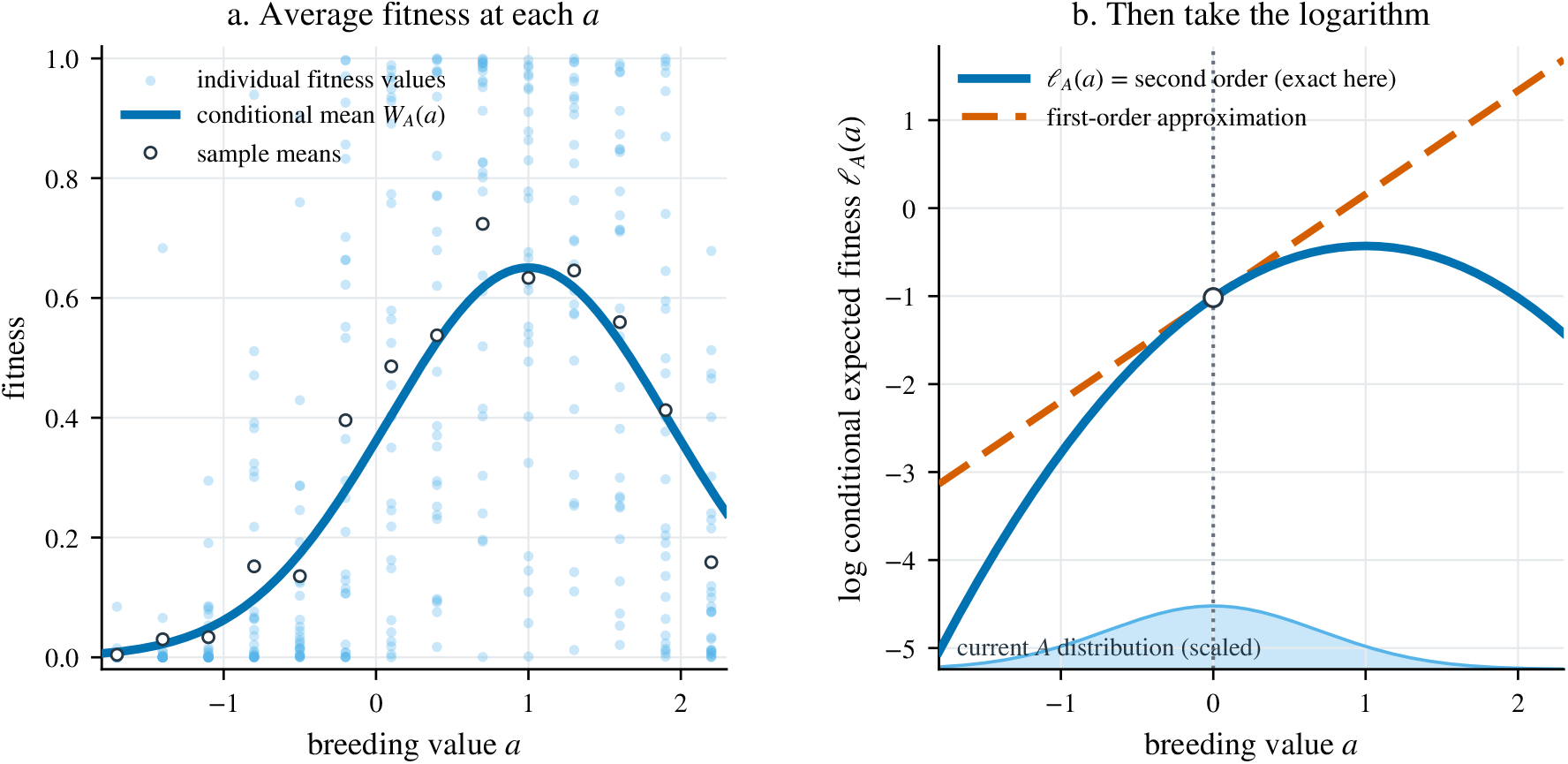
From phenotype-dependent fitness to expected fitness at each breeding value. (a) Small filled points show phenotype-dependent fitness values for individuals that share a breeding value but differ in environmental deviation. Each open circle marks the sample mean at one breeding value. The dark curve gives the corresponding population expectation, *W*_*A*_(*a*) = E[*W* |*A* = *a*] (Equation (1)). (b) The logarithm of this expectation is shown with its tangent and quadratic approximation at the population mean. The shaded density gives the frequency of breeding values in the population and hence their weight in the variance calculation. The equal-sized samples in panel a were chosen to illustrate averaging; variation in offspring counts is not simulated. *Alt text:* Filled points show individual fitness and open circles their sample means. A smooth curve shows expected fitness at each breeding value. The second panel shows the log of that expectation, its tangent at the population mean, and the breeding-value density.

The remaining calculation is to evaluate the linear and quadratic terms at the breeding values in this distribution. Each individual contributes to their variances. Breeding values with greater population frequency receive correspondingly greater weight, linking the fitted fitness relationship to the standing variation in the population.

## 3 Population variation and curvature

We now ask how much of the population’s variation in log expected fitness can be described by the linear term. A Gaussian breeding-value distribution and an adequate quadratic approximation allow the linear and curvature contributions to be separated (Table 1, second row).

### 3.1 First-order and quadratic variation

Assume that ***X*** *∼N*(**0, G**). Once the mean is fixed, **G** specifies the Gaussian population and therefore the frequency weights in both variance calculations. Write the linear term as *L* = ***b***^⊤^***X*** and subtract the mean from the quadratic term to give

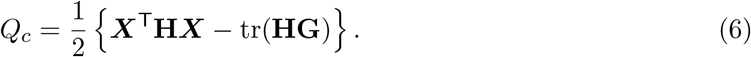

Subtracting this mean changes only the intercept. Gaussian symmetry makes *L* and *Q*_*c*_ uncorrelated, so their variances add. The linear- and quadratic-form moments give

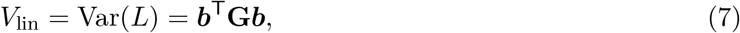

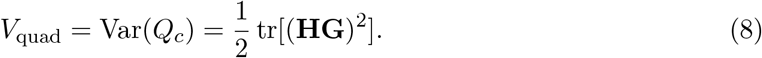

The two expressions quantify variation in the fitness description: *V*_lin_ comes from slope and *V*_quad_ from curvature. Both depend on the breeding-value covariance, but neither is the additive genetic variance of the trait. Likewise, the quadratic component describes curvature of log expected fitness and is not a measure of epistatic genetic variance (Cheverud and Routman, 1995).

Their sum is the variance of the quadratic approximation. Provided *V*_lin_ + *V*_quad_ *>* 0, the fraction contributed by the linear term is

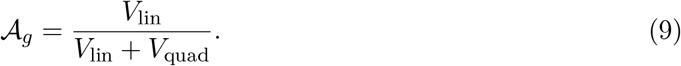

For example, *A*_*g*_ = 0.8 means that the linear term accounts for 80% of the variance in the quadratic approximation and curvature accounts for 20%. The subscript *g* identifies the current breeding-value distribution used in this comparison. This is a relative measure of log expected fitness: the same fraction can describe two large variance components or two small ones. Variation in realized reproductive output includes further sources and is not the denominator of Equation (9).

The boundary cases clarify what the fraction measures. If *V*_lin_ = 0 *< V*_quad_, then *A*_*g*_ = 0; if *V*_quad_ = 0 *< V*_lin_, then _*g*_ = 1. When both terms vanish, there is no variance to partition and the ratio is undefined. Singular **G** causes no diffculty for Equations (7)–(9). Statements involving inverses below are restricted to its support, the subspace with nonzero breeding-value variance.

### 3.2 Connection to linear regression

The share can also be obtained by fitting the line used in the introductory comparison. Fit a line or plane, including an intercept, to the quadratic approximation under the population’s frequency weights. The population least-squares fit is

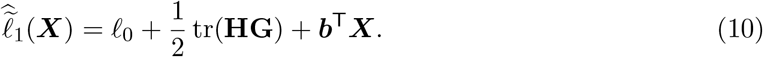

The fitted slope is the local gradient ***b***, while the intercept includes the mean contribution of curvature. Because the remaining centered quadratic term is uncorrelated with the linear term, the population cofficient of determination is

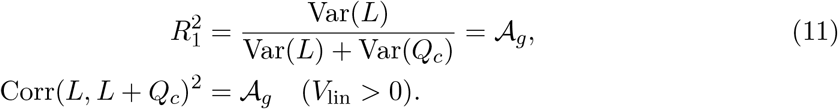

The same comparison can be made for a noisy log-scale outcome under a specified observation model. If 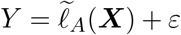 with E[*ε* ***X***] = 0 and E[*ε*^2^ ] *<∞*, the linear and quadratic population regressions share the same total-variance denominator. Their ratio therefore isolates the linear fraction of the fitted quadratic variation:

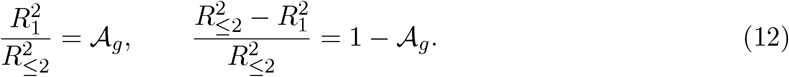

This identity applies to population least-squares regressions under the stated model. Generalized-linear-model pseudo-*R*^2^ values and arbitrary nonlinear sample fits do not automatically share it. When *V*_lin_ = 0 *< V*_quad_, the fitted line is constant and *A*_*g*_ = 0; the correlation expression is then undefined because the linear predictor has no variance.

Figure 2 checks the population identity against finite samples for 360 combinations of slope, curvature and covariance. The analytic share agrees with the squared correlation estimated from Gaussian samples, with median absolute difference 0.00237. Having identified the share with the linear fit, we can now vary the population while holding the fitness surface fixed.

**Figure 2:**
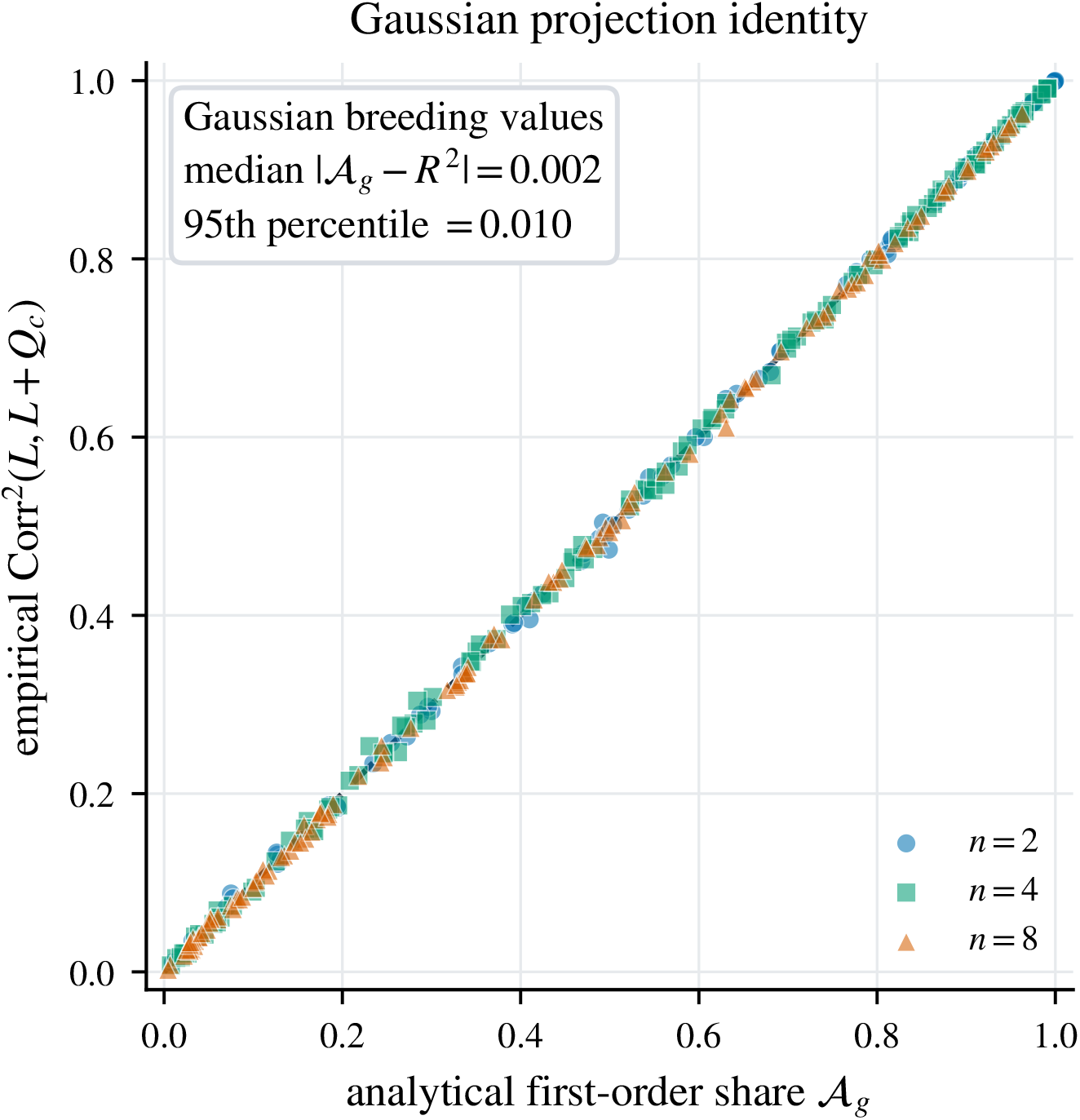
How closely does a finite sample reproduce the predicted linear share? We generated 360 current-state triples (***b*, H, G**) in dimensions 2, 4, and 8 and sampled 20,000 Gaussian breeding-value vectors per triple (seed 20260905). Eigenvalues of **G** ranged from 0.08 to 2.5, absolute eigenvalues of **H** from 0.03 to 0.9, and ||***b***|| from 0.08 to 1.5; both stabilizing and saddle-shaped curvature were included. The horizontal axis is analytic *A*_*g*_, and the vertical axis is the squared sample correlation between *L* and *L* + *Q*_*c*_. Dimensions are distinguished by symbol and the line is equality. The median absolute difference was 0.00237 and its 95th percentile was 0.01033. The calculation checks Equation (11); it does not test ranking or selected-parent displacement. *Alt text:* A scatterplot places 360 points close to the diagonal equality line. The horizontal axis is the analytic first-order variance share and the vertical axis is its squared-correlation estimate. Different symbols identify dimensions 2, 4, and 8.

### 3.3 Changing the breeding-value distribution

Consider two populations with the same mean and the same conditional fitness relationship, but different amounts of breeding-value variation. To change only that amount, set **G** = *s***G**_0_ for a positive scalar *s*, keeping the local slope ***b*** and curvature **H** fixed. All variances and covariances change in the same proportion, preserving the distribution’s shape. The two fitness-variance components then change at different rates:

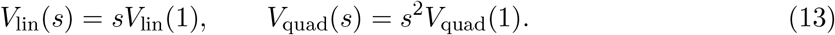

Halving covariance halves the linear variance but reduces the quadratic variance to one quarter. Thus, when both components are positive, a line describes a larger fraction of fitness variation even though the fitness surface and population mean have not changed. This stronger linear description occurs while the absolute linear variance is falling. More generally, let *L*_0_ = *V*_lin_(1) and *Q*_0_ = *V*_quad_(1). Then *A*_*g*_(*s*) = *L*_0_*/*(*L*_0_ + *sQ*_0_) and the quadratic share is *sQ*_0_*/*(*L*_0_ + *sQ*_0_). The quadratic *variance* scales as *s*^2^; its fraction of total variance does not.

Figure 3 makes this population effect visible. The same gray log expected-fitness contours apply to both populations. The blue solid ellipse contains 95% of the baseline population, for which *A*_*g*_ = 0.604. The orange dashed ellipse contains 95% of a population with the same mean and 0.11 times the baseline covariance, for which *A* _*g*_ = 0.933. The narrower population gives less weight to deviations where the quadratic term contributes strongly. Holding the conditional relationship fixed isolates that frequency effect; a biological change in variance need not leave the relationship unchanged.

**Figure 3:**
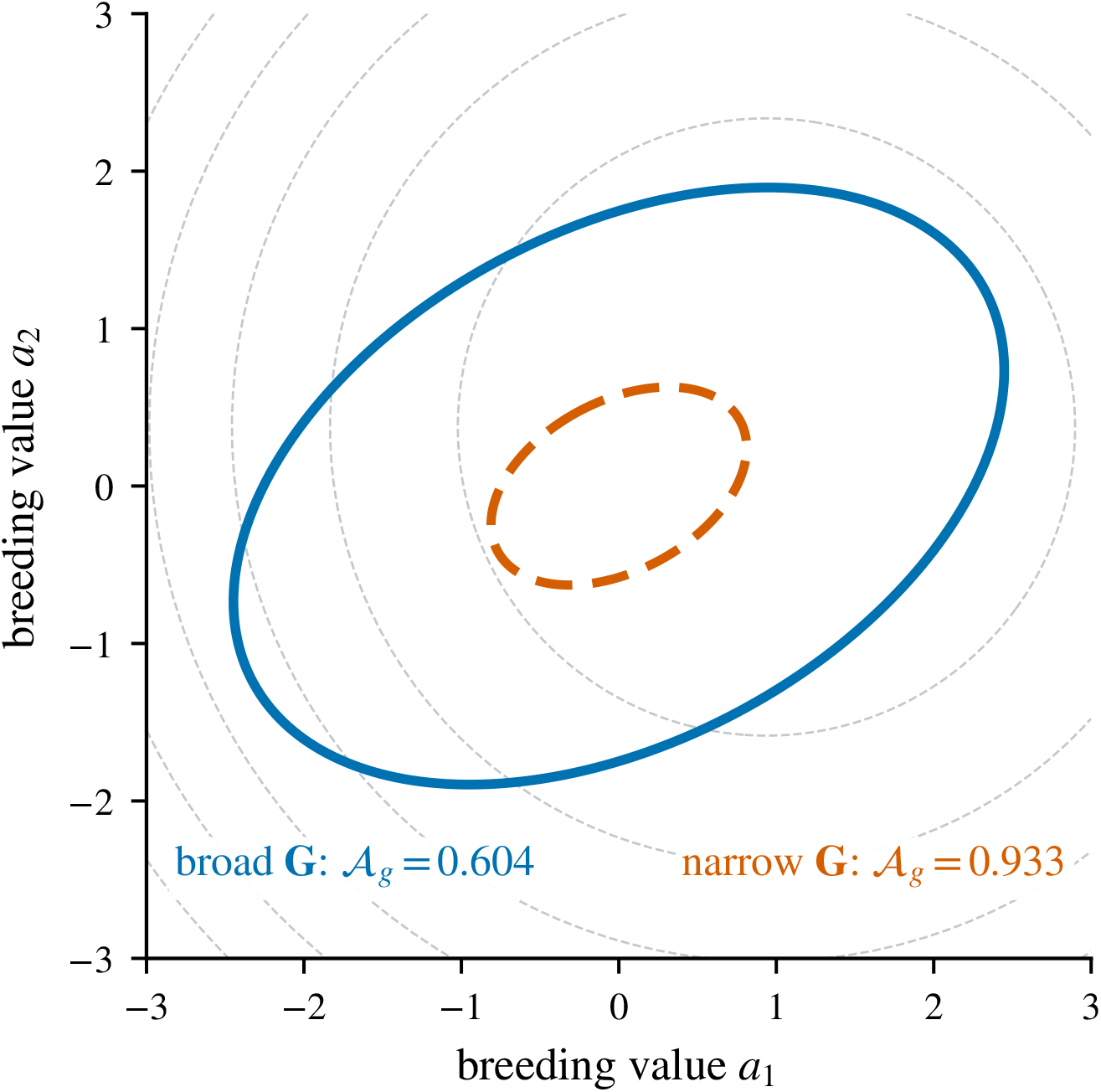
Two populations on the same fitness surface. Gray contours join pairs of breeding values with the same log expected fitness; the slope at the mean is *b* = (0.75, 0.30) ^⊤^ and the curvature is H = 0.8I. The larger blue solid-line ellipse encloses 95% of the baseline population (*A*_*g*_ = 0.604). The smaller orange dashed-line ellipse encloses 95% of the population with covariance 0.11**G** (*A* _*g*_ = 0.933). Both have the same mean. Reducing all variances and covariances by the same proportion reduces the linear contribution by a factor of 0.11 and the quadratic contribution by a factor of 0.11^2^ (Equation (13)). *Alt text:* On one set of gray fitness contours, a larger blue ellipse with a solid outline encloses a smaller orange ellipse with a dashed outline. They have the same center. The shares are 0.604 for the blue ellipse and 0.933 for the orange ellipse.

This conclusion depends on proportional change. If variation falls mainly along the gradient, the linear contribution can decline enough to lower the share. For example, take ***b*** = (1, 0) ^⊤^ and **H** = −**I**. Changing **G** = **I** to diag(0.1, 0.9) reduces both trait variances, but reduces variation along the gradient most strongly. The share falls from 0.5 to 0.1*/*0.51 ≃ 0.196. The direction in which genetic variation changes therefore matters as well as its total amount.

### 3.4 Changing coordinates and changing orientation

A change in the population must be distinguished from a change in how its traits are measured. Changing trait units, or applying another invertible linear coordinate transformation, leaves both variance components and *A*_*g*_ unchanged (Appendix S1). A useful transformation scales and rotates the coordinates so that breeding-value covariance becomes the identity: each transformed variable has unit variance and the variables have zero pairwise covariance. This is *whitening*. Define

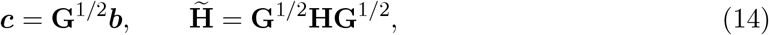

Then *V*_lin_ = *∥****c*** *∥*^2^ and 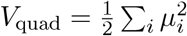, where *µ*_*i*_ are the eigenvalues of 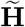. The first component depends on the length of the standardized gradient and the second on the curvature eigenvalues. Neither pairs a coordinate of ***c*** with a curvature eigenvalue. Rotating the gradient in these standardized coordinates therefore leaves *A*_*g*_ unchanged when its norm and the curvature spectrum are held fixed.

Physically changing the direction of the population’s variation on fixed trait axes is a different operation. For a unit vector ***u***, the breeding-value variance in that direction is ***u*** ^⊤^**G*u***, its directional evolvability (Hansen and Houle, 2008). The linear fitness variance is the variance along the gradient multiplied by the squared gradient magnitude. With gradient magnitude and covariance fixed, *V*_lin_ is greatest when the gradient points along the direction of greatest breeding-value variance, connecting the comparison to genetic lines of least resistance (Schluter, 1996). Curvature’s orientation relative to the covariance axes can likewise alter *V*_quad_. These comparisons change how ***b*** or **H** lies relative to **G**. They need not preserve the standardized gradient length or curvature spectrum. Appendix S7 illustrates the population-orientation comparison.

The fitness surface alone therefore does not determine the quality of the linear fit. A narrower population can have a larger share on the same surface, and a change in orientation can alter either component. Reporting *V*_lin_, *V*_quad_ and the population covariance alongside *A*_*g*_ identifies what changed. The remaining question is whether the resulting fit is accurate enough to replace the curved relationship when predicting reproductive contributions.

## 4 Within-generation selection

To use a fitted line for selection, we must compare the reproductive contributions predicted by the line with those predicted by the curved relationship. We now assume that log expected fitness is exactly quadratic over the Gaussian breeding-value distribution and that the population mean fitness is finite (Table 1, third row). This stronger assumption lets us calculate the selected-parent moments exactly, rather than infer them from the quality of a local approximation.

Equation (2) weights the initial distribution by expected reproductive output. Writing **Γ** = −**H**, that weight is

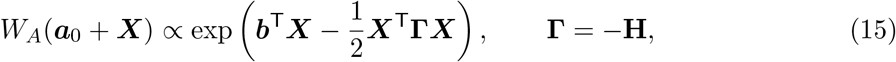

Here **Γ** = −**H** is the sign-reversed curvature of log expected fitness. If **G**^*−*1^ + **Γ ≻** 0, multiplying the initial Gaussian density by this weight and normalizing gives another Gaussian distribution.

Its mean displacement from the preselection mean and its covariance are

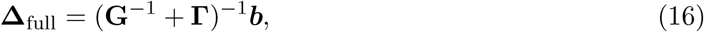

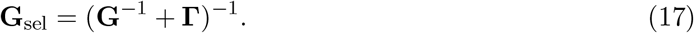

These are moments of parental breeding values weighted by reproductive contributions. They specify the selected parents, while offspring moments also depend on inheritance. Under stabilizing selection, **Γ ⪰** 0, the weighted covariance is no greater than the initial covariance in any trait direction: **G**_sel_ ⪯ **G**. With saddle-shaped curvature, the same formulas apply if **G**^−1^ + **Γ** ≻ 0, but selected covariance need not decrease.

We can now assess the line by the selected-parent mean it predicts. Omitting curvature gives**Δ** _lin_ = **G*b***. In one trait, take *G* = 1, Γ = 0.5, and *b* = 2. The same population then gives

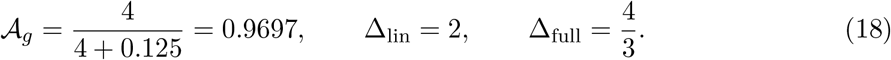

The line accounts for roughly 97% of the variance in log expected fitness, but predicts a selected-parent displacement 50% larger than the exact value. Increasing |*b* |brings *A*_*g*_ still closer to one without reducing this relative error. The source of the discrepancy is the change of weights. The variance calculation weights squared log-fitness deviations by the frequency of breeding values; the selection calculation additionally weights each value by expected reproductive output. Curvature can therefore have a small relative contribution to the first calculation and still matter to the second.

The selection prediction needs a measure of the absolute curvature represented in the population, rather than only its fraction of log-fitness variance. For a nonsingular covariance **G** *≻* 0 and a nonzero standardized gradient **G**^1*/*2^***b***, define

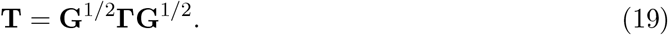

The norm 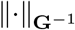 measures displacement in units set by the current breeding-value covariance. In these units, Appendix S2 bounds the error from omitting curvature relative to the full selected-parent displacement:

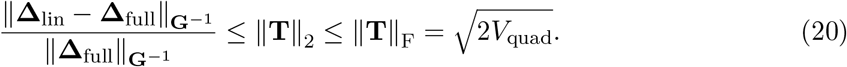

The first bound, ∥ **T**∥ _2_, is the largest absolute curvature after directions are scaled by population covariance. The second uses the absolute quadratic variance. If the standardized gradient is an eigenvector of **T** with eigenvalue *τ*_*k*_, the relative error is exactly |*τ*_*k*_| . Thus the gradient’s direction relative to curvature can affect displacement error even when its standardized norm and the curvature eigenvalues are fixed, the comparison under which *A*_*g*_ is unchanged. If **G** is singular, the inverses and norms are restricted to its positive-variance subspace and the gradient must have a nonzero component there.

The first-order share and Equation (20) now answer the two questions separately: how well a line describes log expected fitness, and how much selected-parent displacement can change when curvature is omitted. Both concern the current population. Following its covariance into the next generation requires an account of how selected parents transmit their genetic variation.

## 5 Covariance across generations

The selected-parent covariance is an input to reproduction, not a description of the offspring covariance by itself. Selection can alter associations among alleles in contributing parents, and recombination and segregation determine how much of those associations persists in offspring. We model that step explicitly, adding the assumptions in the final two rows of Table 1.

Assume additive gene action with fixed allelic effects, so the same linear map from inherited allelic content defines breeding value in every generation. Let **B≻** 0 be the genic covariance contributed by loci before accounting for linkage disequilibrium or other associations between them. Hold this baseline fixed and retain a fraction 0 ≤*ρ<* 1 of the selected covariance’s departure from it after reproduction. With fixed stabilizing curvature **Γ** ⪰0, the resulting update is

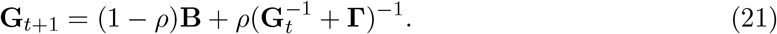

The parameter *ρ* specifies how much of the covariance departure generated by selection remains after reproduction. Freely recombining contributions in an infinitesimal, random-mating population give the benchmark *ρ* = 1*/*2. Recombination rates differ among locus pairs and mating systems differ among populations, so using one retention fraction is itself a summary of those differences (Bulmer, 1971, 1974; Barton and Turelli, 1991).

Fixing allelic effects does not by itself fix the genic baseline. Fixed effects keep breeding-value coordinates comparable across generations, but changes in allele frequencies can alter the variance those effects contribute. For diploid, biallelic loci,

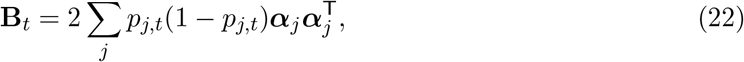

where *p*_*j,t*_ is the allele frequency at locus *j* in generation *t* and ***α***_*j*_ is its fixed allelic-effect vector in the assumed additive model. Directional selection can change **B**_*t*_ through allele frequencies even when every ***α***_*j*_ stays fixed (Walsh and Lynch, 2018). Replacing **B**_*t*_ by a constant **B** is an additional restriction, appropriate to a short-timescale approximation or a formal infinitesimal closure.

Under these assumptions, Equation (21) takes the preselection covariance **G**_*t*_ through the selected-parent state to the next generation’s preselection covariance **G**_*t*+1_. The share is calculated from the slope, curvature and covariance at each generation. It describes that state; it does not determine the covariance update.

Within this exact Gaussian, log-quadratic closure, the covariance update is independent of the directional gradient ***b***_*t*_. A fixed gradient is therefore unnecessary for the covariance fixed point, although a trajectory for *A*_*g*_ requires a specified gradient history. Reproduction can also increase covariance relative to its preselection value even when selection reduces covariance among contributing parents. For example, with *B* = 1, *γ* = 0.3, *ρ* = 0.9, and initial scalar variance *g*_*t*_ = 0.1, the next variance is approximately 0.1874. The recursion therefore allows increasing as well as decreasing covariance trajectories; the isotropic example in Figure 4 begins above its fixed point.

**Figure 4:**
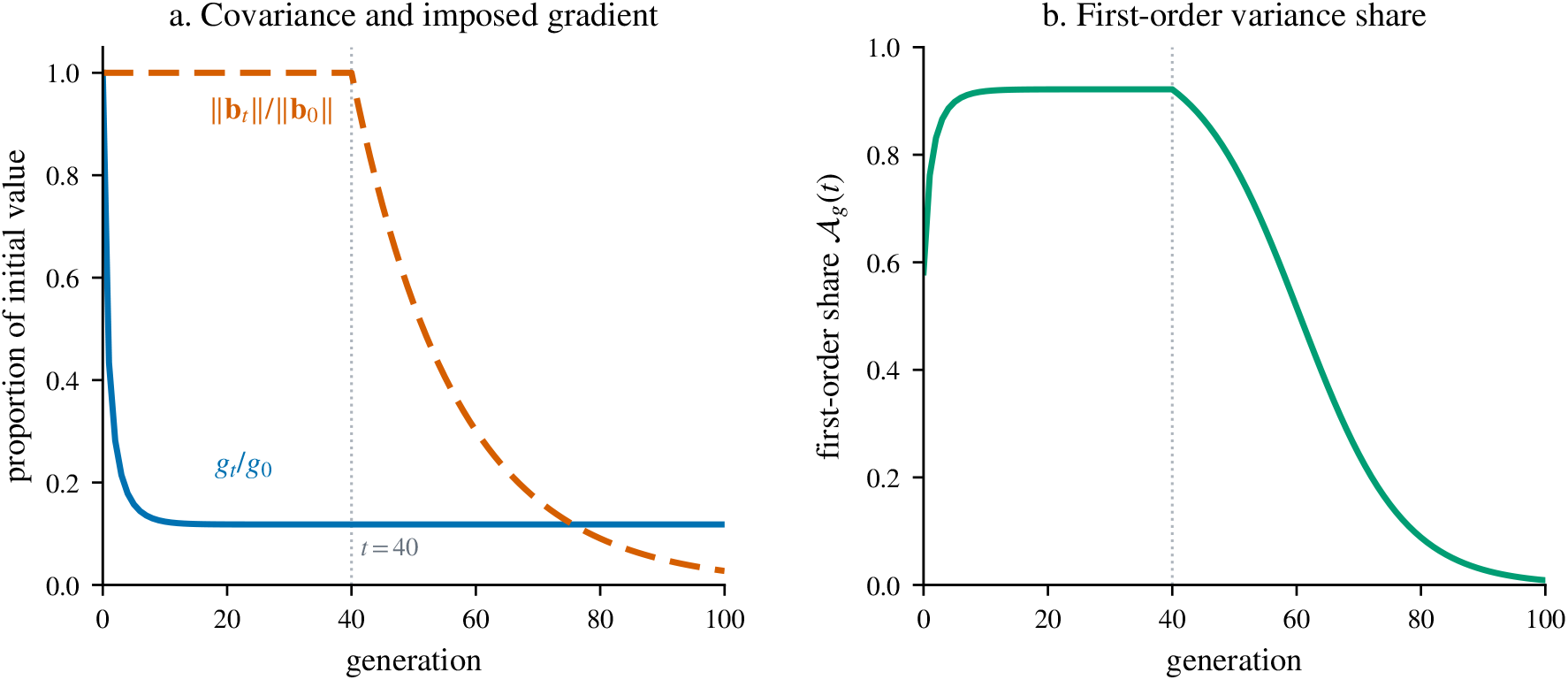
How does the linear share change when covariance falls and the gradient later weakens? The calculation uses Equation (25) with four identical trait dimensions, *g*_0_ = 4, *B* = 1, *γ* = 0.30, *ρ*= 0.90, and initial local log-fitness gradient norm ∥***b*** *∥*= 1. (a) Covariance approaches *g*^*\**^ = 0.472 and is plotted as *g*_*t*_*/g*_0_. The gradient norm is held at its initial value through generation 40 and then declines as exp[ 0.06(*t* 40)] under an imposed history. (b) While the gradient is maintained,*A* _*g*_ rises from 0.581 to 0.922 because *V*_quad_ decreases with 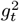whereas *V*_lin_ decreases with *g*_*t*_ (Equation (13)). _*g*_ falls after the imposed gradient declines. The calculation uses a fixed genic baseline and a separately specified gradient history. *Alt text:* Panel a shows normalized covariance falling rapidly to a positive plateau while an imposed gradient remains constant until generation 40 and then declines. Panel b shows the first-order variance share rising during covariance contraction and falling after the gradient begins to relax.

The update allows us to ask whether covariance can persist under the opposing effects of selection and reproduction. Write *F* (**G**) for the right-hand side of Equation (21). For symmetric matrices, **A** ⪯ **C** means that **C** − **A** is positive semidefinite. The map sends the Loewner interval

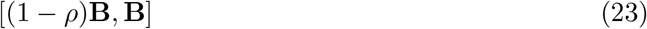

into itself. Brouwer’s fixed-point theorem then gives at least one positive-definite covariance fixed point, and every such fixed point **G**^*\**^ satisfies

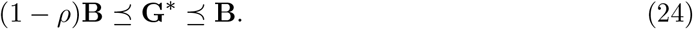

At a fixed point, the covariance reduction caused by selection is balanced by the fraction of the departure from **B** removed during reproduction. The proof establishes existence and the stated bounds. Uniqueness and convergence require a further argument, available for the scalar case below but not established here for a general matrix recursion.

For one trait, or for the isotropic case **G**_*t*_ = *g*_*t*_**I, B** = *B***I**, and **Γ** = *γ***I**, that argument reduces to the scalar update

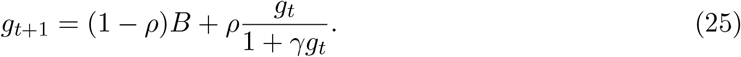

For stabilizing curvature *γ >* 0, its unique positive fixed point is

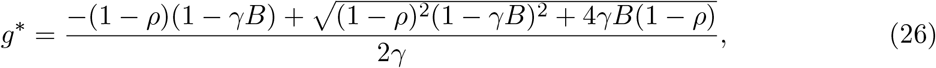

which reduces smoothly to *g*^*\**^= *B* when curvature is absent (*γ* = 0). The absolute derivative of the scalar map is bounded by *ρ <* 1. It follows that every nonnegative starting variance converges to this unique fixed point on [0, *∞* ).

The boundary *ρ* = 1 shows why reproduction’s effect on associations matters. If it retains the entire covariance departure generated by selection, repeated updates give

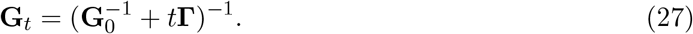

With strictly positive-definite curvature (**Γ ≻**0), covariance approaches zero. If **Γ** is singular, covariance can persist only along its null directions; if **Γ** = **0**, every starting covariance is a fixed point.

An equilibrium for covariance does not fix the fitness differences expressed across it. Those also depend on the local log-fitness gradient. Figure 4 shows the consequence by holding the gradient constant through generation 40 and then imposing a decline. During the constant-gradient period, proportional covariance contraction increases the first-order share. The share falls once the gradient declines. The example therefore follows both the change in standing variation and the separately specified change in the local gradient.

With fixed **B ≻** 0, stabilizing curvature **Γ** ⪰0 and 0 ≤ *ρ <* 1, the model has at least one positive-definite covariance fixed point within the stated bounds. In the scalar or isotropic case, every nonnegative starting variance converges to the unique fixed point. These are covariance predictions under the specified inheritance model, not a joint equilibrium of allele frequencies,population mean and local log-fitness gradient. Applying the variance share to observations is a separate task: the fitness relationship and covariance must be estimated in the population being described.

## 6 Estimating the variance share

An observed population can be assessed without assuming that it follows the covariance recursion. What is needed is its current conditional fitness relationship and breeding-value distribution. Estimate the local slope and curvature of log expected fitness and the breeding-value covariance from the same population, then check the Gaussian and quadratic assumptions used for the variance partition (Table 1).

The observation model must preserve the target in Equation (1). Fit mean reproductive output given breeding value using a distribution appropriate to the observations. Log-link models provide a related approach for phenotypes (Morrissey and Goudie, 2022); logging counts before fitting instead changes the target and excludes zero counts. For a generalized linear mixed model, distinguish model-scale parameters from means on the observed scale (de Villemereuil et al., 2016). The following five steps keep the fitness relationship, population covariance and their uncertainty together:

1. Define absolute fitness and identify the reference population for a single selection episode.
2. Fit the conditional mean to the observed outputs, allowing for uncertainty in each breeding value. Choose a link that keeps the mean in its possible range: positive for a log-linked count mean, or between zero and one for a survival probability.
3. Differentiate the fitted log E[*W* | ***A***] at the population mean to estimate slope ***b*** and curvature **Ĥ** .
4. Estimate covariance 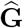for the same population and trait coordinates. Insert the three estimates into Equations (7)–(9).
5. Quantify sampling and breeding-value uncertainty in *V*_lin_, *V*_quad_ and *A*_*g*_. Check whether a quadratic describes the observed range and how the estimates change under other plausible models.

Supplementary File 3 implements these steps using simulated height measurements and surviving-offspring counts. Repeated measurements quantify uncertainty in breeding values, and the fitted expected offspring counts supply slope and curvature. The known generating values let readers check the resulting variance components; the accompanying Shiny app also shows what happens when the quadratic assumption fails. The data and code are in Supplementary File 2.

Report *V*_lin_ and *V*_quad_ with *A*_*g*_. Those components distinguish a high share supported by substantial linear variation from a similar share obtained when both contributions are small.

This is the empirical counterpart of the fixed-surface population comparison.

### 6.1 What existing regressions can supply

An existing regression can supply the share only if its response and predictors represent the required quantities. One special observation model allows a regression on a measured log outcome. Suppose *Y* is the quadratic approximation plus a constant and an error with zero conditional mean:

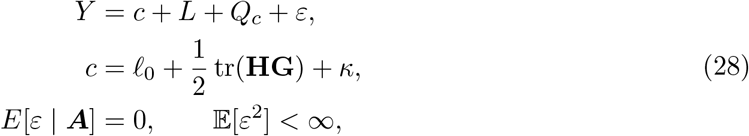

where *κ* does not depend on breeding value. The ratio of population *R*^2^ values in Equation (12) then equals *A*_*g*_. For *Y* = log *W*, this requires E[log *W* ***A***] log E[*W* ***A***] to be constant. For example, let *W* = *W*_*A*_(***A***)*U*, with positive *U* independent of breeding value, E[*U* ] = 1 and E[(log *U* )^2^] *< ∞*. Then *κ* = E[log *U* ] can differ from zero. The observation model must support this assumption; it provides neither a way to log zero counts nor a justification for ordinary least squares on count data.

The predictors need the same care. Estimated breeding values can have a different distribution from the unobserved values they predict (Hadfield et al., 2010). Suppose slope and curvature are known on the true breeding-value scale, but the analysis replaces **G** by the variance of predicted values, Var(***Â*** ) = *r***G**, with scalar reliability 0 *<r <* 1. The naive share is then

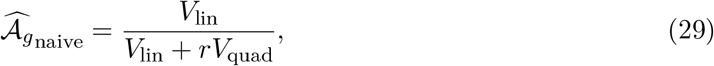

which exceeds the target when both variance components are positive. Shrinkage has made the population appear less variable, reproducing the same scaling effect that raised the share in the fixed-surface comparison. This direction of error is specific to substituting the covariance while keeping slope and curvature on the true scale. It can differ when those coefficients are also fitted to predicted values or reliabilities vary among traits. Their joint uncertainty must therefore be included in interpreting the share.

Some published phenotypic selection analyses provide another route, through the moments of selected parents. This requires compatible first- and second-order coefficients and phenotypic moments, not just a reported fit statistic. Under standard linear–quadratic phenotypic regression and its multivariate generalizations (Lande and Arnold, 1983; Phillips and Arnold, 1989; Blows and Brooks, 2003), let the centered phenotype be ***x***, relative fitness be *w* = *W/*E[*W* ], and the population least-squares cofficients be ***β*** and ***γ***, with the convention

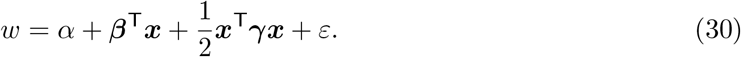

The joint-Gaussian, log-quadratic closure conditions in Appendix S5 allow those moments to be matched in breeding-value coordinates. Define **K** = ***γ*** *−β β* ^⊤^. The conversion gives

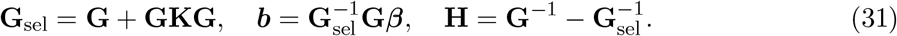

These quantities allow *A*_*g*_, *V*_lin_ and *V*_quad_ to be reported alongside the selection regression if its coefficients use compatible unstandardized coordinates and the same factor-of-two convention (Stinchcombe et al., 2008). If the selected-moment identities hold but log expected fitness is not exactly quadratic, Equation (31) gives a moment-matched quadratic approximation. Joint Gaussianity, mediation of selection, coordinate compatibility and the stated matrix conditions remain necessary for this conversion.

Both routes return a share for a specified outcome and population. Keeping that target explicit, and accounting for measurement error in breeding values, allows the estimated linear description to be interpreted alongside the population’s actual variation.

### 6.2 Code and teaching resources

Supplementary File 2 supplies the figure-generation scripts, mathematical checks, SLiM model with its seeds and archived diagnostic tables, and the worked estimation example with simulated observations and a Shiny app. Its reproduction guide identifies the inputs and commands for each result. Supplementary File 1 explains the calculations, and Appendix S7 illustrates population comparisons for one and two traits. Supplementary File 3 works through estimation. The short companion is additional repository material, separate from these supplementary files.

## 7 Discussion

The apparent linearity of fitness depends on the breeding values present as well as on the relationship between breeding value and reproduction. The fixed-surface comparison isolates that population contribution. For Gaussian breeding values on a quadratic log-fitness surface, multiplying covariance by *s* multiplies the linear variance by *s* and the quadratic variance by *s*^2^. With 0 *<s <* 1 and both components positive, the linear share rises even as the linear variance falls. This extends variance comparisons in selection regressions (Lande and Arnold, 1983; Moorad and Wade, 2013) to log expected fitness conditional on breeding value.

A difference in the share between populations can therefore have more than one explanation. In Figure 3, the surface and mean are fixed, so the larger share comes from the narrower breeding-value distribution. In field comparisons, the conditional fitness relationship may also change. Even on the same phenotype–fitness surface, individuals with a given breeding value can experience different environments or differ in other traits. Estimating the conditional mean and breeding-value distribution in each population, and reporting *V*_lin_ and *V*_quad_ with *A* _*g*_, distinguishes a larger linear fraction from more linear variation. Changes in the mean or covariance orientation require their own assessment; proportional scaling alone does not predict their direction.

Such a comparison also needs the Gaussian and quadratic approximations to describe each population adequately. Higher-order terms can matter, for example near an inflection point. Strong selection can produce departures from Gaussianity (Turelli and Barton, 1994), and nonzero third central moments can make the linear and quadratic terms covary (Lande and Arnold, 1983). In either case, the linear projection under the observed distribution can be estimated and compared with the Gaussian calculation. Predictions across generations add separate assumptions about constant stabilizing curvature, fixed genic covariance and a common retention parameter.

The comparison with selection asks a different question of the same individuals. In the one-trait example, the line explains 97% of variation in log expected fitness but overestimates the selected-parent shift by 50% when curvature is omitted. Expected-output weighting changes the importance of differences that receive little weight in the variance partition. Equation (20) assesses the resulting displacement error using absolute curvature in population-scaled coordinates. For data, the same distinction starts with fitting mean reproductive counts at each breeding value, retaining zero counts and allowing uncertainty in the breeding values. Appendix S5 specifies when an existing selection regression can supply the parameters.

Calling the population–fitness relationship an additive channel does not identify the underlying gene action. Interacting alleles can contribute to additive, dominance and interaction variances in proportions that depend on genotype frequencies (Cheverud and Routman, 1995; Hansen, 2013).

The share describes fitness across the breeding values in a given population. If average effects change as selection shifts the mean, the fixed-effects inheritance model no longer describes that process (Hansen et al., 2006). Directional positive epistasis, for example, can increase additive genetic variance as the mean changes (Carter et al., 2005).

The inheritance model makes those additional assumptions explicit. Selection generates associations between loci, reproduction reduces them, and the genic baseline remains fixed (Bulmer, 1971; Barton and Turelli, 1991). Under this closure, the scalar covariance recursion converges to a unique equilibrium; for several traits we establish fixed-point existence and matrix bounds. Figure 4 follows the share along one covariance trajectory while specifying the gradient history separately. The model provides a conditional account of covariance across generations, excluding changes in the genic baseline caused by mutation or changing allele frequencies.

Following a real population through time need not assume fixed average effects. Breeding values, covariance and the conditional fitness surface could instead be estimated in successive populations, checking the Gaussian and quadratic approximations each time. This would allow average effects to change with allele frequencies (Hansen et al., 2006; Carter et al., 2005). The resulting comparisons would describe when the share changed and whether its absolute components also changed. Explaining the cause of those changes would require additional evidence.

A fitness surface describes how expected reproduction differs among breeding values; a population determines how often each difference occurs. The first-order share and its two variance components quantify their combined effect on a linear description. The same information also bounds the error in the selected-parent mean shift when curvature is omitted. The population and its fitness relationship therefore support two assessments: how much variation a line describes, and how accurately it predicts the shift in mean breeding value among selected parents.

## Data availability

Supplementary File 2 is the archived code and data record for this version. It contains the figure-generation scripts, machine-readable numerical results, mathematical verification code, the SLiM model and diagnostic-analysis script, and the 20 archived replicate-level diagnostic tables used in Appendix S4. Supplementary File 3 works through the estimation procedure; its data, analysis and Shiny app are in Supplementary File 2. The source archive contains the manuscript and Supplementary Files 1 and 3. The reported simulation summary can be regenerated from the diagnostic tables without rerunning SLiM; the supplied model and command also permit a fresh simulation run. The project repository is https://github.com/dortizbarrientos/additive_channels. The code and accompanying documentation are distributed under the MIT licence. No new empirical genotype or phenotype data were generated.

## Acknowledgments

We thank Guillaume Martin and all reviewers for their insightful, rigorous feedback on our work; it substantially improved the quality of this manuscript. We also thank colleagues in the ARC Centre of Excellence for Plant Success in Nature and Agriculture for discussions and Bruce Walsh for discussions concerning the origin and maintenance of quantitative genetic variation.

## 8 Artificial-intelligence assistance

OpenAI Codex and ChatGPT (OpenAI, San Francisco, CA) and Claude (Anthropic, San Francisco, CA; Claude Opus models) assisted during revision by checking mathematical derivations and numerical calculations, and co-piloting Python code for the figures. The authors reviewed the resulting text, calculations and code, and take responsibility for all content in the manuscript.

## Appendix S1: Gaussian moments and population projections

Let ***X*** *∼N*(**0, G**), let *L* = ***b*** ^⊤^***X***, and let *Q*_*c*_ be defined by Equation (6). The linear-form identity gives

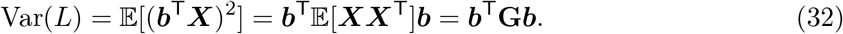

For a centered Gaussian, the covariance of products follows from the Gaussian fourth-moment identity (Isserlis, 1918):

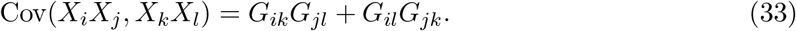

Substitution into the quadratic form gives

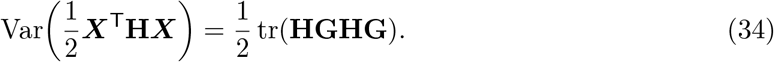

All centered third moments vanish, so Cov(*L, Q*_*c*_) = 0. These identities yield Equations (7) and (8).

For the population affine projection, write *c* + ***d*** ^⊤^***X*** and impose the normal equations. The intercept is the mean of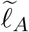,

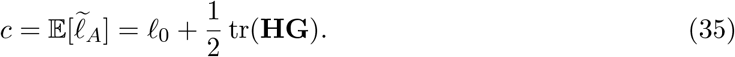

The slope satisfies

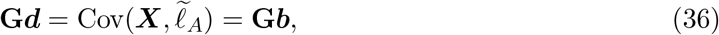

because Cov(***X***, *Q*_*c*_) = **0**. Thus ***d*** = ***b*** on the support of **G**. Its explained variance is *V*_lin_, whereas the variance of the full second-order target is *V*_lin_ + *V*_quad_. This proves Equation (11). Adding a response residual with conditional mean zero and finite variance adds the same residual variance to the denominator of the linear and quadratic *R*^2^ values and proves Equation (12).

An invertible coordinate change ***X*** ^*’*^= **D*X*** gives

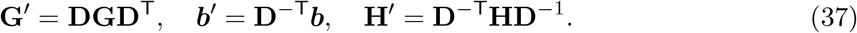

Direct substitution leaves ***b***^T^**G*b*** unchanged. The product **HG** changes by similarity, so cyclicity of the trace leaves tr[(**HG**)^2^] unchanged. Both variance terms and *A* _*g*_ are therefore coordinate invariant.

Finally, diagonalizing 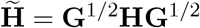 gives

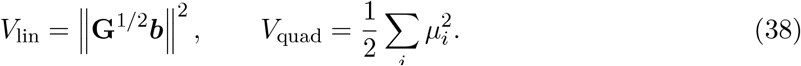

This expression proves that no alignment between the whitened gradient and the eigenvectors of 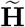 enters *A*_*g*_. The separate Rayleigh-quotient result follows from maximizing ***b*** ^*⊤*^**G*b*** over gradients of fixed Euclidean norm.

If *V*_lin_ = 0 *< V*_quad_, the affine projection is constant and its *R*^2^ can be recorded as zero, but Corr(*L, L* + *Q*_*c*_) is undefined because *L* has zero variance. If *V*_lin_ = *V*_quad_ = 0, both the ratio and the correlation are undefined. For singular **G**, the same derivations hold after restricting coordinates to its positive-eigenvalue support.

## Appendix S2: Gaussian selection and the displacement bound

Multiplying the preselection density of ***X*** *∼ N* (**0, G**) by Equation (15) gives an exponent

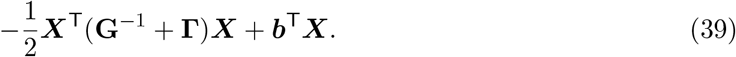

When **G**^*−*1^ + Г ≻ 0, completing the square gives covariance (**G**^*−*1^ + Г)^*−*1^ and mean (**G**^*−*1^ + Г)^*−*1^***b***. This proves Equations (16) and (17). Notice that the covariance is independent of ***b*** only within this exact Gaussian, log-quadratic closure; the selected mean is not.

For the displacement bound, assume **G** ≻0 and **G**^1*/*2^***b*** ≠ **0**. If **G** is singular, all inverses and norms below are defined on its support, where the gradient must have a nonzero component. Define ***c*** = **G**^1*/*2^***b*** and **T** = **G**^1*/*2^Г **G**^1*/*2^. In the norm 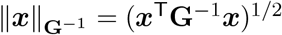,

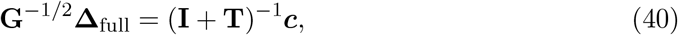

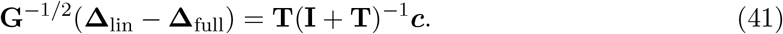

Therefore

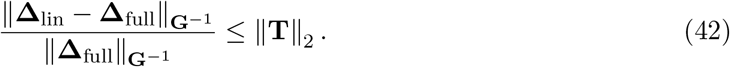

The spectral norm is no larger than the Frobenius norm, and

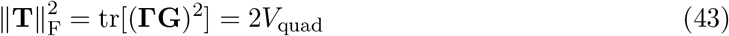

because **H** = − Г. This proves Equation (20). If ***c*** is an eigenvector of **T** with eigenvalue *τ*_*k*_, the numerator and denominator are scalar multiples of that eigenvector and the exact ratio is |*τ*_*k*_ |. The one-trait construction in Figure 1 also illustrates why phenotype-space and breeding-value-space curvature differ. Let

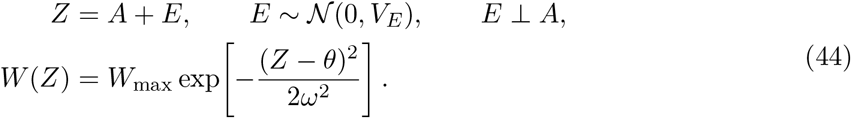

Following the conditional averaging used by Lande (1979), integration over *E* gives

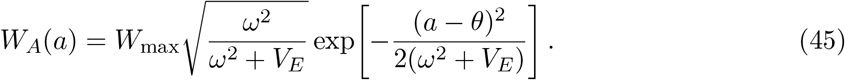

Thus the breeding-value-space curvature is *H* = − (*ω*^2^ + *V*_*E*_)^*−*1^ rather than the phenotype-space value *− ω* ^*−*2^. Environmental variation has already been averaged out before the logarithm is taken.

## Appendix S3: Fixed-baseline covariance recursion

Equation (21) can be written **G**_*t*+1_ = *F* (**G**_*t*_), where

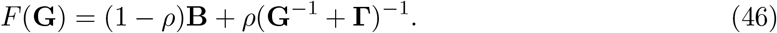

Assume **B** *≻* 0, Г *⪰* 0, and 0 ≤ *ρ <* 1. For every positive-definite **G**,

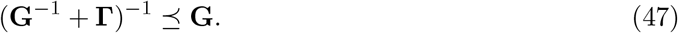

If (1 − *ρ*)**B** *⪯* **G** *⪯* **B**, then Equation (47) gives

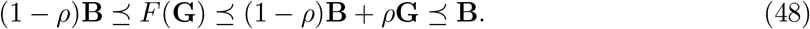

The map is continuous and therefore maps a compact convex order interval into itself. Brouwer’s theorem gives at least one fixed point in that interval.

Every positive-definite fixed point obeys the same bounds, whether or not a trajectory is initialized in the interval. The lower bound follows directly from the nonnegative second term of *F* . For the upper bound, let **G**^*\**^ = *F* (**G**^*\**^) and use Equation (47):

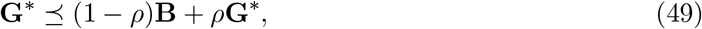

which implies **G**^*\**^ *⪯* **B** because 1*− ρ >* 0. This proves Equation (24). The argument does not prove uniqueness or convergence of the matrix iteration.

In one dimension the fixed-point equation is

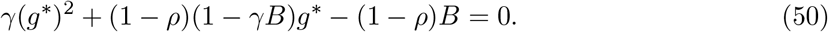

The positive root is Equation (26). For any two nonnegative values *g* and *h*, direct subtraction gives

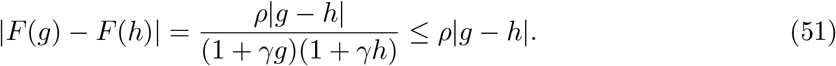

Thus after each update the distance between two scalar trajectories is at most *ρ* times its previous value. The fixed point is consequently unique and globally attracting in the scalar case.

When *ρ* = 1, inversion of the update gives 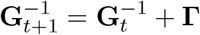. Induction yields Equation (27).

This boundary lacks the positive restoration term that makes the order interval compact. Strictly positive curvature removes covariance in every direction; singular curvature can leave covariance only within its null space.

The model treats *ρ* as a scalar summary of linkage-disequilibrium retention. It does not assert a universal formula from recombination rate, selfing, or inbreeding to *ρ*. Such a formula would depend on locus pairs, their effects, signs of disequilibria, recombination architecture, and the population’s mating system (Bulmer, 1974; Barton and Turelli, 1991).

## Appendix S4: Scope of the forward-time simulation

To examine the Gaussian approximation in finite populations, we used a forward-time individual-based simulation in SLiM 5.2 (Haller et al., 2026). It used 20 replicates of a diploid population of *N* = 2,000 individuals and sequential seeds 42–61. Two hundred quantitative-trait loci, 50 per trait, contributed to four breeding values. The trait-specific standard deviations of allelic effects were (0.346, 0.283, 0.200, 0.141). Effects summed linearly within traits, so inherited allelic content mapped additively to breeding value. Gaussian stabilizing selection had diagonal strengths (0.10, 0.05, 0.04, 0.03). The optimum for trait one moved by 0.3 per generation for 40 generations to (12, 0, 0, 0) and then remained fixed; each replicate ran for 200 generations. No residual environmental deviation was added, so the simulated phenotype used in the fitness calculation equaled the additive trait value.

At each generation, the analysis calculated **G, *b***, and **H** and compared analytic *A*_*g*_ with the squared sample correlation between *L* = ***b***^⊤^***X*** and *L* + *Q*_*c*_. Across the 20 replicates and 200 generations, the median absolute difference was 0.0157. A fresh run with the same seeds reproduced the archived diagnostics to numerical precision. Breeding-value distributions in these finite populations need not be exactly Gaussian, so this comparison assesses the approximation in the simulated design.

The agreement supports using the current-population calculation as an approximation in this multilocus design. It does not test selected-parent displacement or the fixed-**B** recursion. Drift and allele-frequency change alter genic covariance through Equation (22), despite the fixed allelic effects.

The model, seeds, driver scripts, and generation-level summaries are available from the repository cited in the Data availability statement.

## Appendix S5: Moment conversion from phenotype-space selection regressions

This appendix gives a restricted route from a compatible Lande–Arnold regression (Lande and Arnold, 1983) to the breeding-value-space quantities used here. The normalization of relative fitness gives E[*w*] = 1; it does not imply that fitted relative fitness at the mean phenotype equals 1.

Let centered phenotype ***x*** have covariance **P** and let the population least-squares regression be Equation (30), with the residual orthogonal to the constant, linear, and quadratic regressors. The convention places a factor 1*/*2 before the whole quadratic form; software coefficients on squared terms must be converted to this convention (Stinchcombe et al., 2008). Gaussian moments give

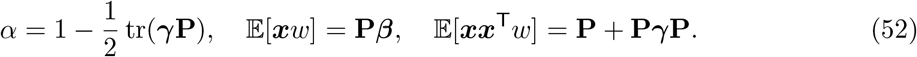

Assume that breeding value ***A*** and phenotype ***Z*** are jointly Gaussian in the same coordinates, with Cov(***A, Z***) = **G**, and that selection is mediated by the measured phenotypes. Gaussian conditioning then gives the selected breeding-value mean

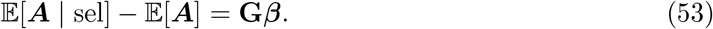

Subtracting the outer product of this mean from the selected second moment gives

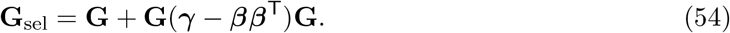

Under an exact Gaussian, log-quadratic model in breeding-value space, Equations (16) and (17) must reproduce these selected moments. Solving for the local parameters yields Equation (31):

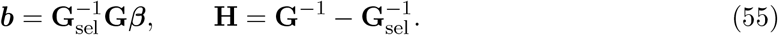

The conversion requires unstandardized coefficients in compatible trait coordinates, the same reference population and selection episode, joint Gaussianity, selection mediated by the measured traits or an equivalent conditional-moment independence, and positive definiteness of **G**_sel_ on the support of **G**. Uncertainty in ***β, γ*, G**, and every transformation must be propagated. If the selected-moment identities remain valid but exact log-quadraticity fails, the equations define a moment-matched quadratic approximation. If joint Gaussianity, selection mediation, coordinate compatibility, or the matrix conditions fail, this conversion is not justified. In neither case do the equations recover point derivatives of an arbitrary log E[*W* | ***A***].

For comparison, direct differentiation of the fitted raw-scale polynomial at ***x*** = **0** would give

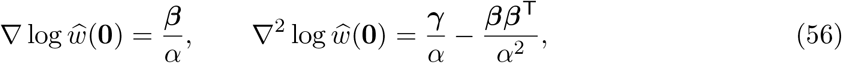

provided *α >* 0. These are derivatives of the fitted phenotype-space projection, not generally the breeding-value-space target of Equation (1).

## Appendix S6: Anticipated questions

This appendix answers recurring questions about the scope, interpretation, and use of the first-order variance share. Table 1 lists the assumptions behind each answer.

### Q1: What does the first-order variance share measure?

The share is

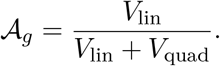

It is the fraction of variation in the quadratic approximation to log expected fitness contributed by the linear term. For example, *A*_*g*_ = 0.8 means that the linear term accounts for 80% and curvature for 20%. This describes the breeding values in the current population. Realized fitness includes further variation, so its variance is not the denominator. Report *V*_lin_ and *V*_quad_ alongside *A*_*g*_: a fraction alone cannot distinguish two large components from two small ones. Equations (7)–(9) define the quantities.

### Q2: Which parts are established, and what does this paper add?

Breeding values summarize trait variation through average allelic effects (Fisher, 1918, 1941). Averaging fitness over environmental deviations (Lande, 1979), weighting by population frequencies and partitioning linear and quadratic fitness variation (Lande and Arnold, 1983; Moorad and Wade, 2013), log-link fitness models (Morrissey and Goudie, 2022) and selection-generated linkage disequilibrium (Bulmer, 1971) are established ideas. Here we calculate the variance share for log expected fitness conditional on breeding value, derive its proportional-covariance scaling, and separate its interpretation from selected-parent prediction and an inheritance model. Showing a population distribution alongside a fitness surface is a way to explain the weighting, not a claim that classical theory omitted the population.

### Q3: Why average fitness first and then take its logarithm?

Averaging reproductive output at each breeding value gives its expected contribution to the selected group (Equation (2)). The logarithm is then used to describe proportional differences: a doubling of expected output gives the same log difference from one offspring to two as from ten to twenty. Selection still weights individuals by expected output itself. A variance partition on the log scale is a description of proportional differences and need not equal a partition on the original fitness scale. The order also matters: if two individuals with the same breeding value leave zero and two offspring, their mean is one and its logarithm is zero; the average of their logged outputs is undefined (Equation (4)).

### Q4: Does an additive channel imply additive gene action or require a numerical cutoff?

No. The term describes a population whose linear contribution to variation in log expected fitness is large relative to the curved contribution. It says nothing about how alleles combine to produce a trait. We impose no universal cutoff on this continuous share; an application would need to justify any decision threshold. The separate inheritance model assumes fixed allelic effects and a linear, time-invariant map from allelic content to breeding value. A large variance share does not establish those assumptions.

### Q5: Which results require Gaussian breeding values?

The conditional selection weight applies to any breeding-value distribution. Gaussian breeding values give the moments used for the variance formulas and regression identity. Symmetry alone is enough to make the centered linear and quadratic terms uncorrelated, but does not in general give the Gaussian fourth-moment formula for *V*_quad_. With a non-Gaussian distribution, that formula may therefore differ from the actual quadratic variance, and asymmetry can also correlate the two terms. One can calculate the regression projection under the observed distribution or label the Gaussian expression as an approximation. The selected-parent formulas additionally require an exactly log-quadratic selection weight and an integrable Gaussian update.

### Q6: How can the first-order variance share be estimated?

Fit a model for expected reproductive output conditional on breeding value, using a response distribution appropriate to the data. Take the fitted mean’s logarithm, estimate its slope and curvature at the current population mean, and combine these with breeding-value covariance in Equations (7)–(9) (Morrissey and Goudie, 2022; de Villemereuil et al., 2016). Account for uncertainty in breeding values and in all fitted quantities (Hadfield et al., 2010). Regressing logged individual fitness estimates a different conditional mean; the ratio of linear and quadratic *R*^2^ values recovers *A* _*g*_ only under the constant-gap model in Equation (28). Equation (29) illustrates a particular bias from treating prediction shrinkage as biological variation. Appendix S5 gives the conditions for using phenotype-space selection regressions.

### Q7: What happens when the gradient or both variance terms vanish?

If *V*_lin_ = 0 *< V*_quad_, then *A* _*g*_ = 0: curvature contributes variation while the linear term contributes none. The correlation form of Equation (11) is then undefined because one variable has zero variance. If both components vanish, *A* _*g*_ itself is undefined. A zero gradient can occur at a stationary point of expected fitness, but it does not explain the curvature or identify gene action.

### Q8: Does a high first-order variance share guarantee correct rankings or an accurate response? No

A regression *R*^2^ describes variance explained, rather than agreement in rank. Nor does it measure the accuracy of a selected-parent shift. In Equation (18), the share is about 0.97, yet omitting curvature overestimates that shift by 50%. Equation (20) instead bounds the displacement error using absolute covariance-weighted curvature. Predicting an offspring response requires the further step of specifying inheritance.

### Q9: How do covariance, dimensionality, and orientation enter the calculation?

The calculation uses the combinations of traits that vary in the population. Rescaling them to remove covariance, a transformation called whitening, gives 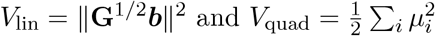, where the *μ*_*i*_ are eigenvalues of **G**^1*/*2^ **HG**^1*/*2^. The number of named traits alone therefore does not determine the share. The relation of the gradient to covariance affects *V*_lin_, and that of curvature to covariance affects *V*_quad_. Once the whitened gradient norm and curvature eigenvalues are fixed, rotating the gradient relative to the curvature eigenvectors leaves *A*_*g*_ unchanged. Although **HG** need not be symmetric, the quadratic variance is nonnegative because

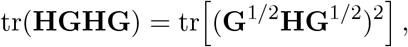

a sum of squared real eigenvalues.

### Q10: What is held fixed across generations, and what do *ρ* and the fixed point mean?

The recursion fixes allelic effects, the genic baseline **B**, stabilizing curvature Г and the retention fraction *ρ*. Fixed effects keep breeding-value coordinates comparable across generations. They do not keep **B** fixed, because allele-frequency changes can alter genic covariance. Holding that baseline fixed is an additional approximation. The parameter *ρ* gives the fraction of the postselection departure from **B** retained after reproduction; one scalar need not represent all genomic directions equally (Bulmer, 1974; Barton and Turelli, 1991). For 0 ≤ *ρ <* 1, the matrix result establishes fixed-point existence and bounds. Uniqueness and convergence are proved only for the scalar case. At *ρ* = 1, covariance collapses when Г ≻ 0, while directions with zero curvature can retain variation. These are covariance results under fixed parameters; they do not establish a joint equilibrium of allele frequencies, means and gradients.

### Q11: If the share predicts neither response nor rankings, what is it for?

It helps compare the quality of a linear description across populations. In Figure 3, the share changes from 0.604 to 0.933 when covariance contracts proportionally on the same local fitness surface. The improved linear description arises from the population change. At fixed slope and curvature, multiplying covariance by *s>* 0 multiplies *V*_lin_ by *s* and *V*_quad_ by *s*^2^. For baseline components *L*_0_ and *Q*_0_, the resulting share is *L*_0_*/*(*L*_0_ + *sQ*_0_) and the omitted fraction is *sQ*_0_*/*(*L*_0_ + *sQ*_0_). The latter does not scale quadratically. This comparison requires proportional covariance change; changes in covariance shape can give a different result. Report the absolute components with the share, and assess predictions of selection separately.

## Appendix S7: A visual explanation of the additive channel

How can a population have a nearly linear fitness description on a curved fitness surface? Figure A1 changes the population while keeping the curve fixed, separating expected reproduction from breeding-value frequency. Figure A2 asks whether the direction of genetic variation matters when two traits vary together. Both examples use specified mathematical populations.

### Read the curve and the population separately

Suppose *a* is a breeding value for plant height, expressed in arbitrary units. Plants with the same breeding value can differ in reproductive output. At each value, the curve gives the logarithm of their mean output. The density below the curve shows how often each breeding value occurs. A fitness value and a frequency therefore answer different questions: how much an individual is expected to reproduce, and how many individuals have that breeding value.

To measure fitness variation, evaluate the curve at the breeding value of each individual. Each individual contributes one squared deviation from mean log expected fitness. Common breeding values receive more weight because more individuals carry them. This frequency weighting is the precise sense in which the population determines which parts of the fitness relationship contribute to the description. It requires no change in the fitness curve itself.

### Compare the curve with a population-fitted line

The dashed line uses those same frequency weights to minimize mean squared differences from the curve across the population. Its predictions vary when breeding value varies; the remaining vertical differences are residuals. In these Gaussian, quadratic examples, variation in the fitted line and variation in the residuals add to the total variance. The linear share is the first component divided by that total. A population and fitness relationship with a large share constitute an additive channel.

The word *additive* refers here to the linear representation: changing breeding value by one unit changes the prediction by the same amount at every value. With several traits, the prediction adds one such contribution per trait. This describes fitness across breeding values in the present population. It does not identify interactions among alleles or specify a route that the population must follow through trait space.

### Change the population while holding the curve fixed

For Figure A1, use

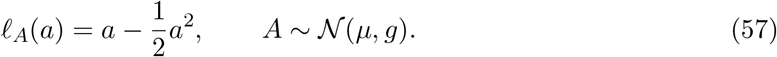

The maximum occurs at *a* = 1. The local slope at the population mean is 1 *− μ*, and curvature is *−*1. Equations (7) and (8) reduce to

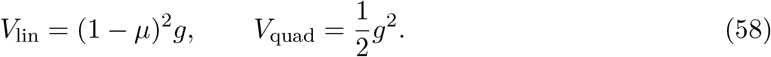

Population A has mean zero and variance one. Its linear and quadratic variances are 1 and 0.5, so the line accounts for 1*/*1.5 = 66.7% of the total. Population B has the same mean but one tenth of the variance. Its components are 0.1 and 0.005, giving 0.1*/*0.105 = 95.2%. Curvature has not changed; its contribution to variance has fallen faster than the linear contribution. B therefore has a stronger additive representation, even though its absolute linear variance is smaller.

Does the narrow range by itself explain B’s high share? Population C has B’s variance but is centered at the peak. The slope there is zero. Plants on either side of the mean have lower expected fitness, so a straight line cannot describe that symmetric decrease. The fitted line is horizontal, its variance is zero, and all of the remaining variance, 0.005, is quadratic. A narrow distribution alone is therefore insufficient for a large linear share. The size of fitness variation and the fraction represented by a line are separate quantities.

To calculate the dashed line itself, include its intercept:

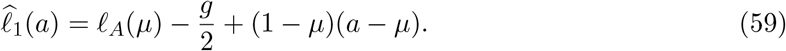

It has the slope of the tangent at the mean but a different height. The constant shift accounts for the average effect of curvature. The residual is −(*a −μ*)^2^ −*g /*2. The bars in Figure A1 compare the variances of these two varying terms, rather than the visual distance between the curve and its tangent.

### Extend the picture to two traits

With two traits, does it matter which combinations vary most? In Figure A2, each point gives an individual’s pair of breeding values, and the contours give log expected output. The ellipse contains 90% of a Gaussian population. The mean, both principal variances and the fitness surface stay fixed across the maps; only the direction of population variation changes. This changes the population on fixed trait axes, rather than relabeling coordinates.

When most variation lies in the direction towards the peak, differences in breeding value produce substantial linear differences in log expected fitness. Turning that variation across the slope reduces those linear differences. Here the surface has the same curvature in every direction, so the quadratic variance remains fixed. The shares fall from 0.911 to 0.839 to 0.203. At 90° the share remains positive because the population still has some width towards the peak. The side view follows only the long axis: its horizontal dashed section at 90 ° does not mean that the complete fitted plane is horizontal.

For reproducibility, these coordinates point towards and across the peak used in Figure 3.

Relative to the value at the mean,

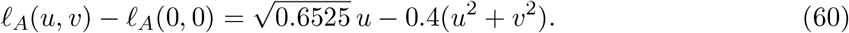

The principal variances are 0.20 and 0.005. If the long axis makes an angle *Ø* with the direction towards the peak, then

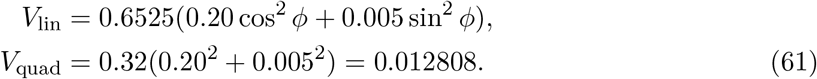

Thus the changing shares in this example arise entirely from changing linear variance.

### Does a close linear description also predict selection?

The figures show how the population distribution determines a linear description of fitness. Selection asks how much each part of that population contributes to reproduction. Multiply each breeding-value frequency by expected reproductive output and normalize (Equation (2)). These weights explain why a close linear description can still predict the selected-parent mean inaccurately. The main text’s one-trait example gives a 97% linear share but a 50% overestimate of selected-parent displacement. Equation (20) provides a separate check on that error by combining curvature with population covariance. To extend the calculation to offspring, specify what parents transmit; the inheritance model in the main text uses this additional step to describe covariance in the next generation.

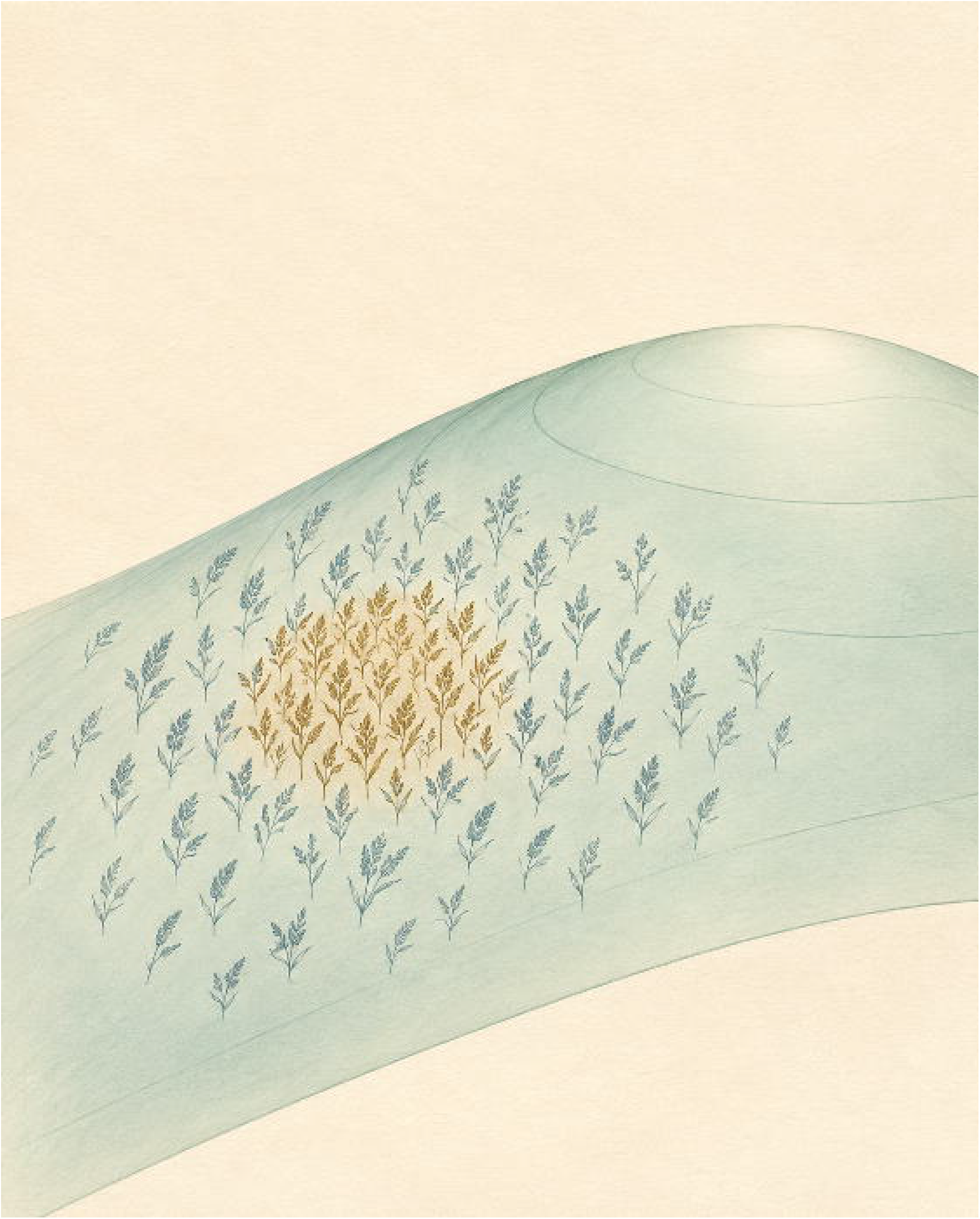

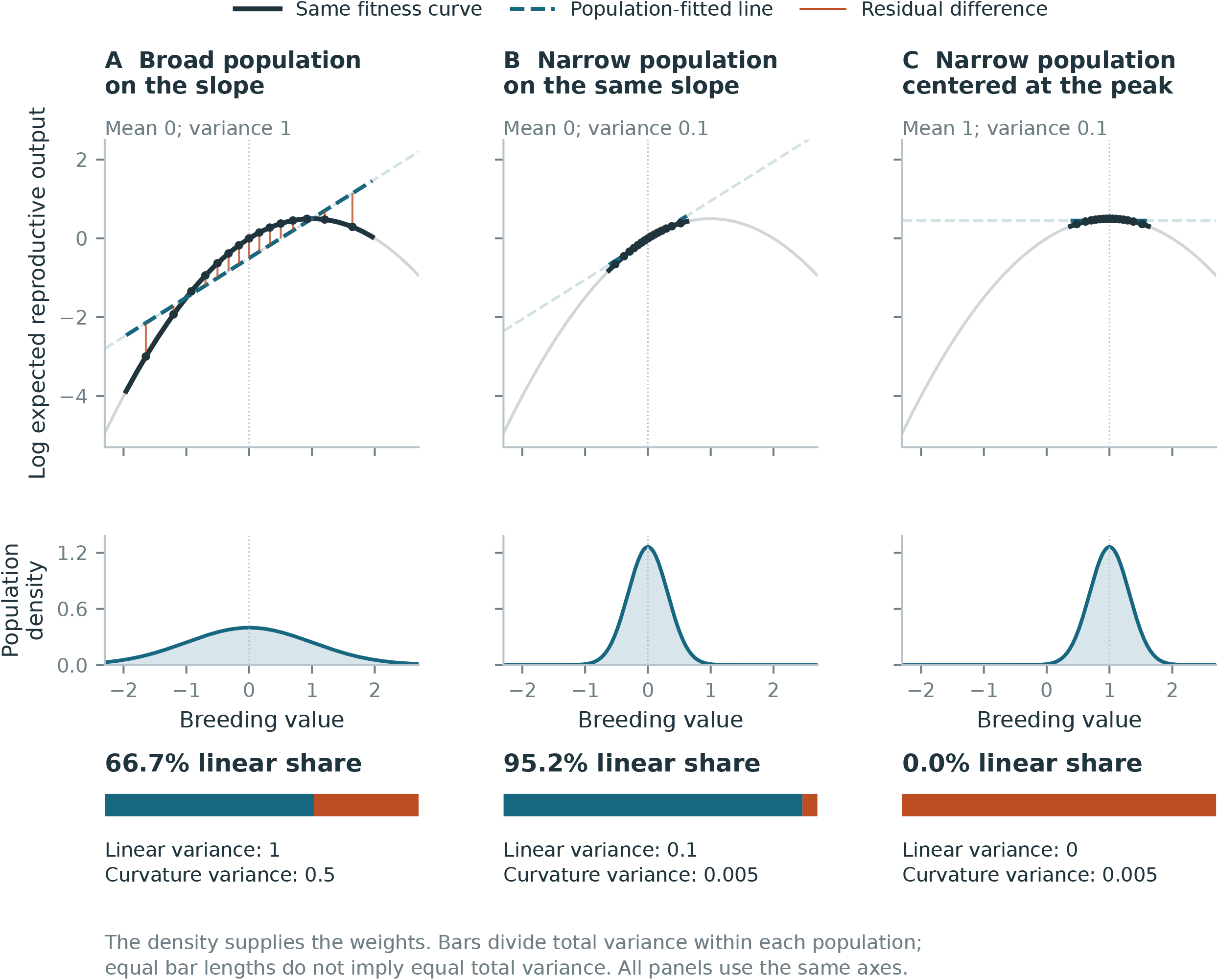

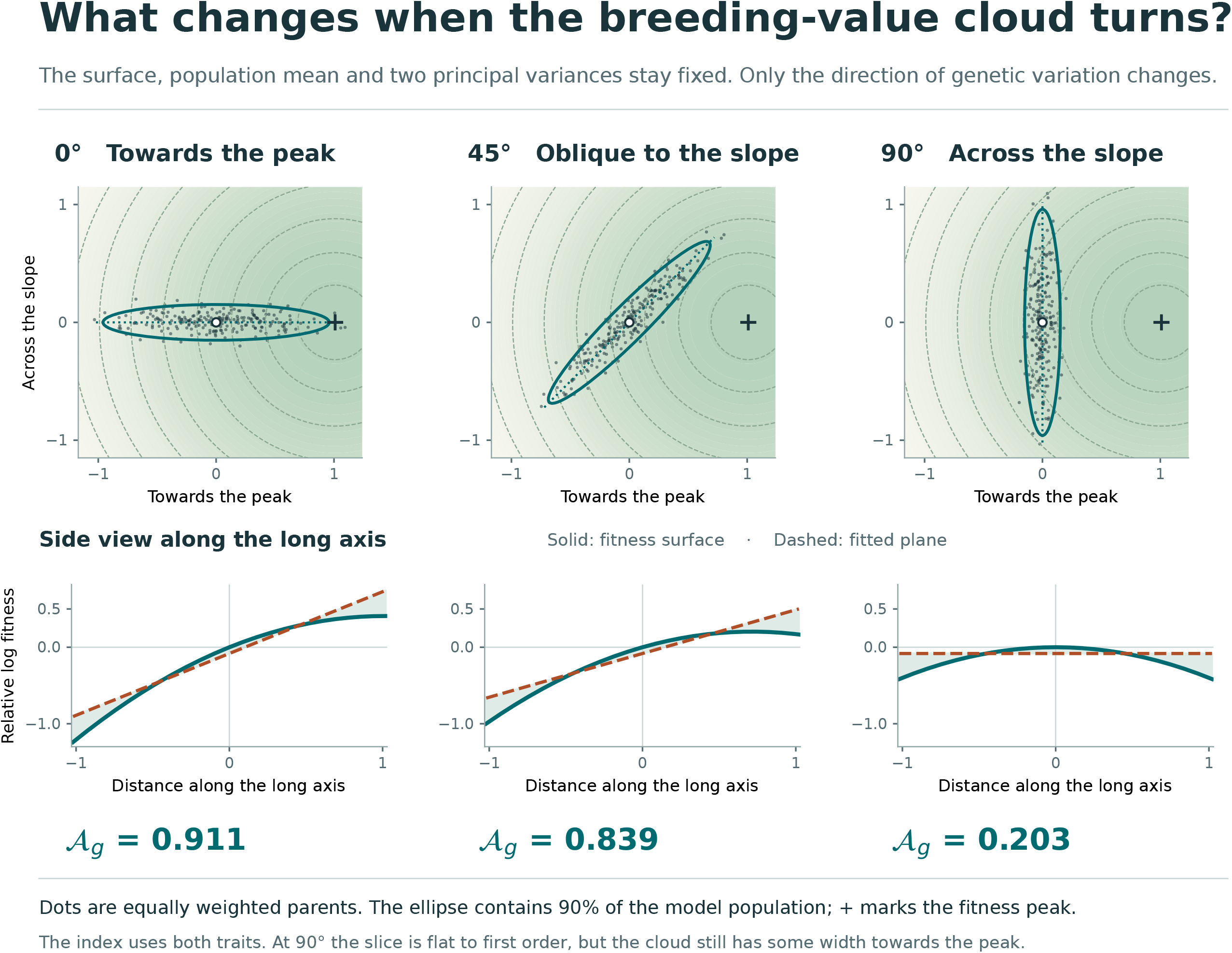

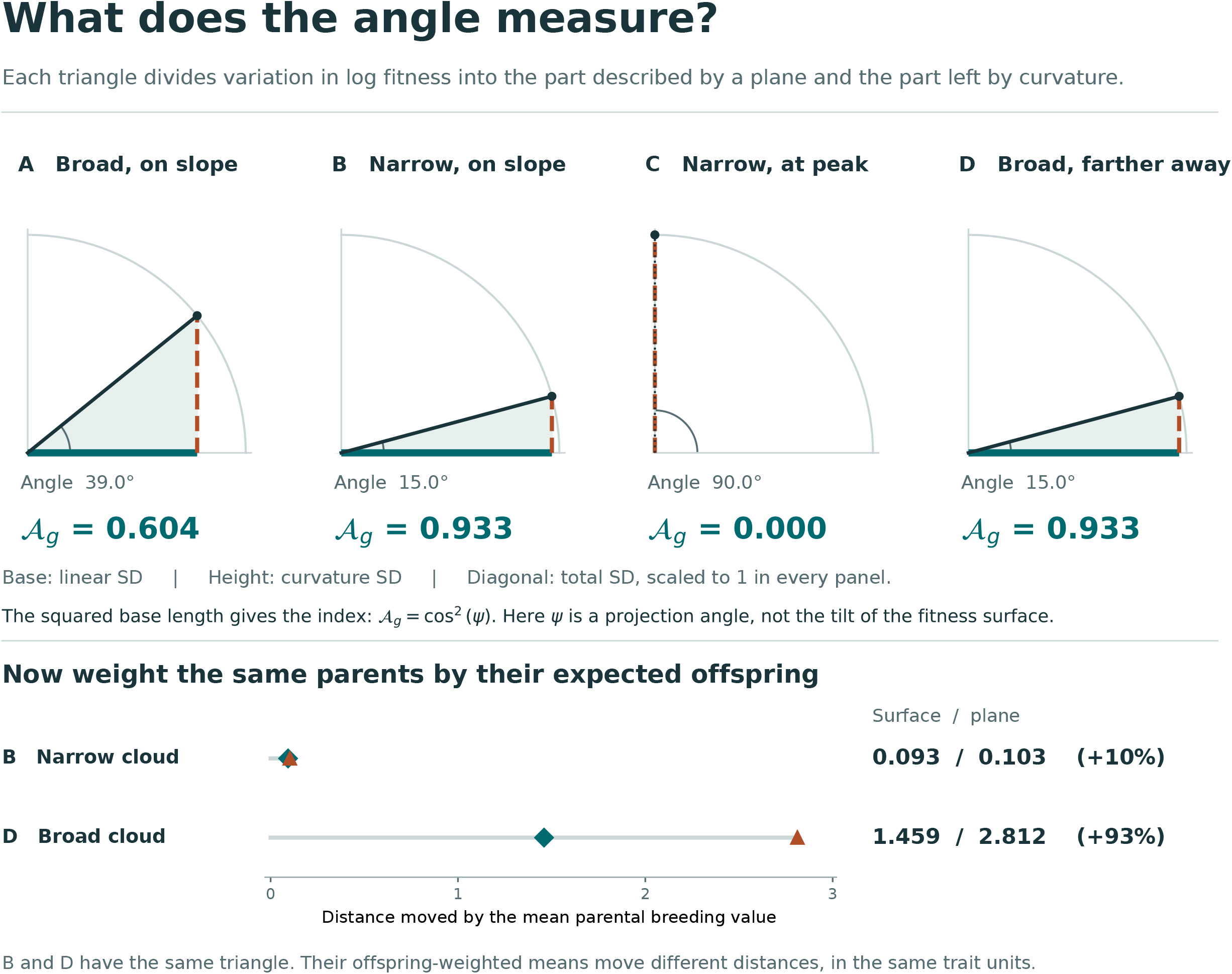

**Supplementary File 4:**
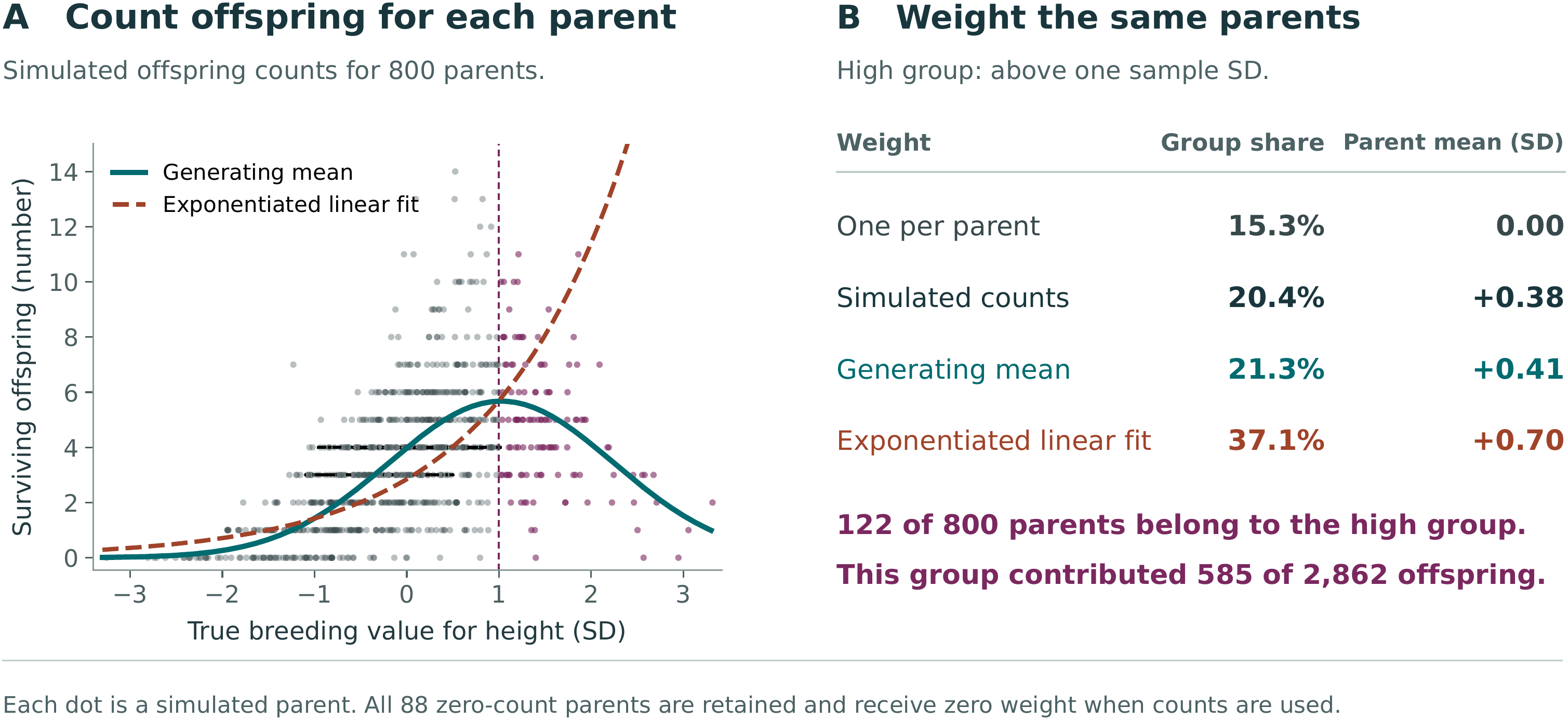
Parental weighting in a simulated population. Additive Channels in Curved Fitness Landscapes | Daniel Ortiz-Barrientos and Mark Cooper **Parent frequency and reproductive output assign different weights to the same breeding values.** (A) Each point represents one of 800 simulated parents at its known breeding value and realised count of surviving offspring. Breeding values are centred on the starting sample mean and divided by the sample SD, calculated with divisor 800. Counts were drawn from Poisson distributions with conditional mean 4 exp(0.7a - 0.35a^2^), where a is the unstandardised breeding value in centimetres. The solid curve shows this generating mean. The dashed curve exponentiates an ordinary least-squares line, including an intercept, fitted to the known log conditional mean across these parents; it was not fitted to the offspring counts. Purple points identify the 122 parents above one sample SD. (B) Each row normalises its weights over all 800 parents to calculate the high group’s contribution and the weighted parental mean. The high group accounts for 585 of 2,862 simulated offspring. All 88 zero-count parents remain in the starting population and have zero weight under realised reproduction. These calculations use the simulation truth from Supplementary File 3; they do not re-estimate breeding values from noisy height measurements. The means describe parental contributions. No offspring breeding values or inheritance process are simulated. Data and calculations: [parental-weighting module in the accompanying repository](https://github.com/dortizbarrientos/additive_channels/tree/55dda24e46979ce4cac000599fa6af62090319ce/tutorials/parental_weighting). The numerical values reproduce the archived example.

## References

Nicholas H. Barton and Michael Turelli. Natural and sexual selection on many loci. Genetics, 127(1): 229–255, 1991. doi: 10.1093/genetics/127.1.229.

Mark W. Blows and Robert Brooks. Measuring nonlinear selection. The American Naturalist, 162(6): 815–820, 2003. doi: 10.1086/378905.

M. G. Bulmer. Linkage disequilibrium and genetic variability. Genetical Research, 23(3):281–289, 1974. doi: 10.1017/S0016672300014920.

Michael G. Bulmer. The effect of selection on genetic variability. The American Naturalist, 105(943): 201–211, 1971. doi: 10.1086/282718.

Ashley J. R. Carter, Joachim Hermisson, and Thomas F. Hansen. The role of epistatic gene interactions in the response to selection and the evolution of evolvability. Theoretical Population Biology, 68(3): 179–196, 2005. doi: 10.1016/j.tpb.2005.05.002.

James M. Cheverud and Eric J. Routman. Epistasis and its contribution to genetic variance components. Genetics, 139(3):1455–1461, 1995. doi: 10.1093/genetics/139.3.1455.

Pierre de Villemereuil, Holger Schielzeth, Shinichi Nakagawa, and Michael B. Morrissey. General methods for evolutionary quantitative genetic inference from generalized mixed models. Genetics, 204(3): 1281–1294, 2016. doi: 10.1534/genetics.115.186536.

R. A. Fisher. Average excess and average effect of a gene substitution. Annals of Eugenics, 11(1):53–63, 1941. doi: 10.1111/j.1469-1809.1941.tb02272.x.

Ronald A. Fisher. The correlation between relatives on the supposition of Mendelian inheritance. Transactions of the Royal Society of Edinburgh, 52(2):399–433, 1918. doi: 10.1017/S0080456800012163.

Jarrod D. Hadfield, Alastair J. Wilson, Dany Garant, Ben C. Sheldon, and Loeske E. B. Kruuk. The misuse of BLUP in ecology and evolution. The American Naturalist, 175(1):116–125, 2010. doi: 10.1086/648604.

Benjamin C. Haller, Peter L. Ralph, and Philipp W. Messer. SLiM 5: Eco-evolutionary simulations across multiple chromosomes and full genomes. Molecular Biology and Evolution, 43(1):msaf313, 2026. doi: 10.1093/molbev/msaf313.

Thomas F. Hansen. Why epistasis is important for selection and adaptation. Evolution, 67(12):3501–3511, 2013. doi: 10.1111/evo.12214.

Thomas F. Hansen and David Houle. Measuring and comparing evolvability and constraint in multivariate characters. Journal of Evolutionary Biology, 21(5):1201–1219, 2008. doi: 10.1111/j.1420-9101.2008.01573.x.

Thomas F. Hansen, José M. Álvarez-Castro, Ashley J. R. Carter, Joachim Hermisson, and Günter P. Wagner. Evolution of genetic architecture under directional selection. Evolution, 60(8):1523–1536, 2006. doi: 10.1111/j.0014-3820.2006.tb00498.x.

L. Isserlis. On a formula for the product-moment coefficient of any order of a normal frequency distribution in any number of variables. Biometrika, 12(1–2):134–139, 1918. doi: 10.1093/biomet/12.1-2.134.

Russell Lande. Quantitative genetic analysis of multivariate evolution, applied to brain:body size allometry. Evolution, 33(1Part2):402–416, 1979. doi: 10.1111/j.1558-5646.1979.tb04694.x.

Russell Lande and Stevan J. Arnold. The measurement of selection on correlated characters. Evolution, 37(6):1210–1226, 1983. doi: 10.1111/j.1558-5646.1983.tb00236.x.

Richard C. Lewontin and Daniel Cohen. On population growth in a randomly varying environment. Proceedings of the National Academy of Sciences of the United States of America, 62(4):1056–1060, 1969. doi: 10.1073/pnas.62.4.1056.

Jacob A. Moorad and Michael J. Wade. Selection gradients, the opportunity for selection, and the coefficient of determination. The American Naturalist, 181(3):291–300, 2013. doi: 10.1086/669158.

Michael B. Morrissey and I. B. J. Goudie. Analytical results for directional and quadratic selection gradients for log-linear models of fitness functions. Evolution, 76(7):1378–1390, 2022. doi: 10.1111/evo.14486.

Patrick C. Phillips and Stevan J. Arnold. Visualizing multivariate selection. Evolution, 43(6):1209–1222, 1989. doi: 10.1111/j.1558-5646.1989.tb02569.x.

Dolph Schluter. Adaptive radiation along genetic lines of least resistance. Evolution, 50(5):1766–1774, 1996. doi: 10.1111/j.1558-5646.1996.tb03563.x.

John R. Stinchcombe, Aneil F. Agrawal, Paul A. Hohenlohe, Stevan J. Arnold, and Mark W. Blows. Estimating nonlinear selection gradients using quadratic regression coefficients: Double or nothing? Evolution, 62(9):2435–2440, 2008. doi: 10.1111/j.1558-5646.2008.00449.x.

Frank Technow, Carlos D. Messina, L. Radu Totir, and Mark Cooper. Integrating crop growth models with whole genome prediction through approximate Bayesian computation. PLOS ONE, 10(6):e0130855, 2015. doi: 10.1371/journal.pone.0130855.

Michael Turelli and Nicholas H. Barton. Genetic and statistical analyses of strong selection on polygenic traits: What, me normal? Genetics, 138(3):913–941, 1994. doi: 10.1093/genetics/138.3.913.

Bruce Walsh and Michael Lynch. Evolution and Selection of Quantitative Traits. Oxford University Press, Oxford, 2018. ISBN 9780198830870. doi: 10.1093/oso/9780198830870.001.0001. Online ISBN 9780191868986.

