## Supplementary material for "Additive Channels in Curved Fitness Landscapes": Supp Material 1

### A Step-by-Step Guide to the Mathematics of "Additive Channels in Curved Fitness Landscapes"

**Abstract**

Why can the same fitness surface look nearly linear in one population and show a substantial contribution from curvature in another? The answer depends on the breeding values present. This guide develops that result for one trait, starting with average reproductive output and working through the variance calculation. The same example then shows why a linear term that accounts for 97% of variation in log expected fitness can still overestimate the change among selected parents by 50%. The worked answers use one trait first. The final section shows what changes when the breeding values of several traits covary. The guide accompanies Appendices S1–S5 of the paper; no prior calculus or linear algebra is required for the main reading path.

**10 Answers to the checks** **33**

**11 Glossary** **36**

### 1 Reading this guide

Plants can differ in height because of their alleles, their growing conditions and interactions among these influences. Which genetic differences in height are associated with greater reproductive success? We will describe those differences using breeding values, then calculate average reproductive output at each value. To understand variation in a population, we also need the frequency of the breeding values. A curve supplies the expected output; a distribution supplies the frequencies. This guide explains how the two enter the calculation.

We will use one numerical example to calculate the contributions of slope and curvature, then ask what those calculations tell us about selection. Begin with Sections 2–5; they establish the central result and its interpretation. Section 6 adds inheritance, and Sections 7 and 8 connect the analysis to simulations and measurements. The integral in the optional derivation can be skipped on a first reading. Section 9 introduces matrices only after the one-trait calculation is complete.

Keep three quantities separate as you work through the example:

1. Individual reproductive output, which can differ even among individuals with the same breeding value.
2. Expected reproductive output at a given breeding value, found by averaging those individual outcomes.
3. The mean breeding value among parents, weighted by their reproductive contributions. Predicting the offspring mean also requires assumptions about transmission.

The calculations use familiar ideas from quantitative genetics in combination. Breeding values come from the regression of traits on allelic content [Fisher, 1918, 1941]; averaging over environmental deviations gives expected fitness at each additive genetic value [Lande, 1979]. Linear and quadratic regressions describe selection [Lande and Arnold, 1983], and their explained variances can be compared [Moorad and Wade, 2013]. Log-link models connect this regression approach to proportional changes in expected fitness [Morrissey and Goudie, 2022]. We introduce the results on selection-generated linkage disequilibrium when we reach inheritance [Bulmer, 1971, Barton and Turelli, 1991]. The question throughout is what these calculations say about the population being studied.

**Where to find each derivation**

| Paper part | Formal content | Teaching route here |
|---|---|---|
| Appendix S1 | Gaussian moments, variance formulas, population projection, coordinate invariance, and boundary cases | Sections 3 and 4; optional Sections 9.1 and 9.2 |
| Appendix S2 | Gaussian selection, selected-parent moments, displacement bound, and environmental averaging | Section 5; optional Section 9.3 |
| Appendix S3 | Fixed-baseline covariance recursion, fixed point, bounds, scalar convergence, and the $\rho = 1$ boundary | Section 6; optional Section 9.4 |
| Appendix S4 | Scope of the forward-time simulation | Section 7 |
| Appendix S5 | Conversion from a compatible phenotype-space selection regression | Section 8; optional Section 9.5 |
| Appendix S6 | Eleven anticipated questions | In the paper; each answer cites the paper's equations |

#### 2 Which fitness are we describing?

##### 2.1 Why use breeding values?

A plant's measured height is its phenotype. Its breeding value for height is the prediction obtained by adding the average effects of its alleles in a specified population [Fisher, 1918, 1941]. These effects come from a linear description of genetic differences in the trait using allele counts. Estimating them from measured phenotypes requires accounting for environmental effects. They can be defined even when alleles interact biologically. Adding average effects to form a predictor is therefore a different statement from assuming that alleles act independently in producing height.

Breeding values retain the units of the trait. A difference of two centimeters between breeding values means a two-centimeter difference between the additive genetic predictions of height. The plants' actual heights can differ by another amount because the prediction does not include every genetic and environmental influence. We use breeding values because we want to relate reproductive success to this additive genetic component. The value alone does not specify an offspring's phenotype: that also depends on the other parent, segregation, recombination and environment.

##### 2.2 Average reproductive output at a breeding value

For a plant, a fitness measure might be the number of offspring that survive from its seeds. Two plants with the same breeding value can leave different numbers of offspring. They can differ in genotype, growing conditions, damage or unmeasured traits. Write their individual output during the chosen selection episode as $W$. Write $A$ for breeding value and $a$ for a particular value. We ask for the mean output among individuals with that value:

$$W_A(a) = \mathbb{E}[W \mid A = a] \tag{1}$$

The symbol $\mathbb{E}$ means an average, and the bar means "among individuals with $A = a$." Thus $W_A(a)$ averages the reproductive output associated with breeding value $a$. In field data, two plants rarely have exactly the same estimated breeding value. We therefore fit a curve to estimate this conditional mean; grouping defines the target of the fit.

Repeating the average across breeding values gives a fitness curve. With several traits, the same operation gives a fitness surface over combinations of breeding values. It summarizes variation that a map assigning fitness to each genotype or phenotype would keep separate. It also depends on the population and environment in which the averages are taken. For example, a change in growing conditions among plants with a given breeding value can change their mean output without changing the numerical value used to group them.

#### 2.3 Average first, then take the logarithm

Selection depends on relative reproductive contributions. An increase from one expected offspring to two and an increase from ten to twenty each double the contribution, although their absolute increases differ. Logarithms give equal differences to equal ratios. We therefore take the logarithm of the conditional mean:

$$\ell_A(a) = \log W_A(a) = \log\{\mathbb{E}[W \mid A = a]\}. \tag{2}$$

The natural logarithm used here is the inverse of the exponential function: $\exp\{\log(x)\} = x$ for positive $x$. Thus a difference on the log scale can always be converted back to a ratio of expected outputs. For two breeding values,

$$\ell_A(a_2) - \ell_A(a_1) = \log\left\{\frac{W_A(a_2)}{W_A(a_1)}\right\}. \tag{3}$$

A log difference of 0.10 therefore means an expected-fitness ratio of $\exp(0.10) = 1.105$: about 10.5% more expected offspring. A straight line on the log scale describes a constant proportional change in expected output per unit change in breeding value. It need not be a straight line on the original output scale. Taking the exponential of a fitted log value also gives a positive expected output. These properties explain the use of log-link fitness models [Morrissey and Goudie, 2022]. The rest of the guide uses "log expected fitness" for the conditional quantity in Equation (2).

The order of operations is important. In general,

$$\log\{\mathbb{E}[W \mid A]\} \neq \mathbb{E}[\log W \mid A]. \tag{4}$$

Suppose two individuals with the same breeding value leave zero and two offspring. Their mean output is one, and $\log(1) = 0$. Averaging their logged outputs cannot give that answer because $\log(0)$ is not finite. This is why the order matters: first calculate the expected reproductive contribution, then describe it on the log scale.

> **Common misreading.** "Log expected fitness" here always means the logarithm of conditional expected fitness. It does not mean the expected logarithm of realized fitness.

#### 2.4 What the population distribution adds

Imagine plotting one point for each plant at its breeding value for height. The concentration of points shows which values are common. With two traits, each point has two coordinates, and the plot shows how breeding values for the traits vary together. This is the population distribution sometimes drawn as a "cloud." Its mean gives its position. Its variances give the spread along each trait axis, and its covariance describes their association.

The distribution and the fitness surface answer separate questions. A fitness contour joins combinations with the same expected reproductive output. A density contour joins combinations with equal population density; choosing its level can enclose a specified proportion of the population. A region can have high expected fitness yet contain few individuals. Its contribution to a population calculation depends on both facts.

For example, suppose breeding values of 0 and 2 cm have mean reproductive outputs of one and two offspring. In one population, 25% of plants have the higher value; in another, 75% do. Mean output is then $0.75(1) + 0.25(2) = 1.25$ in the first population and $0.25(1) + 0.75(2) = 1.75$ in the second. The expected output at each value was unchanged; its frequency changed. The same principle determines how much each value contributes when we calculate the variance of a fitness description.

This weighting is already part of classical selection analysis [Lande and Arnold, 1983]. Our calculation uses it to separate variation in log expected fitness into linear and curved contributions. Holding a surface fixed while changing the distribution lets us examine the effect of frequency or covariance alone. In biological comparisons, the conditional fitness curve can change as well and must be estimated for each population.

##### 2.5 How frequency and fitness determine reproductive contributions

To calculate each class's share of expected reproductive contributions, multiply its frequency by its mean output, then divide by the population's mean output. For continuously distributed breeding values, let $p_A(a)$ be the initial density, $W_A(a)$ the conditional mean fitness and $\bar{W}$ the population mean fitness. The same weighting rule gives

$$p_{\text{sel}}(a) = \frac{W_A(a) p_A(a)}{\bar{W}}. \tag{5}$$

In the preceding two-class example, the plants with breeding value 2 cm supply $0.25(2)/1.25 = 0.40$ of expected reproductive contributions in the first population and $0.75(2)/1.75 \simeq 0.857$ in the second. The weights are expected offspring numbers, not their logarithms. The resulting distribution describes parents weighted by their contributions. Inheritance is a further step because an individual's breeding value need not be transmitted unchanged to an offspring.

Figure 1 shows how the expected-fitness curve is constructed. The vertical groups show variation in reproductive output at a given breeding value; they do not show the frequency of breeding values in a population. The second panel shows the logarithm of each conditional mean.

> **Check 1.** Individuals with one breeding value have realized fitness values 0 and 2. Individuals with another have values 1 and 3. Calculate the logarithm of conditional expected fitness for each group. Why can the same calculation not be made by averaging the logarithms in the first group? See Answer 1.

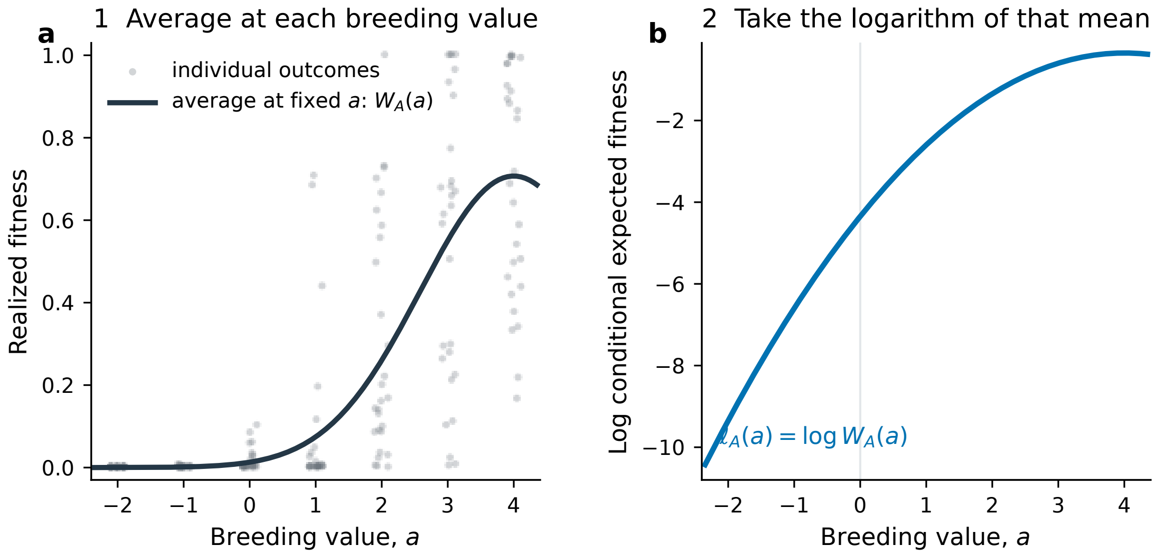

Figure 1: Construction of the paper's target. Individual realized-fitness values are first averaged at each breeding value to obtain $W_A(a)$, as in Equation (1). The logarithm is then taken. Weighting the preselection distribution by $W_A(a)$ gives the selected-contributor distribution in Equation (5); that final weighting step is not plotted. The vertical samples illustrate variation at fixed breeding values, rather than the population frequency of those values. None of these operations specifies inheritance.

*Alt text:* The first panel groups individual reproductive outputs vertically by breeding value and shows the conditional expected-fitness curve through their averages. The second panel shows the logarithm of that curve.

#### 3 One trait: from phenotype to expected fitness

We will now choose an explicit model so that every calculation can be worked out. Think of the trait as plant height, measured relative to a reference mean in arbitrary units. In this example, phenotype is the sum of breeding value and an independent environmental deviation. This is an additional assumption of the example, rather than a consequence of defining a breeding value:

$$Z = A + E. \tag{6}$$

Let breeding value $A$ and environmental deviation $E$ each follow a normal distribution with mean zero and variance one. The notation $N(0, 1)$ states those two numbers. Independence means that knowing one component does not change the distribution of the other. Expected reproductive output is highest at phenotype 4 and declines on either side, according to

$$W(Z) = W_{\max} \exp\left\{-\frac{(Z-4)^2}{2}\right\}. \tag{7}$$

Here $W_{\max}$ is the maximum expected output, and the width parameter is one. Plants with the same breeding value have different phenotypes because their environmental deviations differ. Averaging fitness over those deviations gives

$$W_A(a) = \frac{W_{\max}}{\sqrt{2}} \exp\left\{-\frac{(a-4)^2}{4}\right\}. \tag{8}$$

The factor $1/\sqrt{2}$ changes the height of the fitness curve, while the denominator four makes it broader than the phenotype–fitness curve. Taking the logarithm and expanding $(a-4)^2 = a^2 - 8a + 16$ gives

$$\begin{aligned} \ell_A(a) &= \log\left(\frac{W_{\max}}{\sqrt{2}}\right) - \frac{(a-4)^2}{4} \\ &= \ell_0 + 2a - \frac{1}{4}a^2, \end{aligned} \tag{9}$$

The constant $\ell_0 = \log(W_{\max}/\sqrt{2}) - 4$ is log expected fitness at the current mean, $a = 0$. Because it is added to every value, it does not affect differences among individuals or either variance component.

##### 3.1 Slope and curvature without calculus

Near the population mean, a small change in breeding value produces a change in log expected fitness. Their ratio is the slope, $b$. We can estimate it without taking a derivative: choose two nearby values, $-h$ and $h$, and divide the change in log expected fitness by their separation:

$$b \approx \frac{\ell_A(h) - \ell_A(-h)}{2h}. \tag{10}$$

If the slope changes along the curve, a single line will miss part of that relationship. Curvature, $H$, measures this change in slope. Using the values at $-h$, zero and $h$, we can calculate it as

$$H \approx \frac{\ell_A(h) - 2\ell_A(0) + \ell_A(-h)}{h^2}. \tag{11}$$

For our quadratic example, these formulas give $b = 2$ and $H = -0.5$ for any nonzero $h$. The positive slope means that an increase in breeding value near the mean increases log expected fitness. The negative curvature means that this increase becomes smaller as breeding value rises towards the optimum. Calculus calls $b$ and $H$ the first and second derivatives; the numerical example lets us use them before needing that notation.

Near the current mean, the local description is

$$\widetilde{\ell}_A(X) = \ell_0 + bX + \frac{1}{2}HX^2, \tag{12}$$

where $X = A - \mathbb{E}[A]$ is the breeding value measured from its current mean. The tilde over $\ell$ denotes an approximation. In the recurring example $\ell_A$ is exactly quadratic, so Equation (12) is exact rather than approximate.

**Check 2.** Use $h = 0.2$ and omit the constant $\ell_0$. Equation (9) gives $\ell_A(0.2) - \ell_0 = 0.39$ and $\ell_A(-0.2) - \ell_0 = -0.41$. Use Equations (10) and (11) to recover $b$ and $H$. See Answer 2.

##### 3.2 Optional derivation of the environmental average

Equation (8) is all we need for the next section. The following calculation shows where it comes from. Readers who want to continue with the variance partition can go directly to Section 4.

Here is the averaging step explicitly. Conditional on $A = a$, write one possible environmental deviation as $e$. Its normal density supplies the first exponential below, and phenotype-dependent fitness supplies the second:

$$W_A(a) = \frac{W_{\max}}{\sqrt{2\pi}} \int_{-\infty}^{\infty} \exp\left(-\frac{e^2}{2}\right) \exp\left\{-\frac{(a+e-4)^2}{2}\right\} de. \tag{13}$$

The integral sign means to take the continuous weighted average over all possible values of $e$. Put $\delta = a - 4$. The two squared terms combine as

$$e^2 + (e+\delta)^2 = 2\left(e + \frac{\delta}{2}\right)^2 + \frac{\delta^2}{2}. \tag{14}$$

The part involving $e + \delta/2$ is another bell-shaped curve. Its total area, together with the normalizing factor $1/\sqrt{2\pi}$, is $1/\sqrt{2}$; the part that still depends on $a$ is $\exp(-\delta^2/4)$. Their product is exactly Equation (8). No later calculation in the one-trait guide requires performing this integral again.

Appendix S2 gives the same calculation with general parameters. Let the environmental variance be $V_E$, let the optimum be $\theta$, and let $\omega^2$ describe the width of the phenotype–fitness relationship:

$$E \sim N(0, V_E), \qquad W(Z) = W_{\max} \exp\left\{-\frac{(Z-\theta)^2}{2\omega^2}\right\}. \tag{15}$$

Writing $\delta = a - \theta$, the two terms in the exponent can be rearranged as

$$\frac{e^2}{V_E} + \frac{(e+\delta)^2}{\omega^2} = \frac{V_E + \omega^2}{V_E\omega^2}\left(e + \frac{V_E\delta}{V_E+\omega^2}\right)^2 + \frac{\delta^2}{V_E+\omega^2}. \tag{16}$$

The first term on the right is a shifted bell-shaped curve in $e$. Averaging that curve supplies the scale factor below; the final term supplies the part that varies with $a$:

$$W_A(a) = W_{\max}\sqrt{\frac{\omega^2}{\omega^2+V_E}} \exp\left\{-\frac{(a-\theta)^2}{2(\omega^2+V_E)}\right\}. \tag{17}$$

Setting $V_E = \omega^2 = 1$ and $\theta = 4$ recovers Equation (8). The denominator $\omega^2 + V_E$ also shows directly why environmental variation makes the relationship in breeding-value space broader than the original relationship in phenotype space.

### 4 How much variation comes from slope and curvature?

#### 4.1 Averages, variance, and covariance

For any quantity $Y$, $\mathbb{E}[Y]$ is its population average. Its variance is

$$\mathrm{Var}(Y) = \mathbb{E}\left[(Y - \mathbb{E}[Y])^2\right]. \tag{18}$$

Variance is an average squared distance from the mean. It is nonnegative and describes the spread of a quantity. Covariance describes whether two quantities change together:

$$\mathrm{Cov}(U, V) = \mathbb{E}[(U - \mathbb{E}[U])(V - \mathbb{E}[V])]. \tag{19}$$

Positive covariance means that large values tend to occur together. Negative covariance means that a large value of one tends to accompany a small value of the other. Zero covariance means that there is no linear association; it does not necessarily mean statistical independence.

For a normal variable $X$ with mean 0 and variance $G$, the moments used in Appendix S1 are

$$\mathbb{E}[X] = 0, \qquad \mathbb{E}[X^2] = G, \qquad \mathbb{E}[X^3] = 0, \qquad \mathbb{E}[X^4] = 3G^2. \tag{20}$$

Here a moment is simply an average of a power, such as $\mathbb{E}[X^2]$ or $\mathbb{E}[X^4]$.

#### 4.2 Separating the linear and quadratic terms

The linear, or first-order, contribution is the slope multiplied by the deviation from the mean:

$$L = bX. \tag{21}$$

The quadratic term $HX^2/2$ does not generally have mean zero. Its mean is $HG/2$. Subtracting that mean gives

$$Q_c = \frac{1}{2}H(X^2 - G). \tag{22}$$

The subscript $c$ means centered: $Q_c$ has mean zero. Subtracting a constant shifts a curve up or down without changing how much its values differ, so this step leaves the quadratic variance unchanged. It makes the linear and quadratic terms easier to compare.

In the recurring example, $G = 1$, $b = 2$, and $H = -0.5$, so

$$L = 2X, \qquad Q_c = -\frac{1}{4}(X^2 - 1). \tag{23}$$

#### 4.3 Variance of the first-order term

If every deviation from the mean doubles, every squared deviation becomes four times as large. Variance therefore changes with the square of a multiplier. Applying that rule to $L = bX$ gives

$$V_{\text{lin}} = \text{Var}(L) = b^2 G. \tag{24}$$

For the example, $V_{\text{lin}} = 2^2 \times 1 = 4$.

#### 4.4 Variance of the quadratic term

For the curved contribution, we need the variance of $X^2$. Subtract the square of its mean from its mean square, using the normal moments above:

$$\begin{aligned} \text{Var}(X^2) &= \mathbb{E}[X^4] - \{\mathbb{E}[X^2]\}^2 \\ &= 3G^2 - G^2 = 2G^2. \end{aligned} \tag{25}$$

Consequently,

$$V_{\text{quad}} = \text{Var}(Q_c) = \frac{1}{2}H^2 G^2. \tag{26}$$

For the example, $V_{\text{quad}} = (1/2)(0.5)^2(1)^2 = 0.125$.

##### 4.5 Why the covariance is zero

A centered normal distribution is symmetric: $x$ and $-x$ have the same frequency. Replacing $x$ by $-x$ changes the sign of $L = bx$ but does not change $Q_c$, which depends on $x^2$. The product $LQ_c$ at $x$ is therefore the negative of the product at $-x$. The paired products cancel in the population average, giving

$$\mathrm{Cov}(L, Q_c) = 0. \tag{27}$$

For example, at $x = 2$, $L = 4$ and $Q_c = -0.75$, so $LQ_c = -3$. At $x = -2$, $L = -4$ and $Q_c = -0.75$, so $LQ_c = 3$.

Because the covariance is zero, the variance of their sum is

$$\mathrm{Var}(L + Q_c) = V_{\mathrm{lin}} + V_{\mathrm{quad}}. \tag{28}$$

The same cancellation gives the squared-correlation identity used in Appendix S1. When $V_{\mathrm{lin}} > 0$,

$$\begin{aligned}\mathrm{Corr}(L, L + Q_c)^2 &= \frac{\mathrm{Cov}(L, L + Q_c)^2}{\mathrm{Var}(L)\,\mathrm{Var}(L + Q_c)} \\ &= \frac{V_{\mathrm{lin}}^2}{V_{\mathrm{lin}}(V_{\mathrm{lin}} + V_{\mathrm{quad}})} = \frac{V_{\mathrm{lin}}}{V_{\mathrm{lin}} + V_{\mathrm{quad}}} = \mathcal{A}_g.\end{aligned} \tag{29}$$

This statement requires $L$ to have nonzero variance. The zero-variance cases are listed below.

##### 4.6 The first-order variance share

Provided the denominator is positive, define

$$\mathcal{A}_g = \frac{V_{\mathrm{lin}}}{V_{\mathrm{lin}} + V_{\mathrm{quad}}}. \tag{30}$$

For the example,

$$\mathcal{A}_g = \frac{4}{4 + 0.125} = 0.9697. \tag{31}$$

About 97% of the variance in our quadratic description of log expected fitness comes from the linear term. The curvature accounts for the remaining 3%. These percentages describe the fitted relationship across breeding values; individual reproductive output can vary further around that relationship.

This is what the paper calls an *additive channel*: a population whose variation in log expected fitness has a large linear share. The term refers to that relationship, not to interactions among alleles. The value is continuous, so its interpretation does not depend on crossing a universal threshold.

##### 4.7 The regression interpretation

A least-squares regression chooses the line that minimizes the average squared difference between the outcome and the line. The coefficient of determination, $R^2$, is the fraction of outcome variance accounted for by the fitted regression. This use of regression and explained variation has close precedents in quantitative-genetic analyses of selection [Lande and Arnold, 1983, Moorad and Wade, 2013].

Consider the outcome $L + Q_c$. The zero covariance derived above means that a linear regression on $X$ recovers $L = bX$ and cannot absorb any of $Q_c$. The line therefore accounts for $V_{\text{lin}}$ from a total variance of $V_{\text{lin}} + V_{\text{quad}}$. Hence

$$R_1^2 = \mathcal{A}_g. \tag{32}$$

The algebra is short. For $G > 0$, if $d$ is the fitted slope, the least-squares formula is

$$d = \frac{\text{Cov}(X, L + Q_c)}{\text{Var}(X)} = \frac{bG + 0}{G} = b. \tag{33}$$

The zero in the numerator is the covariance between $X$ and the quadratic term. With that contribution absent, the regression slope is exactly $b$. The variance-share calculation has recovered a familiar regression quantity.

For the original, uncentered curve, the fitted line has slope $b$ and intercept $\ell_0 + HG/2$. Thus it is parallel to the tangent at the mean but shifted by the average quadratic contribution. We write this intercept as $k$ below; the main paper uses $c$.

If an observed outcome can be written as

$$Y = k + L + Q_c + \varepsilon, \tag{34}$$

the residual $\varepsilon$ describes variation around the fitted relationship. Assume that its mean is zero at each breeding value and its variance is finite. Compare a linear fit, with coefficient of determination $R_1^2$, with a fit that also includes the quadratic term, with $R_{\leq 2}^2$. The two fits have the same outcome variance in their denominators, so

$$\frac{R_1^2}{R_{\leq 2}^2} = \mathcal{A}_g. \tag{35}$$

Equivalently, the fraction of the modeled first- and second-order signal added by the quadratic terms is

$$\frac{R_{\leq 2}^2 - R_1^2}{R_{\leq 2}^2} = 1 - \mathcal{A}_g. \tag{36}$$

For example, if $\text{Var}(\varepsilon) = 4$, then

$$R_1^2 = \frac{4}{8.125} = 0.4923, \qquad R_{\leq 2}^2 = \frac{4.125}{8.125} = 0.5077, \tag{37}$$

Their ratio is still 0.9697. The residual has reduced both coefficients of determination, but it has not changed the division between the model's linear and curved contributions. A large share can therefore coexist with a low $R^2$ for the observed outcome.

> **Common misreading.** Equation (35) does not generally justify regressing the logarithm of realized reproductive output. Such a regression targets $\mathbb{E}[\log W \mid A]$, not $\log \mathbb{E}[W \mid A]$. The identity applies when the fitted outcome and its residual satisfy the stated conditional-mean model on the required scale.

Figure 2 brings the two descriptions together. The upper panel shows the linear and curved contributions to log expected fitness. The density below it gives the frequency of breeding values used to calculate each variance. Values near the center receive more weight because more individuals occur there.

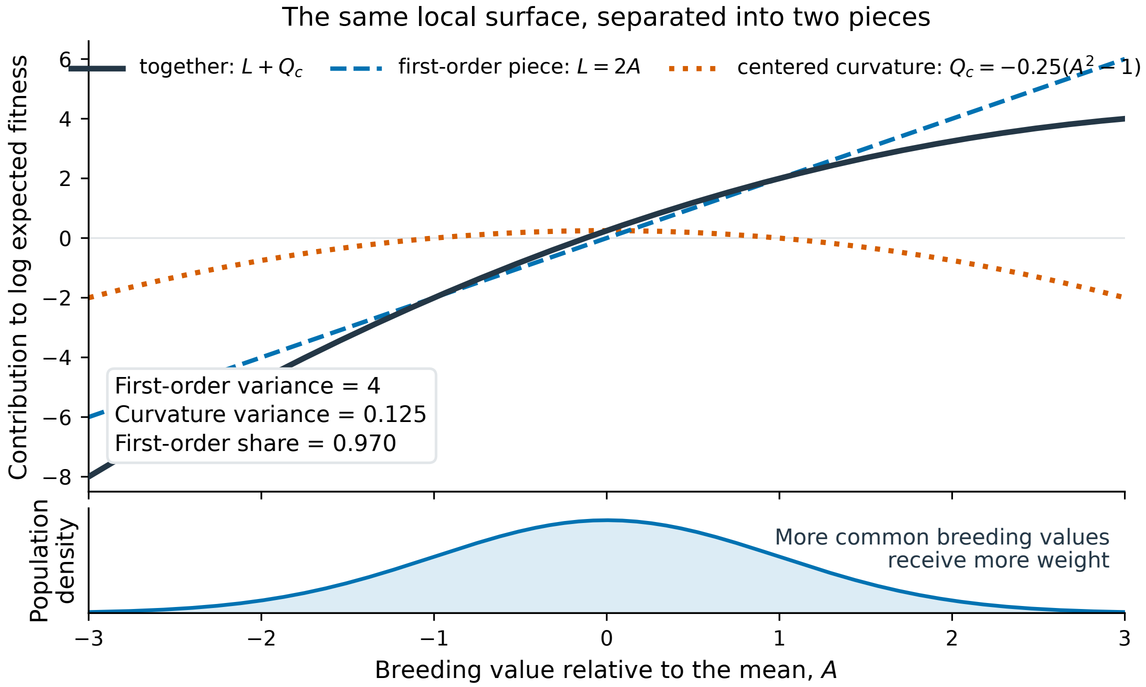

Figure 2: The Appendix S1 variance and population-projection results in one trait. The first-order term $L$ changes sign around the population mean, whereas the centered quadratic term $Q_c$ is symmetric. Their covariance is zero under the centered Gaussian distribution. The best affine projection therefore accounts for the fraction in Equation (30).

*Alt text:* The upper panel shows the full second-order curve for the logarithm of conditional expected fitness, its first-order line, and its centered quadratic term. The lower panel shows the bell-shaped breeding-value density that weights their variances.

#### 4.8 Changing population spread while holding the surface fixed

Suppose the breeding-value variance changes from $G$ to $sG$, where $s$ is a positive scaling factor, while $b$ and $H$ remain fixed. Equations (24) and (26) give

$$V_{\text{lin}}(s) = sV_{\text{lin}}(1), \qquad V_{\text{quad}}(s) = s^2 V_{\text{quad}}(1). \tag{38}$$

Halving the breeding-value variance therefore halves the linear contribution but reduces the quadratic contribution to one quarter. In our example, changing $G$ from 1 to 0.5 gives $V_{\text{lin}} = 2$, $V_{\text{quad}} = 0.03125$ and $\mathcal{A}_g = 0.9846$. The linear fraction has increased from about 97% to about 98.5%, even though both absolute components have decreased. Here "spread" means variance; it is the variance multiplier $s$ that is squared in the formula.

Nothing about the fitness curve changed in this comparison. The result isolates the effect of a narrower breeding-value distribution while holding slope and curvature fixed. Along an evolutionary trajectory, those quantities can also change and must be recalculated. For several traits, the same scaling result requires all variances and covariances to change in the same proportion.

#### 4.9 Changing units does not change the share

Suppose breeding value is rescaled from $a$ to $a' = 10a$. The variance becomes $G' = 100$, the slope per new unit becomes $b' = 0.2$, and the curvature per squared new unit becomes $H' = -0.005$. Substitution gives

$$(b')^2 G' = 0.2^2 \times 100 = 4 \tag{39}$$

and

$$\frac{1}{2}(H')^2(G')^2 = \frac{1}{2}(0.005)^2(100)^2 = 0.125. \tag{40}$$

Both variance terms, and therefore $\mathcal{A}_g$, are unchanged. Appendix S1 proves the corresponding result for any invertible change of coordinates involving several traits.

#### 4.10 Boundary cases

If $V_{\text{lin}} = 0$ and $V_{\text{quad}} > 0$, then $\mathcal{A}_g = 0$. The first-order regression is constant. A squared correlation involving $L$ is undefined because $L$ has no variance, even though the projection share can be recorded as zero. If both variance terms are zero, $\mathcal{A}_g$ and the correlation are both undefined because there is no local variation to divide between the terms.

A trait with no breeding-value variation contributes no variation to this calculation. With several traits, we keep the combinations that do vary and restrict any matrix inverses to those combinations.

**Check 3.** Recalculate $V_{\text{lin}}$, $V_{\text{quad}}$, and $\mathcal{A}_g$ for $G = 1$, $b = 2$, and $H = -0.5$. Then state what each number measures. See Answer 3.

**Check 4.** Explain why the products $LQ_c$ at $x = 2$ and $x = -2$ cancel. Which assumption about the distribution makes the two breeding values equally frequent? See Answer 4.

**Check 5.** After changing units to $a' = 10a$, verify the transformed values of $G$, $b$, and $H$ and confirm that $\mathcal{A}_g$ is unchanged. See Answer 5.

### 5 How selection changes the contributing parents

#### 5.1 Combining population frequencies and reproductive output

The preceding calculation compared variation in log expected fitness. To find the distribution among selected contributors, we must instead multiply the initial distribution by expected fitness itself, as in Equation (5). For normally distributed deviations $X$ with variance $G$, the initial density is proportional to

$$\exp\left(-\frac{X^2}{2G}\right). \tag{41}$$

"Proportional to" means that a constant needed to make the total probability one has been omitted. Under stabilizing curvature, write $\Gamma = -H$, so $\Gamma \geq 0$. The conditional expected-fitness weight, after ignoring a constant shared by all breeding values, is

$$\exp\left(bX - \frac{1}{2}\Gamma X^2\right). \tag{42}$$

Multiplying exponential terms adds their exponents. In the recurring example, $G = 1$, $b = 2$, and $\Gamma = 0.5$, so the combined exponent is

$$\begin{aligned} -\frac{1}{2}X^2 + 2X - \frac{1}{4}X^2 &= -\frac{3}{4}X^2 + 2X \\ &= -\frac{3}{4}\left(X - \frac{4}{3}\right)^2 + \frac{4}{3}. \end{aligned} \tag{43}$$

Writing the expression as a multiple of $(X - 4/3)^2$ identifies the center of the selected distribution. This operation is called completing the square. The constant $4/3$ affects only the factor needed to make the density integrate to one. The selected distribution has mean $4/3$ and variance $2/3$: the contributing parents have shifted towards the optimum and have less variation.

For general one-trait values,

$$g_{\text{sel}} = \frac{1}{1/g + \Gamma}, \qquad \Delta\bar{a}_{\text{full}} = g_{\text{sel}} b. \tag{44}$$

The symbol $g$ is the preselection breeding-value variance, $g_{\text{sel}}$ is the variance among selected contributors, and $\Delta\bar{a}_{\text{full}}$ is the change in their mean breeding value. The combined coefficient $1/g + \Gamma$ must be positive; otherwise the weighted distribution does not have a finite total. With stabilizing curvature $\Gamma \geq 0$ and $g > 0$, this condition is automatic.

Within the exact normal, log-quadratic model, $g_{\text{sel}}$ depends on curvature but not on $b$, whereas the selected mean depends on both. This separation need not hold for a nonnormal population or a nonquadratic function for the logarithm of conditional expected fitness.

#### 5.2 Why a high linear share can still give a poor selection prediction

If curvature were omitted, the predicted selected-parent displacement would be

$$\Delta\bar{a}_{\text{lin}} = gb. \tag{45}$$

Our example gave a linear share of 0.9697. Yet the linear-only calculation predicts a shift of two units, while the calculation including curvature gives 4/3. Dividing the difference by the full result gives

$$\frac{|2 - 4/3|}{|4/3|} = 0.5. \tag{46}$$

In one trait, when $b \neq 0$, the exact relative discrepancy is $|\Gamma g|$. Appendix S2 also proves

$$|\Gamma g| \leq \sqrt{2V_{\text{quad}}}. \tag{47}$$

Both sides equal 0.5 in one trait. Increasing $b$ enlarges the linear variance and pushes $\mathcal{A}_g$towards one, but leaves this relative error unchanged. The reason is the difference between the two calculations: the variance share compares squared deviations on the log scale, while the selected mean uses expected reproductive output to weight individuals. A good linear description for the first calculation need not be a good substitute in the second.

We have calculated a difference between the initial population and the parents weighted by their reproductive contributions. An inheritance model is needed to turn that result into a prediction for offspring.

Nor does the regression $R^2$ determine which individuals rank highest. On the far side of an optimum, expected fitness can decline while a linear score continues to rise. Ranking would require a separate comparison.

Figure 3 compares the initial distribution with the distribution among parents weighted by their expected reproductive contributions. It shows the change in both the mean and the variance calculated above.

#### 5.3 Why environmental variation changes the curvature

Equation (7) has phenotype-space curvature $-1$. After environmental deviations with variance 1 are averaged out, Equation (8) has breeding-value-space curvature $-1/(1+1) = -0.5$. Individuals with one breeding value occupy a range of phenotypes, so their mean fitness changes more gradually with breeding value than individual fitness changes with phenotype.

More generally, if phenotype-space fitness has width $\omega^2$ and the environmental variance is $V_E$, Gaussian averaging gives width $\omega^2 + V_E$ and curvature

$$H = -\frac{1}{\omega^2 + V_E}, \tag{48}$$

rather than $-1/\omega^2$. This is why phenotype-space and breeding-value- space curvature cannot be treated as interchangeable.

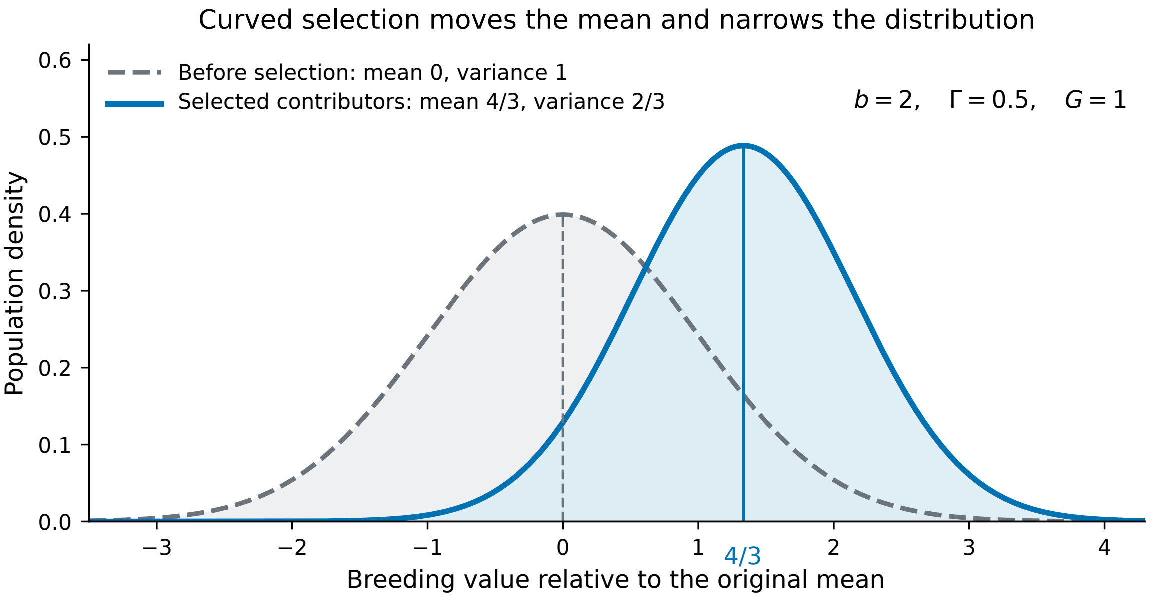

Figure 3: The Appendix S2 selected-parent moments and the distinction between the first-order variance share and displacement accuracy. Weighting $A \sim N(0, 1)$ by the recurring log-quadratic fitness function changes the mean from 0 to $4/3$ and the variance from 1 to $2/3$, as in Equation (44). The first-order displacement is 2, producing the relative discrepancy in Equation (47), although $\mathcal{A}_g = 0.9697$.

*Alt text:* The preselection and selected-contributor normal distributions are drawn with vertical lines at means zero and four thirds. Their variances are one and two thirds, respectively.

**Check 6.** Use Equation (44) with $g = 1$, $\Gamma = 0.5$, and $b = 2$. Calculate the selected variance and selected mean displacement. See Answer 6.

**Check 7.** Replace $b = 2$ by $b = 10$, keeping $g = 1$ and $\Gamma = 0.5$. Calculate $\mathcal{A}_g$ and the relative discrepancy between the first-order and full displacements. Explain why the two answers move in different ways. See Answer 7.

### 6 Adding inheritance: variation across generations

#### 6.1 The added inheritance assumptions

What happens to the reduced variance among selected parents when they reproduce? Recombination can reduce the associations between alleles created by selection. We now represent that process with a simple rule, while retaining the normal distribution and exactly quadratic log expected fitness used above. The one-trait update is

$$g_{t+1} = (1 - \rho)B + \rho \frac{g_t}{1 + \gamma g_t}. \tag{49}$$

Read this equation as a weighted average of two variances: the baseline $B$ and the selected variance $g_t/(1 + \gamma g_t)$. Its symbols are

- $g_t$ is breeding-value variance in generation $t$;
- $B$ is a fixed genic variance baseline;
- $\gamma \geq 0$ is fixed stabilizing curvature;
- $\rho$, between 0 and 1, is the fraction of the postselection covariance departure from the baseline retained after reproduction.

**Common misreading.** In this section, lowercase $\gamma$ follows the main paper's scalar recursion and denotes stabilizing curvature. Appendix S5 later uses lowercase $\gamma$ for the conventional quadratic coefficient in a phenotype-space selection regression. Those coefficients belong to different models and should not be interchanged. Uppercase $\Gamma = -H$ in Appendix S2 is the one-trait sign-reversed curvature corresponding to the present scalar $\gamma$.

The model assumes that allelic effects stay fixed and breeding values are their linear sums in every generation. It separately assumes a fixed genic variance $B$, the variance contributed by loci before accounting for associations between them. To see why the second assumption does not follow from the first, consider one biallelic locus. Its genic variance is $2p(1 - p)u^2$, where $u$ is the allelic effect. With $u = 1$, changing allele frequency from $p = 0.5$ to $p = 0.8$ reduces the variance from 0.50 to 0.32. The effect stayed fixed; its contribution to variance did not.

Holding $B$ fixed excludes the effects of such allele-frequency changes on genic variance from the update. The equation also omits mutation, drift, migration and changing allelic effects. It isolates how selection-generated associations and their retention affect variance under fixed parameters.

#### 6.2 A numerical iteration

Take

$$B = 1, \qquad \gamma = 0.5, \qquad \rho = 0.5. \tag{50}$$

Starting at $g_0 = 1$, the selected variance is $1/(1 + 0.5) = 2/3$. Reproduction then gives

$$g_1 = 0.5 + 0.5\left(\frac{2}{3}\right) = 0.8333. \tag{51}$$

Repeating the same calculation gives

$$g_2 = 0.7941, \qquad g_3 = 0.7842, \qquad g_4 = 0.7817. \tag{52}$$

Every update lies between

$$(1 - \rho)B = 0.5 \qquad \text{and} \qquad B = 1. \tag{53}$$

For this starting value, every iterate stays in the interval $[0.5, 1]$. The lower bound is the baseline contribution supplied at each update. The upper bound follows because selection reduces the current variance, and the update averages that selected variance with $B = 1$. A starting variance outside this interval need not enter it immediately.

##### 6.3 The scalar fixed point and convergence

A fixed point $g^*$ is a value that the update leaves unchanged. Setting $g_{t+1} = g_t = g^*$ gives

$$g^* = 0.5 + 0.5\frac{g^*}{1 + 0.5g^*}. \tag{54}$$

Multiplying both sides by $1 + 0.5g^*$ and collecting terms gives

$$2(g^*)^2 + g^* - 2 = 0. \tag{55}$$

The positive solution is

$$g^* = \frac{\sqrt{17} - 1}{4} = 0.7808. \tag{56}$$

For general $B > 0$, $\gamma > 0$, and $0 \le \rho < 1$, the same steps start from

$$g^* = (1 - \rho)B + \rho\frac{g^*}{1 + \gamma g^*}. \tag{57}$$

Multiplication by $1 + \gamma g^*$ gives

$$g^* + \gamma(g^*)^2 = (1 - \rho)B + (1 - \rho)\gamma B g^* + \rho g^*. \tag{58}$$

Moving every term to the left and collecting the two terms proportional to $g^*$ produces the quadratic displayed in Appendix S3:

$$\gamma(g^*)^2 + (1-\rho)(1-\gamma B)g^* - (1-\rho)B = 0. \tag{59}$$

The quadratic formula has one nonnegative root, namely

$$g^* = \frac{-(1-\rho)(1-\gamma B) + \sqrt{(1-\rho)^2(1-\gamma B)^2 + 4\gamma(1-\rho)B}}{2\gamma}. \tag{60}$$

When $\gamma = 0$, selection does not change variance and the fixed point is $g^* = B$ for $0 \le \rho < 1$.

Convergence can be shown without derivatives. Let $F(g)$ denote the right side of Equation (49). For any two nonnegative values $u$ and $v$,

$$|F(u) - F(v)| = \frac{\rho|u-v|}{(1+\gamma u)(1+\gamma v)} \le \rho|u-v|. \tag{61}$$

Since $\rho < 1$, two starting values become closer after each update. In particular, the distance from a starting value to a fixed point becomes at most $\rho$ times as large at every step. It tends to zero, proving convergence. Two different fixed points would have to become closer while remaining unchanged, which is impossible; the scalar fixed point is unique. The matrix proof in Appendix S3 establishes existence and bounds, but does not make those additional claims.

##### 6.4 The boundary $\rho = 1$

When $\rho = 1$, the baseline term disappears and

$$g_{t+1} = \frac{g_t}{1+\gamma g_t}. \tag{62}$$

For a positive starting variance, taking reciprocals gives

$$\frac{1}{g_{t+1}} = \frac{1}{g_t} + \gamma. \tag{63}$$

Thus precision, defined as inverse variance, rises by $\gamma$ each generation:

$$\frac{1}{g_t} = \frac{1}{g_0} + t\gamma. \tag{64}$$

If $\gamma > 0$, the variance approaches zero. There is no positive interior fixed point at this boundary. If $\gamma = 0$, every positive starting variance is fixed because the update becomes $g_{t+1} = g_t$. With several traits, variance can remain only in directions with exactly zero curvature; if $\mathbf{\Gamma} = \mathbf{0}$, every starting covariance is fixed.

The paper treats $\rho$ as a summary of linkage-disequilibrium retention, not as a universal function of a recombination rate or mating system. Its value depends on locus pairs, effects, signs of disequilibria, recombination architecture, and the population's mating system [Bulmer, 1974, Barton and Turelli, 1991].

Figure 4 shows the successive variance updates and their approach to the scalar fixed point. Each update uses the same baseline, curvature and retention parameter.

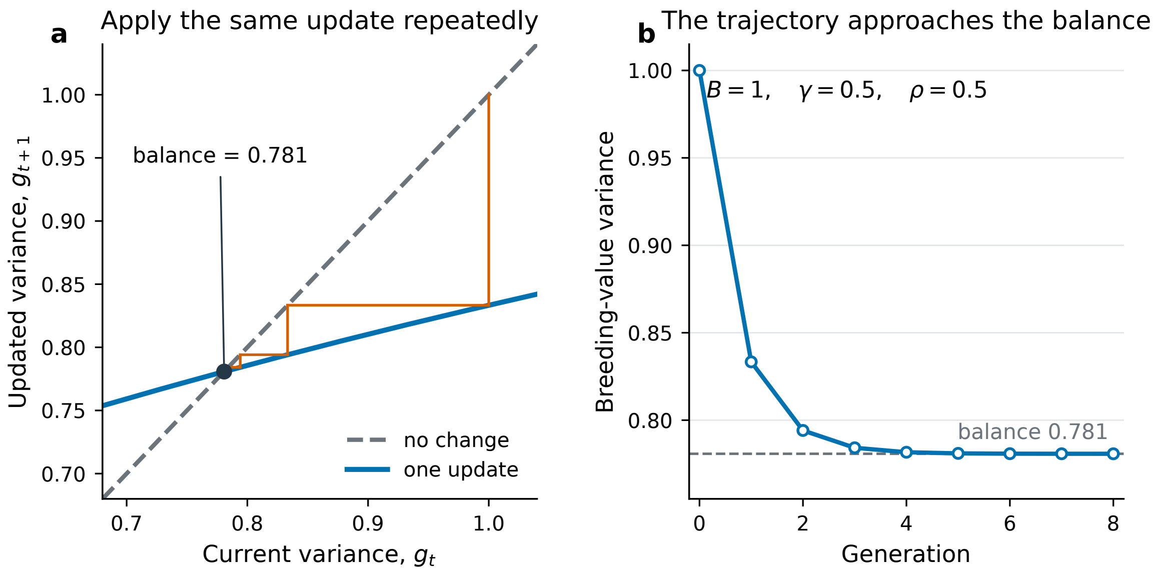

Figure 4: The scalar fixed-baseline covariance recursion of Appendix S3 for $B = 1$, $\gamma = 0.5$, and $\rho = 0.5$. Successive values remain in the interval $[(1-\rho)B, B] = [0.5, 1]$ and approach the conditional fixed point $g^* = 0.7808$. This is a covariance fixed point under fixed parameters, not a mutation–selection equilibrium.

*Alt text:* A plot of the scalar update against the identity line marks the fixed point at 0.7808. A short sequence beginning at variance one approaches the crossing while remaining between 0.5 and one.

**Check 8.** Starting from $g_0 = 1$, calculate $g_1$, $g_2$, and $g_3$ for $B = 1$, $\gamma = 0.5$, and $\rho = 0.5$. Are all three values inside the interval in Equation (53)? See Answer 8.

**Check 9.** Let $g_0 = 1$, $\gamma = 0.5$, and $\rho = 1$. Use Equation (64) to calculate $g_4$. Why is $B$ irrelevant at this boundary? See Answer 9.

### 7 Checking the approximation in simulated populations

The Gaussian formulas are exact under their stated assumptions. Breeding values in a finite population need not follow an exact normal distribution, so the paper also asks how closely the formula agrees with its squared-correlation counterpart in a multilocus simulation.

The paper's forward-time simulation used SLiM 5.2 [Haller et al., 2026]. It contained 20 replicate diploid populations of 2,000 individuals, using seeds 42–61. Two hundred loci contributed additively to four breeding values, with 50 loci per trait. The trait-specific standard deviations of allelic effects were 0.346, 0.283, 0.200, and 0.141. Stabilizing-selection strengths were 0.10, 0.05, 0.04, and 0.03. The optimum for trait one moved by 0.3 per generation for 40 generations, reaching 12, and then remained fixed. Each replicate ran for 200 generations. No environmental deviation was added in the simulation, so simulated phenotype equaled the additive trait value.

At each generation, the analysis estimated covariance and calculated the slope and curvature of log expected fitness. It then compared the formula for $\mathcal{A}_g$ with the squared sample correlation between the linear term and the full quadratic expression. Across the 20 replicates and 200 generations, the median absolute difference was 0.0157. A fresh run with the same seeds reproduced the archived diagnostics to numerical precision.

The agreement means that the Gaussian calculation remained close to the sample comparison in this simulated design. It does not establish an exact identity for every finite-population distribution. The simulation also did not test the displacement error bound or the fixed-baseline recursion.

Allele frequencies change through selection and drift in this simulation. Genic variance can therefore change even though the allelic effects remain fixed. That is why this simulation checks the variance-share approximation, while the recursion in Section 6 is a separate calculation with a fixed baseline.

You can make a simpler check using our one-trait example. Draw many values from $N(0, 1)$, calculate $L = 2X$ and $Q_c = -(X^2 - 1)/4$, then compare the squared sample correlation of $L$ and $L + Q_c$ with 0.9697. Independent samples give slightly different answers. As sample size grows, the estimate converges to the population value, although the error need not decrease at every increase in sample size.

**Check 10.** Suppose a simulation reproduces the first-order variance share closely but does not transmit parental breeding values to offspring using the fixed-$B$ recursion. Can it validate the recursion's fixed point? Explain. See Answer 10.

### 8 Using results from a phenotypic selection regression

#### 8.1 The regression being converted

Many selection studies measure phenotypes and reproductive output rather than breeding values. Under specified assumptions, a linear–quadratic regression from such a study can be used to calculate the selected breeding-value mean and variance. We can then find a log-quadratic model that gives the same two quantities. Begin with the classical regression of relative fitness on centered phenotype [Lande and Arnold, 1983]:

$$w = \alpha + \beta x + \frac{1}{2}\gamma x^2 + \varepsilon. \tag{65}$$

Relative fitness, $w$, is individual fitness divided by its population mean, so $\mathbb{E}[w] = 1$. Here $x$ is phenotype measured from its mean, $\alpha$ is the fitted intercept, and $\beta$ and $\gamma$ are the linear and quadratic coefficients. Least squares leaves a residual $\varepsilon$ with mean zero and zero covariance with $x$ and $x^2$. The factor $1/2$ is a convention: a software coefficient on $x^2$ must be doubled to give $\gamma$ if the software omits that factor [Stinchcombe et al., 2008].

Let phenotypic variance be $P = 1.5$, with $\beta = 0.4$ and $\gamma = -0.2$. Since $\mathbb{E}[w] = 1$, averaging Equation (65) gives

$$1 = \alpha + \frac{1}{2}\gamma P, \tag{66}$$

and therefore

$$\alpha = 1 - \frac{1}{2}\gamma P = 1.15. \tag{67}$$

The fitted value at mean phenotype is 1.15, not 1. Relative fitness equals one on average over the population; it need not equal one at the population mean.

##### 8.2 The two selected moments

For a centered normal phenotype, Equation (20) applies with $G$ replaced by $P$. Multiplying Equation (65) by $x$ and averaging gives

$$\mathbb{E}[xw] = P\beta. \tag{68}$$

This is the mean phenotype after weighting by relative fitness. Multiplying by $x^2$ and averaging gives

$$\mathbb{E}[x^2 w] = P + \gamma P^2. \tag{69}$$

Variance is the second moment minus the square of the mean, so selected phenotypic variance is

$$P + P^2(\gamma - \beta^2). \tag{70}$$

Now let centered breeding value $A$ have variance $G = 0.5$ and $\mathrm{Cov}(A, x) = G$. Under the joint-normal and selection-mediation assumptions stated below, the corresponding selected breeding-value mean and variance have a short derivation. For an individual with centered phenotype $x$, normal conditioning gives

$$\mathbb{E}[A \mid x] = \frac{G}{P}x, \qquad \mathrm{Var}(A \mid x) = G - \frac{G^2}{P}. \tag{71}$$

For these jointly normal variables, the predicted breeding value at phenotype $x$ is $Gx/P$. Individuals with that phenotype still differ in breeding value, with conditional variance $G - G^2/P$. To find the mean among selected contributors, average the prediction using relative fitness as the weight:

$$\mathbb{E}[A \mid \mathrm{sel}] = \frac{G}{P}\mathbb{E}[xw] = G\beta. \tag{72}$$

For the second moment, add the conditional variance to the squared conditional mean, then average with the same fitness weights:

$$\begin{aligned}\mathbb{E}[A^2 \mid \mathrm{sel}] &= G - \frac{G^2}{P} + \frac{G^2}{P^2}\mathbb{E}[x^2 w] \\ &= G + G^2\gamma. \end{aligned} \tag{73}$$

Subtracting the square of the selected mean gives

$$\Delta\bar{A}_{\mathrm{sel}} = G\beta, \qquad G_{\mathrm{sel}} = G + G^2(\gamma - \beta^2). \tag{74}$$

For the numerical values,

$$\Delta\bar{A}_{\mathrm{sel}} = 0.5(0.4) = 0.2, \qquad G_{\mathrm{sel}} = 0.5 + 0.25(-0.2 - 0.16) = 0.41. \tag{75}$$

#### 8.3 Matching those moments to a log-quadratic model

Appendix S2 gives the selected moments produced by a normal, log-quadratic breeding-value model. Requiring that model to produce the same selected mean and variance gives

$$b = \frac{G\beta}{G_{\mathrm{sel}}}, \qquad H = \frac{1}{G} - \frac{1}{G_{\mathrm{sel}}}. \tag{76}$$

For the example,

$$b = \frac{0.2}{0.41} = 0.4878, \qquad H = 2 - \frac{1}{0.41} = -0.4390. \tag{77}$$

These values of $b$ and $H$ were chosen to reproduce the selected mean and variance. A different calculation would take the logarithm of the fitted phenotype regression and differentiate it at the mean. Provided its intercept is positive, that would give

$$b_{\mathrm{raw}} = \frac{\beta}{\alpha} = 0.3478, \qquad H_{\mathrm{raw}} = \frac{\gamma}{\alpha} - \frac{\beta^2}{\alpha^2} = -0.2949. \tag{78}$$

The answers differ because the calculations describe different relationships. The values $b_{raw}$ and $H_{raw}$ describe the fitted regression on phenotype. Equation (76) describes a quadratic log-fitness relationship on breeding value that reproduces the selected breeding-value mean and variance.

Figure 5 follows the calculation from a compatible phenotype regression to the selected breeding-value mean and variance, then to the log-fitness model that matches those moments.

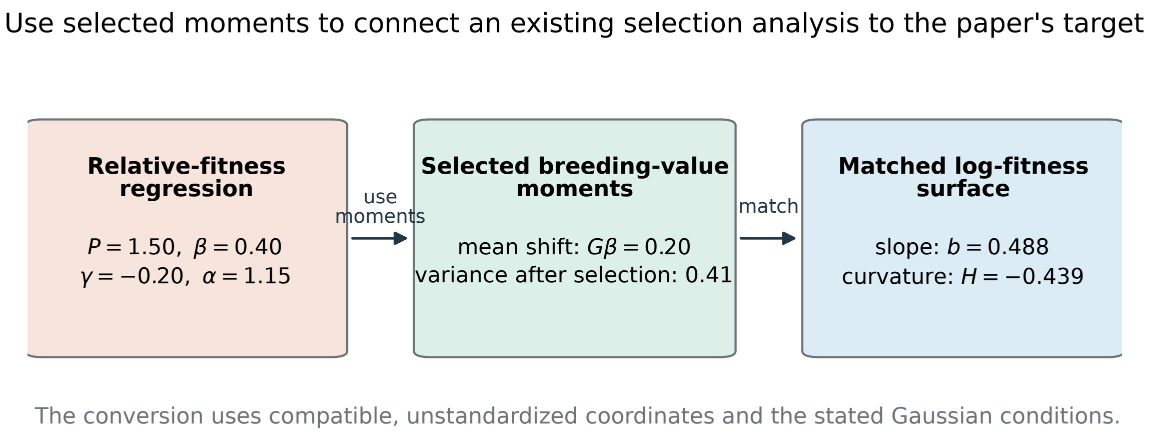

Figure 5: The Appendix S5 moment-conversion route. A compatible phenotype- space regression supplies $\beta$ and $\gamma$. Together with breeding-value variance $G$, these give the selected breeding-value mean and variance in Equation (74). Matching those moments to the Appendix S2 Gaussian selection result gives $b$ and $H$ through Equation (76). Direct differentiation of the phenotype regression targets a different function.

*Alt text:* A left-to-right diagram begins with phenotype regression coefficients beta and gamma plus breeding-value variance G, passes through selected breeding-value mean and variance, and ends with breeding-value-space slope b and curvature H.

#### 8.4 Conditions required for the conversion

Before making this conversion, check the following conditions. They connect the measured phenotypes to the breeding values whose fitness relationship we want to describe.

1. Coefficients are unstandardized or have been converted into compatible trait coordinates and the same quadratic convention.
2. The selection coefficients and $G$ refer to the same population and selection episode.
3. Breeding value and phenotype are jointly normal, or the required conditional-moment relations are otherwise justified.
4. Selection is mediated by the measured phenotypes, or the equivalent conditional-independence condition holds.
5. $G_{sel}$ is positive on every direction with breeding-value variation.
6. Uncertainty in $\beta$, $\gamma$, $G$, and every transformation is propagated to $b$, $H$, $V_{lin}$, $V_{quad}$, and $\mathcal{A}_g$.

If the moment conditions hold but exact log-quadraticity does not, the result is a moment-matched quadratic approximation. If the normality, mediation, coordinate, or positivity conditions fail, the conversion is not justified. It does not recover point derivatives of an arbitrary $\log \mathbb{E}[W \mid A]$.

In a direct empirical analysis, a conditional-mean model appropriate for the fitness response should target $\mathbb{E}[W \mid A]$ [Morrissey and Goudie, 2022]. Predicted breeding values should be treated as uncertain latent quantities rather than error-free measurements [Hadfield et al., 2010]. Generalized mixed models also require care when moving between latent and observed scales [de Villemereuil et al., 2016]. Regressing logged realized fitness is valid for the paper's target only under an additional error model that makes the gap in Equation (4) constant with breeding value.

Prediction uncertainty can also affect the variance share. Suppose $b$ and $H$ are known on the true breeding-value scale, but the variance of predicted breeding values is only $rG$, where $0 < r < 1$. If we mistakenly use that reduced variance as the population's biological variance, we obtain $rV_{\text{lin}}$ and $r^2 V_{\text{quad}}$. Their ratio gives

$$\widehat{\mathcal{A}g}_{\text{naive}} = \frac{V_{\text{lin}}}{V_{\text{lin}} + rV_{\text{quad}}}. \tag{79}$$

With reliability $r = 0.6$ in the recurring example, this gives $4/(4 + 0.6 \times 0.125) = 0.9816$, compared with the true value 0.9697. This direction of bias belongs only to the stated substitution. It is not a general correction when $b$ and $H$ are themselves estimated from predicted breeding values or when reliability differs among directions.

Supplementary File 3 shows how to estimate the share from noisy height measurements and offspring counts. Its Shiny app lets readers vary measurement error, compare the estimated components with their generating values, and check the consequences of a nonquadratic fitness relationship.

> **Check 11.** Using $P = 1.5$, $\beta = 0.4$, $\gamma = -0.2$, and $G = 0.5$, calculate $\alpha$, $G_{\text{sel}}$, $b$, and $H$. Why is $\alpha$ not one? See Answer 11.

#### 9 Extending the calculation to several traits

With several traits, individuals differ in combinations of breeding values. A covariance matrix records both the variation in each trait and the associations between traits. The operations are the same as in one trait—form a linear term, form a quadratic term, and calculate their variances—but the notation keeps track of all the trait combinations. We introduce that notation first and then work through the two-trait example from Figure 3 of the paper.

##### 9.1 Notation and operators

A *vector* is an ordered list of numbers. For $n$ traits, $\boldsymbol{X} = (X_1, \dots, X_n)^\mathsf{T}$ lists centered breeding values. The superscript $\mathsf{T}$, called transpose, changes a column into a row. The product $\boldsymbol{b}^\mathsf{T}\boldsymbol{X} = \sum_i b_i X_i$ is the weighted sum across traits; the symbol $\sum_i$ means add one term for every trait $i$.

A *matrix* is a rectangular table of numbers. The covariance matrix $\mathbf{G}$ has variances $G_{ii}$ on its diagonal and covariances $G_{ij}$ away from the diagonal. It is symmetric because $G_{ij} = G_{ji}$. Matrix multiplication forms weighted sums using rows of the first matrix and columns of the second.

The identity matrix $\mathbf{I}$ leaves a vector unchanged. An inverse $\mathbf{M}^{-1}$ undoes multiplication by $\mathbf{M}$, so $\mathbf{M}^{-1}\mathbf{M} = \mathbf{I}$. An inverse exists only on directions that the matrix does not collapse to zero. The *support* of $\mathbf{G}$ is the set of directions with positive breeding-value variance; inverse-based formulas for a singular $\mathbf{G}$ are evaluated only on that support.

The trace $\mathrm{tr}(\mathbf{M})$ is the sum of diagonal entries of $\mathbf{M}$. A quadratic form $\boldsymbol{x}^\top \mathbf{M}\boldsymbol{x}$ returns one number. A symmetric matrix is positive semidefinite, written $\mathbf{M} \succeq 0$, if this number is never negative. The notation $\mathbf{M} \preceq \mathbf{N}$ means that $\mathbf{N} - \mathbf{M}$ is positive semidefinite. For covariance matrices it says that $\mathbf{N}$ has at least as much variance as $\mathbf{M}$ in every direction.

An eigenvector is a vector whose image under a matrix is a scalar multiple of itself. The scalar is its eigenvalue; a negative value reverses its direction. For a covariance matrix, eigenvectors identify uncorrelated combinations of traits, and their eigenvalues give the corresponding variances. A positive semidefinite covariance matrix also has a symmetric positive semidefinite square root, $\mathbf{G}^{1/2}$, satisfying $\mathbf{G}^{1/2}\mathbf{G}^{1/2} = \mathbf{G}$.

The Euclidean norm $\|\boldsymbol{x}\| = (\boldsymbol{x}^\top \boldsymbol{x})^{1/2}$ is ordinary length. The covariance-scaled norm $\|\boldsymbol{x}\|_{\mathbf{G}^{-1}} = (\boldsymbol{x}^\top \mathbf{G}^{-1}\boldsymbol{x})^{1/2}$ measures a displacement relative to the population's covariance. The spectral norm $\|\mathbf{M}\|_2$ is the largest factor by which $\mathbf{M}$ stretches any unit vector. The Frobenius norm $\|\mathbf{M}\|_{\mathrm{F}}$ is the square root of the sum of squared matrix entries. The largest directional stretch cannot exceed that total root-sum-of-squares, so $\|\mathbf{M}\|_2 \le \|\mathbf{M}\|_{\mathrm{F}}$.

#### 9.2 Appendix S1 in several traits

The local terms are

$$L = \boldsymbol{b}^\top \boldsymbol{X}, \qquad Q_c = \frac{1}{2}\{\boldsymbol{X}^\top \mathbf{H}\boldsymbol{X} - \mathrm{tr}(\mathbf{H}\mathbf{G})\}. \tag{80}$$

For a centered multivariate normal distribution, the product-moment rule is

$$\mathrm{Cov}(X_i X_j, X_k X_l) = G_{ik}G_{jl} + G_{il}G_{jk}. \tag{81}$$

This is the several-trait counterpart of $\mathbb{E}[X^4] = 3G^2$ and is a form of the Gaussian fourth-moment identity [Isserlis, 1918]. Expanding the variance of the centered quadratic term means summing over every four-index combination:

$$\begin{aligned}\mathrm{Var}(Q_c) &= \frac{1}{4}\sum_{i,j,k,l} H_{ij}H_{kl}\,\mathrm{Cov}(X_iX_j, X_kX_l)\\ &= \frac{1}{4}\sum_{i,j,k,l} H_{ij}H_{kl}(G_{ik}G_{jl} + G_{il}G_{jk})\\ &= \frac{1}{2}\,\mathrm{tr}[(\mathbf{H}\mathbf{G})^2]. \end{aligned} \tag{82}$$

The two sums in the middle line are equal because $\mathbf{H}$ and $\mathbf{G}$ are symmetric. Matrix multiplication and the trace provide a compact way to write that repeated sum. Similarly,

$$\mathrm{Cov}(L, Q_c) = \frac{1}{2}\sum_{i,j,k} b_i H_{jk}\mathbb{E}[X_iX_jX_k] = 0, \tag{83}$$

because every third-order moment of a centered multivariate normal is zero. Together with the usual variance of a weighted sum, these calculations give

$$V_{\text{lin}} = \boldsymbol{b}^{\mathsf{T}}\mathbf{G}\boldsymbol{b}, \qquad V_{\text{quad}} = \frac{1}{2}\operatorname{tr}[(\mathbf{HG})^2], \qquad \operatorname{Cov}(L, Q_c) = 0. \tag{84}$$

For a numerical example, take the values from Figure 3 of the paper:

$$\boldsymbol{b} = \begin{pmatrix} 0.75 \\ 0.30 \end{pmatrix}, \qquad \mathbf{G} = \begin{pmatrix} 1 & 0.3 \\ 0.3 & 0.6 \end{pmatrix}, \qquad \mathbf{H} = -0.8\mathbf{I}.$$

The linear variance includes the two trait variances and their covariance:

$$V_{\text{lin}} = (0.75)^2(1) + 2(0.75)(0.30)(0.3) + (0.30)^2(0.6) = 0.7515.$$

The middle term is the contribution from the association between the two breeding values. Because curvature here is $-0.8$ times the identity matrix, the quadratic variance is

$$V_{\text{quad}} = \frac{0.8^2}{2}\operatorname{tr}(\mathbf{G}^2) = 0.32\{1^2 + 0.6^2 + 2(0.3)^2\} = 0.4928.$$

In the paper's Figure 3, the axes give these two breeding values. Gray contours join combinations with equal log expected fitness. The blue and orange ellipses each contain 95% of the corresponding Gaussian population; they are density contours, not fitness contours. Under the Gaussian assumption the mean and covariance specify the complete distribution, so they determine how frequently each combination contributes to these variances. The first-order share is therefore $0.7515/(0.7515 + 0.4928) = 0.604$ after rounding. Multiplying every entry of $\mathbf{G}$ by 0.11 changes the two variances to $0.11(0.7515)$ and $0.11^2(0.4928)$, giving a share of 0.933. The same proportional-scaling result holds for any number of traits.

For completeness, write an affine predictor as $k + \boldsymbol{d}^{\mathsf{T}}\boldsymbol{X}$. A least-squares fit makes its residual uncorrelated with each predictor. These normal equations give

$$k = \ell_0 + \frac{1}{2}\operatorname{tr}(\mathbf{HG}), \qquad \mathbf{G}\boldsymbol{d} = \operatorname{Cov}(\boldsymbol{X}, \widetilde{\ell}_A) = \mathbf{G}\boldsymbol{b}. \tag{85}$$

The last equality follows because $\operatorname{Cov}(\boldsymbol{X}, Q_c) = \mathbf{0}$. Therefore $\boldsymbol{d} = \boldsymbol{b}$ on the positive-variance support of $\mathbf{G}$. The explained variance is $V_{\text{lin}}$ and the full second-order variance is $V_{\text{lin}} + V_{\text{quad}}$, so the population $R^2$ is $\mathcal{A}_g$. A conditional-mean-zero response residual adds the same residual variance to the linear and quadratic denominators, giving the normalized identity in Equation (35).

Under an invertible coordinate change $\boldsymbol{X}' = \mathbf{D}\boldsymbol{X}$,

$$\mathbf{G}' = \mathbf{D}\mathbf{G}\mathbf{D}^{\mathsf{T}}, \quad \boldsymbol{b}' = \mathbf{D}^{-\mathsf{T}}\boldsymbol{b}, \quad \mathbf{H}' = \mathbf{D}^{-\mathsf{T}}\mathbf{H}\mathbf{D}^{-1}. \tag{86}$$

Substitution leaves both quantities in Equation (84) unchanged. The one-trait unit conversion in Section 4 is the simplest instance of this result.

For the linear term, the cancellation can be seen directly:

$$(\boldsymbol{b}')^{\mathsf{T}}\mathbf{G}'\boldsymbol{b}' = \boldsymbol{b}^{\mathsf{T}}\mathbf{D}^{-1}(\mathbf{D}\mathbf{G}\mathbf{D}^{\mathsf{T}})\mathbf{D}^{-\mathsf{T}}\boldsymbol{b} = \boldsymbol{b}^{\mathsf{T}}\mathbf{G}\boldsymbol{b}. \tag{87}$$

For the quadratic term, $\mathbf{H}'\mathbf{G}' = \mathbf{D}^{-\mathsf{T}}\mathbf{H}\mathbf{G}\mathbf{D}^{\mathsf{T}}$. Multiplying two copies and cycling the factors inside the trace cancels $\mathbf{D}^{\mathsf{T}}\mathbf{D}^{-\mathsf{T}}$, leaving $\mathrm{tr}[(\mathbf{HG})^2]$. The transformation changes the numerical coordinates but leaves the variance components unchanged.

Whitening means changing coordinates so that breeding-value covariance becomes the identity matrix. Define $\widetilde{\mathbf{H}} = \mathbf{G}^{1/2}\mathbf{H}\mathbf{G}^{1/2}$ and let $\mu_i$ be its eigenvalues. Then

$$V_{\mathrm{lin}} = \left\|\mathbf{G}^{1/2}\boldsymbol{b}\right\|^2, \qquad V_{\mathrm{quad}} = \frac{1}{2}\sum_i \mu_i^2. \tag{88}$$

This expression also shows that $V_{\mathrm{quad}} \geq 0$ even though $\mathbf{HG}$ need not be symmetric. No alignment between the whitened gradient and the eigenvectors of $\widetilde{\mathbf{H}}$ enters $\mathcal{A}_g$. Two different orientation statements remain valid: alignment of $\boldsymbol{b}$ with high-variance directions of $\mathbf{G}$ changes $V_{\mathrm{lin}}$, and orientation of $\mathbf{H}$ relative to $\mathbf{G}$ changes $V_{\mathrm{quad}}$. For gradients of fixed ordinary length, $\boldsymbol{b}^{\mathsf{T}}\mathbf{G}\boldsymbol{b}$ is largest along the leading eigenvector of $\mathbf{G}$. To see this, write $\boldsymbol{b} = r\boldsymbol{u}$, where $r$ is fixed and $\|\boldsymbol{u}\| = 1$. Then

$$V_{\mathrm{lin}} = r^2\boldsymbol{u}^{\mathsf{T}}\mathbf{G}\boldsymbol{u} \leq r^2\lambda_{\max}(\mathbf{G}), \tag{89}$$

with equality when $\boldsymbol{u}$ is the eigenvector of the largest eigenvalue. This is the connection to the genetic line of least resistance [Schluter, 1996].

##### 9.3 Appendix S2 in several traits

Let $\boldsymbol{\Gamma} = -\mathbf{H}$ denote sign-reversed curvature. When $\boldsymbol{\Gamma} \succeq 0$, it represents stabilizing curvature; the result below also allows an indefinite $\boldsymbol{\Gamma}$, and therefore saddle-shaped curvature, if the stated integrability condition holds. Multiplying a normal breeding-value density by the exact log-quadratic selection weight gives

$$\mathbf{G}_{\mathrm{sel}} = (\mathbf{G}^{-1} + \boldsymbol{\Gamma})^{-1}, \qquad \boldsymbol{\Delta}_{\mathrm{full}} = \mathbf{G}_{\mathrm{sel}}\boldsymbol{b}, \tag{90}$$

provided $\mathbf{G}^{-1} + \boldsymbol{\Gamma}$ is positive definite on the support of $\mathbf{G}$. Here positive definite means that the combined precision coefficient is positive in every occupied direction, so the exponent bends downward sufficiently to define a finite weighted distribution. The covariance in Equation (90) is independent of $\boldsymbol{b}$ only within this exact closure.

Define the standardized gradient and curvature

$$\boldsymbol{c} = \mathbf{G}^{1/2}\boldsymbol{b}, \qquad \mathbf{T} = \mathbf{G}^{1/2}\boldsymbol{\Gamma}\mathbf{G}^{1/2}. \tag{91}$$

The first-order displacement is $\boldsymbol{\Delta}_{\mathrm{lin}} = \mathbf{G}\boldsymbol{b}$. Expressing both displacements in coordinates measured in units of breeding-value covariance gives

$$\mathbf{G}^{-1/2}\boldsymbol{\Delta}_{\mathrm{full}} = (\mathbf{I} + \mathbf{T})^{-1}\boldsymbol{c}, \tag{92}$$

and, after subtracting this expression from the first-order displacement,

$$\mathbf{G}^{-1/2}(\boldsymbol{\Delta}_{\text{lin}} - \boldsymbol{\Delta}_{\text{full}}) = \mathbf{T}(\mathbf{I} + \mathbf{T})^{-1}\boldsymbol{c}. \tag{93}$$

The first equation is the exact selected-parent displacement; the second is the part introduced by omitting curvature. Their Euclidean lengths in these coordinates are the $\mathbf{G}^{-1}$-norms in the next display.

When $\mathbf{G}^{1/2}\boldsymbol{b} \neq \mathbf{0}$, so the full displacement in the denominator is nonzero, Appendix S2 proves

$$\frac{\|\boldsymbol{\Delta}_{\text{lin}} - \boldsymbol{\Delta}_{\text{full}}\|_{\mathbf{G}^{-1}}}{\|\boldsymbol{\Delta}_{\text{full}}\|_{\mathbf{G}^{-1}}} \leq \|\mathbf{T}\|_2 \leq \|\mathbf{T}\|_{\text{F}} = \sqrt{2V_{\text{quad}}}. \tag{94}$$

If $\boldsymbol{c}$ is an eigenvector of $\mathbf{T}$ with eigenvalue $\tau_k$, the relative error is exactly $|\tau_k|$. It equals the spectral-norm bound only when $|\tau_k|$ is the largest absolute eigenvalue. Thus the gradient's direction can affect the selected-parent displacement even though its alignment with curvature eigenvectors does not enter the variance share.

##### 9.4 Appendix S3 in several traits

With fixed allelic effects $\boldsymbol{\alpha}_j$, the genic covariance that would be present without linkage disequilibrium is

$$\mathbf{B}_t = 2\sum_j p_{j,t}(1 - p_{j,t})\boldsymbol{\alpha}_j\boldsymbol{\alpha}_j^{\mathsf{T}}. \tag{95}$$

Here $p_{j,t}$ is the allele frequency at locus $j$ in generation $t$, and $\boldsymbol{\alpha}_j\boldsymbol{\alpha}_j^{\mathsf{T}}$ is the table obtained by multiplying each trait effect at that locus by every trait effect. Fixed allelic effects do not make $\mathbf{B}_t$ fixed: allele-frequency change can change it. Replacing $\mathbf{B}_t$ by a constant $\mathbf{B}$ is therefore a separate short-timescale closure [Walsh and Lynch, 2018].

The covariance update is

$$F(\mathbf{G}) = (1 - \rho)\mathbf{B} + \rho(\mathbf{G}^{-1} + \boldsymbol{\Gamma})^{-1}. \tag{96}$$

For $\mathbf{B}$ positive definite, $\boldsymbol{\Gamma} \succeq 0$, and $0 \leq \rho < 1$, stabilizing selection gives

$$(\mathbf{G}^{-1} + \boldsymbol{\Gamma})^{-1} \preceq \mathbf{G}. \tag{97}$$

Write $S(\mathbf{G}) = (\mathbf{G}^{-1} + \boldsymbol{\Gamma})^{-1}$. For any covariance matrix in the interval

$$(1 - \rho)\mathbf{B} \preceq \mathbf{G} \preceq \mathbf{B} \tag{98}$$

the selection inequality $\mathbf{0} \prec S(\mathbf{G}) \preceq \mathbf{G}$ gives both sides of the next calculation:

$$(1 - \rho)\mathbf{B} \preceq F(\mathbf{G}) \preceq (1 - \rho)\mathbf{B} + \rho\mathbf{G} \preceq \mathbf{B}. \tag{99}$$

Thus the update maps the whole interval back into itself. This matrix interval is nonempty, closed, bounded, and convex; convex means that an average of two members is also a member. The update is continuous. Brouwer's fixed-point theorem states that a continuous map from such a finite-dimensional set into itself has at least one fixed point. The theorem establishes existence here; it does not establish uniqueness or convergence.

The bounds apply to every positive-definite fixed point, even when a trajectory starts outside the interval. The lower bound follows because the second term of $F$ is nonnegative. For the upper bound, substitute a fixed point into the selection inequality:

$$\mathbf{G}^* = F(\mathbf{G}^*) \preceq (1-\rho)\mathbf{B} + \rho\mathbf{G}^*. \tag{100}$$

Subtracting $\rho\mathbf{G}^*$ and dividing by the positive number $1-\rho$ gives $\mathbf{G}^* \preceq \mathbf{B}$. Combining the two arguments gives

$$(1-\rho)\mathbf{B} \preceq \mathbf{G}^* \preceq \mathbf{B}. \tag{101}$$

Uniqueness and convergence of the scalar recursion in Equation (49) follow from the contraction inequality in Equation (61).

At $\rho = 1$, for positive-definite $\mathbf{G}_0$ or with every inverse restricted to one fixed positive-variance support,

$$\mathbf{G}_{t+1}^{-1} = \mathbf{G}_t^{-1} + \boldsymbol{\Gamma}, \qquad \mathbf{G}_t^{-1} = \mathbf{G}_0^{-1} + t\boldsymbol{\Gamma}. \tag{102}$$

Strictly positive stabilizing curvature removes covariance in every direction. If curvature is zero in some directions, covariance can remain in that null space, meaning the set of directions that $\boldsymbol{\Gamma}$ maps to zero.

#### 9.5 Appendix S5 in several traits

Let $\boldsymbol{x}$ be centered phenotype, $\mathbf{P}$ its covariance matrix, $\boldsymbol{\beta}$ the vector of linear coefficients, and $\boldsymbol{\gamma}$ the symmetric matrix of quadratic coefficients in

$$w = \alpha + \boldsymbol{\beta}^\mathsf{T}\boldsymbol{x} + \frac{1}{2}\boldsymbol{x}^\mathsf{T}\boldsymbol{\gamma}\boldsymbol{x} + \varepsilon. \tag{103}$$

Gaussian moments give

$$\alpha = 1 - \frac{1}{2}\operatorname{tr}(\boldsymbol{\gamma}\mathbf{P}), \quad \mathbb{E}[\boldsymbol{x}w] = \mathbf{P}\boldsymbol{\beta}, \quad \mathbb{E}[\boldsymbol{x}\boldsymbol{x}^\mathsf{T}w] = \mathbf{P} + \mathbf{P}\boldsymbol{\gamma}\mathbf{P}. \tag{104}$$

If centered breeding value $\boldsymbol{A}$ and centered phenotype $\boldsymbol{x}$ are jointly Gaussian in compatible coordinates, if $\operatorname{Cov}(\boldsymbol{A}, \boldsymbol{x}) = \mathbf{G}$, if $\mathbf{P}$ is positive definite on the occupied support, and if selection is mediated by the measured phenotypes, then

$$\mathbb{E}[\boldsymbol{A} \mid \boldsymbol{x}] = \mathbf{G}\mathbf{P}^{-1}\boldsymbol{x}, \qquad \operatorname{Var}(\boldsymbol{A} \mid \boldsymbol{x}) = \mathbf{G} - \mathbf{G}\mathbf{P}^{-1}\mathbf{G}. \tag{105}$$

Average the first expression using the selection weights and substitute $\mathbb{E}[\boldsymbol{x}w] = \mathbf{P}\boldsymbol{\beta}$. This gives

$$\mathbb{E}[\boldsymbol{A} \mid \mathrm{sel}] - \mathbb{E}[\boldsymbol{A}] = \mathbf{G}\boldsymbol{\beta}, \tag{106}$$

For the second moment, add the conditional covariance in Equation (105) to the outer product of its conditional mean, average, and substitute $\mathbb{E}[\boldsymbol{x}\boldsymbol{x}^{\mathsf{T}}w] = \mathbf{P} + \mathbf{P}\boldsymbol{\gamma}\mathbf{P}$. The resulting selected second moment is $\mathbf{G} + \mathbf{G}\boldsymbol{\gamma}\mathbf{G}$. Subtracting the outer product of the selected mean gives

$$\mathbf{G}_{\mathrm{sel}} = \mathbf{G} + \mathbf{G}(\boldsymbol{\gamma} - \boldsymbol{\beta}\boldsymbol{\beta}^{\mathsf{T}})\mathbf{G}. \tag{107}$$

Matching these moments to Equation (90) yields

$$\boldsymbol{b} = \mathbf{G}_{\mathrm{sel}}^{-1}\mathbf{G}\boldsymbol{\beta}, \qquad \mathbf{H} = \mathbf{G}^{-1} - \mathbf{G}_{\mathrm{sel}}^{-1}. \tag{108}$$

For comparison, one could instead differentiate the fitted raw-scale phenotype regression itself. At the centered phenotype $\boldsymbol{x} = \mathbf{0}$ and provided $\alpha > 0$, that separate calculation gives

$$\nabla \log \widehat{w}(\mathbf{0}) = \frac{\boldsymbol{\beta}}{\alpha}, \qquad \nabla^2 \log \widehat{w}(\mathbf{0}) = \frac{\boldsymbol{\gamma}}{\alpha} - \frac{\boldsymbol{\beta}\boldsymbol{\beta}^{\mathsf{T}}}{\alpha^2}. \tag{109}$$

These are derivatives of the fitted phenotype-space projection. They are not generally the breeding-value-space $\boldsymbol{b}$ and $\mathbf{H}$ above. This is why Appendix S5 matches selected breeding-value moments rather than equating the two sets of derivatives.

These formulas require the conditions listed in Section 8. If $\mathbf{G}$ or $\mathbf{P}$ is singular, inverses and calculations are restricted to the relevant positive-variance support.

### 10 Answers to the checks

#### Answer 1

The first conditional mean is $(0+2)/2 = 1$, so its logarithm is 0. The second mean is $(1+3)/2 = 2$, so its logarithm is $\log 2 \simeq 0.693$. The mean of the logged values is undefined in the first group because $\log 0$ is not finite. This demonstrates why averaging and taking the logarithm cannot generally be reversed.

#### Answer 2

The slope calculation is

$$\frac{0.39 - (-0.41)}{2(0.2)} = \frac{0.80}{0.40} = 2. \tag{110}$$

Because $\ell_A(0) - \ell_0 = 0$, the curvature calculation is

$$\frac{0.39 - 2(0) - 0.41}{0.2^2} = \frac{-0.02}{0.04} = -0.5. \tag{111}$$

#### Answer 3

$V_{\text{lin}} = b^2G = 4$ is variance in the first-order term over the current breeding-value distribution. $V_{\text{quad}} = H^2G^2/2 = 0.125$ is variance in the centered quadratic term. Their share is $\mathcal{A}_g = 4/(4+0.125) = 0.9697$. It is a share of variance in the second-order approximation to log conditional expected fitness, not of realized-fitness variance.

#### Answer 4

At $x = 2$, the product is $4(-0.75) = -3$; at $x = -2$, it is $(-4)(-0.75) = 3$. A normal distribution centered at zero assigns equal density to $x$ and $-x$, so these contributions cancel. The same pairing holds for every positive and negative value.

#### Answer 5

Multiplying breeding values by 10 multiplies variance by $10^2$, so $G' = 100$. A one-unit change on the new scale is one tenth of a unit on the old scale, so $b' = b/10 = 0.2$ and $H' = H/10^2 = -0.005$. The transformed variance terms are 4 and 0.125, giving the same value 0.9697 for $\mathcal{A}_g$.

#### Answer 6

$$g_{\text{sel}} = \frac{1}{1/1 + 0.5} = \frac{2}{3}, \qquad \Delta\bar{a}_{\text{full}} = \frac{2}{3}(2) = \frac{4}{3}. \tag{112}$$

These describe selected contributors, not offspring.

#### Answer 7

With $b = 10$, $V_{\text{lin}} = 100$ and $V_{\text{quad}} = 0.125$, so

$$\mathcal{A}_g = \frac{100}{100.125} = 0.9988. \tag{113}$$

The first-order displacement is 10 and the full displacement is $10/1.5 = 6.6667$. Their relative discrepancy is $|10 - 6.6667|/6.6667 = 0.5$. Increasing $b$ enlarges the numerator of $\mathcal{A}_g$, but the relative displacement discrepancy in one trait depends on $\Gamma g$, which has not changed.

#### Answer 8

$$g_1 = 0.5 + 0.5\frac{1}{1 + 0.5} = 0.8333, \tag{114}$$
$$g_2 = 0.5 + 0.5\frac{0.8333}{1 + 0.5(0.8333)} = 0.7941, \tag{115}$$
$$g_3 = 0.5 + 0.5\frac{0.7941}{1 + 0.5(0.7941)} = 0.7842. \tag{116}$$

All three values lie between 0.5 and 1.

#### Answer 9

Equation (64) gives $1/g_4 = 1/g_0 + 4\gamma = 1 + 2 = 3$, so $g_4 = 1/3$. When $\rho = 1$, the multiplier $1 - \rho$ is zero, so the baseline contribution containing $B$ is absent.

#### Answer 10

No. Agreement for the current-state share checks the variance or projection calculation under the simulated distribution. The fixed point belongs to a separate across-generation model with fixed $B$, fixed curvature, and a specified retention rule. A simulation that does not implement that update cannot test its fixed point.

#### Answer 11

$$\alpha = 1 - \tfrac{1}{2}(-0.2)(1.5) = 1.15, \tag{117}$$

$$G_{\text{sel}} = 0.5 + (0.5)^2\{-0.2 - (0.4)^2\} = 0.41, \tag{118}$$

$$b = 0.5(0.4)/0.41 = 0.4878, \tag{119}$$

$$H = 1/0.5 - 1/0.41 = -0.4390. \tag{120}$$

$\alpha$ is the fitted value at mean phenotype. The normalization $\mathbb{E}[w] = 1$ fixes the fitted average over the whole phenotype distribution, not the fitted value at its mean.

### 11 Glossary

| Term or symbol | Meaning in this guide |
|---|---|
| $A$, $a$ | A breeding-value variable and one possible value of it. |
| $E$ | Environmental deviation in the recurring example. |
| $Z$ | Phenotype, with $Z = A + E$ in the recurring example. |
| $W$ | Realized reproductive output in one specified selection episode. In the recurring example, $W(Z)$ is the deterministic selection weight assigned to phenotype $Z$. |
| $\bar{W}$, $w$ | Population mean fitness and relative fitness $w = W/\bar{W}$. |
| $W_A(a)$ | Mean realized reproductive output conditional on breeding value $a$. |
| $\ell_A(a)$ | $\log W_A(a)$, the logarithm of conditional expected fitness. |
| $p_A(a)$, $p_{\text{sel}}(a)$ | Breeding-value distributions before and after weighting by conditional expected fitness. |
| $X$ | Random breeding value centered on the current population mean. |
| $x$ | A possible value of centered breeding value in Appendix S1; centered phenotype in Appendix S5. |
| $Y$ | Generic observed outcome in the nested-regression identity. |
| $G$, $g$ | Breeding-value variance; bold $\mathbf{G}$ denotes the covariance matrix for several traits. |
| $P$, $\mathbf{P}$ | Phenotypic variance; bold $\mathbf{P}$ denotes the phenotypic covariance matrix for several traits. |
| $b$, $\boldsymbol{b}$ | Local slope in one trait; vector of slopes in several traits. |
| $H$, $\mathbf{H}$ | Local curvature in one trait; curvature matrix in several traits. |
| $L$ | First-order contribution to the local description of the logarithm of conditional expected fitness. |
| $Q_c$ | Quadratic contribution after subtracting its population mean. |
| $V_{\text{lin}}$ | Variance of $L$ over the current breeding-value distribution. |
| $V_{\text{quad}}$ | Variance of $Q_c$ over the current breeding-value distribution. |
| $\mathcal{A}_g$ | First-order variance share $V_{\text{lin}}/(V_{\text{lin}} + V_{\text{quad}})$. |

| Term or symbol | Meaning in this guide |
|---|---|
| $\alpha$ | Intercept in the phenotype-space relative-fitness regression. |
| $\beta$, $\boldsymbol{\beta}$ | Linear coefficient in that phenotype-space regression. |
| $\gamma$, $\boldsymbol{\gamma}$ | Context-dependent notation. In Appendix S3, scalar $\gamma$ is stabilizing curvature in the one-trait recursion. In Appendix S5, $\gamma$ and matrix $\boldsymbol{\gamma}$ are conventional quadratic coefficients in the phenotype-space regression. |
| $\varepsilon$ | Regression residual: the part not represented by the fitted constant, linear, and quadratic terms. |
| $\boldsymbol{\alpha}_j$ | Fixed vector of allelic effects at locus $j$ in the across-generation model; this is distinct from the regression intercept $\alpha$. |
| $\Gamma$, $\mathbf{\Gamma}$ | Sign-reversed curvature, defined as $-H$ or $-\mathbf{H}$; it represents stabilizing curvature when it is nonnegative in one trait or positive semidefinite in several traits. |
| $g_{\text{sel}}$, $G_{\text{sel}}$, $\mathbf{G}_{\text{sel}}$ | Breeding-value variance or covariance among selected contributors. |
| $\Delta\bar{a}_{\text{lin}}$, $\Delta\bar{a}_{\text{full}}$, $\mathbf{\Delta}_{\text{lin}}$, $\mathbf{\Delta}_{\text{full}}$ | Selected-parent mean displacements from the first-order model and the full log-quadratic model, in one or several traits. |
| $d$, $\boldsymbol{d}$, $k$ | Fitted slope, fitted slope vector, and intercept in the least-squares projection. The paper calls the intercept $c$; the guide uses $k$ to distinguish it from $\boldsymbol{c}$. |
| $\boldsymbol{c}$, $\mathbf{T}$ | Slope and sign-reversed curvature after rescaling trait coordinates by the breeding-value covariance. |
| $p$, $u$ | Allele frequency and allelic effect in the one-locus illustration. |
| $B$, $\mathbf{B}$ | Fixed genic variance or covariance baseline in the across-generation closure. |
| $\rho$ | Scalar retention of the postselection covariance departure from the baseline. |
| $\mathbb{E}$ | Population average or expectation. |
| Var | Variance: average squared distance from the mean. |
| Cov | Covariance: average product of two centered quantities. |
| Corr | Correlation: covariance divided by the two standard deviations. |
| $R^2$ | Fraction of outcome variance accounted for by a fitted regression. |

### References

Nicholas H. Barton and Michael Turelli. Natural and sexual selection on many loci. *Genetics*, 127(1):229–255, 1991. doi: 10.1093/genetics/127.1.229.

M. G. Bulmer. Linkage disequilibrium and genetic variability. *Genetical Research*, 23(3):281–289, 1974. doi: 10.1017/S0016672300014920.

Michael G. Bulmer. The effect of selection on genetic variability. *The American Naturalist*, 105(943):201–211, 1971. doi: 10.1086/282718.

Pierre de Villemereuil, Holger Schielzeth, Shinichi Nakagawa, and Michael B. Morrissey. General methods for evolutionary quantitative genetic inference from generalized mixed models. *Genetics*, 204(3):1281–1294, 2016. doi: 10.1534/genetics.115.186536.

R. A. Fisher. Average excess and average effect of a gene substitution. *Annals of Eugenics*, 11(1):53–63, 1941. doi: 10.1111/j.1469-1809.1941.tb02272.x.

Ronald A. Fisher. The correlation between relatives on the supposition of Mendelian inheritance. *Transactions of the Royal Society of Edinburgh*, 52(2):399–433, 1918. doi: 10.1017/S0080456800012163.

Jarrod D. Hadfield, Alastair J. Wilson, Dany Garant, Ben C. Sheldon, and Loeske E. B. Kruuk. The misuse of BLUP in ecology and evolution. *The American Naturalist*, 175(1):116–125, 2010. doi: 10.1086/648604.

Benjamin C. Haller, Peter L. Ralph, and Philipp W. Messer. SLiM 5: Eco-evolutionary simulations across multiple chromosomes and full genomes. *Molecular Biology and Evolution*, 43(1):msaf313, 2026. doi: 10.1093/molbev/msaf313.

L. Isserlis. On a formula for the product-moment coefficient of any order of a normal frequency distribution in any number of variables. *Biometrika*, 12(1–2):134–139, 1918. doi: 10.1093/biomet/12.1-2.134.

Russell Lande. Quantitative genetic analysis of multivariate evolution, applied to brain:body size allometry. *Evolution*, 33(1Part2):402–416, 1979. doi: 10.1111/j.1558-5646.1979.tb04694.x.

Russell Lande and Stevan J. Arnold. The measurement of selection on correlated characters. *Evolution*, 37(6): 1210–1226, 1983. doi: 10.1111/j.1558-5646.1983.tb00236.x.

Jacob A. Moorad and Michael J. Wade. Selection gradients, the opportunity for selection, and the coefficient of determination. *The American Naturalist*, 181(3):291–300, 2013. doi: 10.1086/669158.

Michael B. Morrissey and I. B. J. Goudie. Analytical results for directional and quadratic selection gradients for log-linear models of fitness functions. *Evolution*, 76(7):1378–1390, 2022. doi: 10.1111/evo.14486.

Dolph Schluter. Adaptive radiation along genetic lines of least resistance. *Evolution*, 50(5):1766–1774, 1996. doi: 10.1111/j.1558-5646.1996.tb03563.x.

John R. Stinchcombe, Aneil F. Agrawal, Paul A. Hohenlohe, Stevan J. Arnold, and Mark W. Blows. Estimating nonlinear selection gradients using quadratic regression coefficients: Double or nothing? *Evolution*, 62(9): 2435–2440, 2008. doi: 10.1111/j.1558-5646.2008.00449.x.

Bruce Walsh and Michael Lynch. *Evolution and Selection of Quantitative Traits*. Oxford University Press, Oxford, 2018. ISBN 9780198830870. doi: 10.1093/oso/9780198830870.001.0001. Online ISBN 9780191868986.
