## Supplementary material for "Additive Channels in Curved Fitness Landscapes": Supp Material 2

### Supplementary File 3
### Estimating the variance share from simulated observations

Daniel Ortiz-Barrientos and Mark Cooper

Suppose taller plants tend to leave more offspring, but the advantage weakens and eventually reverses. How much of the variation in log expected fitness can a straight line still describe? This example estimates that share from noisy height measurements and offspring counts. It follows the five steps in the manuscript's section *Estimating the variance share*: define the observations, fit expected fitness, obtain slope and curvature, calculate the variance components, and assess uncertainty and model adequacy. The accompanying Shiny app uses exactly the same analysis code.

We simulate one population during one reproductive episode, so we know the true share before we fit the model. Comparing the estimate with that value checks the estimation procedure. It does not test inheritance or predict an offspring mean.

#### 1 Step 1: define the population, trait and reproductive output

Consider 800 unrelated plant genotypes. In this model, a breeding value is the additive genetic contribution to height. Each genotype's value $A_i$ is expressed in centimeters relative to the population mean. To keep the example identifiable, assume that the genetic contribution to height is wholly additive. Three independent clonal replicates of each genotype provide height deviations $Y_{ij}$:

$$A_i \sim N(0, G), \qquad G = 1\ \text{cm}^2, \tag{1}$$

$$Y_{ij} = A_i + e_{ij}, \qquad e_{ij} \sim N(0, E), \quad E = 1\ \text{cm}^2. \tag{2}$$

The environmental deviations are independent across replicates and genotypes. There are no permanent environmental effects, dominance or epistasis. These are assumptions of this teaching model. In real clonal data, between-genotype variance can contain nonadditive components; clonal replication alone does not identify additive variance. Here that identification follows from the assumed additive trait model.

A separate reproductive assay provides one count $W_i$ of surviving offspring per genotype. Conditional on $A_i$, that count is independent of the environments used to measure height:

$$W_i \mid A_i = a \sim \text{Poisson}\{W_A(a)\}, \tag{3}$$

$$\log W_A(a) = \alpha + ba + \tfrac{1}{2}Ha^2, \qquad \alpha = \log 4, \quad b = 0.7\ \text{cm}^{-1}, \quad H = -0.7\ \text{cm}^{-2}. \tag{4}$$

The value 4 is expected output at the mean breeding value; it is not the population mean output. The negative curvature allows expected output to increase and then decrease across breeding values. We simulate the conditional relationship directly. We do not construct it from a separate phenotype–fitness function or simulate an inheritance process.

The seed is 20261005. The saved observations contain genotype identifiers, three height measurements and offspring counts. The true breeding values are stored in a separate file and are never passed to the estimator. The default dataset contains 88 zero counts; each is retained in the likelihood.

Before fitting anything, calculate the answer implied by the generating model:

$$V_{\text{lin}} = b^2 G = 0.7^2 \times 1 = 0.49, \tag{5}$$

$$V_{\text{quad}} = \tfrac{1}{2} H^2 G^2 = \tfrac{1}{2} \times 0.7^2 \times 1^2 = 0.245, \tag{6}$$

$$\mathcal{A}_g = \frac{0.49}{0.49 + 0.245} = \frac{2}{3}. \tag{7}$$

This is a share of variation in log *expected* fitness across breeding values. Poisson variation in individual offspring counts is not included in this denominator.

### 2 Step 2: fit the conditional mean while allowing for uncertain breeding values

#### 2.1 Use replication to estimate measurement uncertainty

Let $m = 3$, let $\bar{Y}_i$ be the mean height deviation for genotype $i$, and let $\bar{Y}$ be the mean across genotypes. First estimate environmental variance from variation within genotypes:

$$\widehat{E} = \frac{\sum_{i=1}^{n} \sum_{j=1}^{m} (Y_{ij} - \bar{Y}_i)^2}{n(m-1)}. \tag{8}$$

Averaging $m$ independent measurements reduces environmental variance to $E/m$. Because $\mathrm{Var}(\bar{Y}_i) = G + E/m$, the corresponding estimate of genetic variance is

$$\widehat{G} = \frac{\sum_i (\bar{Y}_i - \bar{Y})^2}{n-1} - \frac{\widehat{E}}{m}. \tag{9}$$

The default data give $\widehat{E} = 0.9799$ and $\widehat{G} = 1.0021$. A nonpositive estimate of $G$ stops this analysis: the code does not replace it silently with a small positive number.

Let $\mu$ denote the population mean, which is zero in the generating model and estimated by $\widehat{\mu} = \bar{Y}$ when fitting the observations. Under the normal measurement model, a genotype's centered breeding value $Z_i = A_i - \mu$ is normally distributed conditional on its height measurements. Substituting the calibrated parameters gives

$$r = \frac{\widehat{G}}{\widehat{G} + \widehat{E}/m}, \qquad M_i = r(\bar{Y}_i - \bar{Y}), \tag{10}$$

$$Z_i \mid Y_{i1}, \ldots, Y_{im} \sim N(M_i, v), \qquad v = \widehat{G}(1 - r). \tag{11}$$

Here $r = 0.7542$. The quantity $M_i$ is a prediction of the centered breeding value; $v$ expresses the uncertainty that remains. Replacing $Z_i$ by $M_i$ and discarding $v$ would treat a prediction as an observation. Problems with using estimated breeding values in subsequent regressions are discussed by Hadfield et al. (2010).

#### 2.2 Fit counts, without taking their logarithms

We now estimate $\alpha, b, H$ from reproductive counts. For each genotype, average the Poisson probability over the possible breeding values in Equation (11):

$$L_i(\alpha, b, H) = \int \frac{\lambda(z)^{W_i} e^{-\lambda(z)}}{W_i!} \phi(z; M_i, v)\, dz, \qquad \lambda(z) = \exp(\alpha + bz + Hz^2/2). \tag{12}$$

The symbol $\phi$ denotes a normal density. Each possible breeding value contributes according to its probability given the height measurements. We maximize $\sum_i \log L_i$. This takes logarithms of probabilities, not of offspring counts, so observations with $W_i = 0$ present no difficulty. The use of a log link for expected fitness follows the distinction between response and link scales discussed by Morrissey and Goudie (2022).

The implementation uses normal quadrature: it evaluates the integral at weighted points representing the normal distribution. The default fit uses 41 points. Repeating it with 81 points changes $b$, $H$ and $\mathcal{A}_g$ by less than $4 \times 10^{-13}$ in this dataset. For other settings and every bootstrap sample, the code increases the quadrature order and checks that the fitted coefficients and likelihood stabilize. It stops rather than reports an unresolved integral. Two starting values are used for optimization. The primary fit restricts $H \leq 0$, matching the stabilizing family used here; this example is not designed to fit disruptive selection. Numerical parameter bounds are reported in the code, and a fit at a bound is flagged.

This is a *two-stage estimator*. The height measurements determine $\widehat{G}$, $\widehat{E}$ and the conditional normal distributions; the counts then determine the fitness parameters. We do not claim that it is the most efficient estimator or that it replaces a joint pedigree or genomic analysis. The bootstrap below repeats both stages.

Figure 1 distinguishes the relationship fitted to noisy height measurements from the relationship estimated over breeding values. Averaging over uncertainty is needed for both the likelihood and predictions at observed measurements.

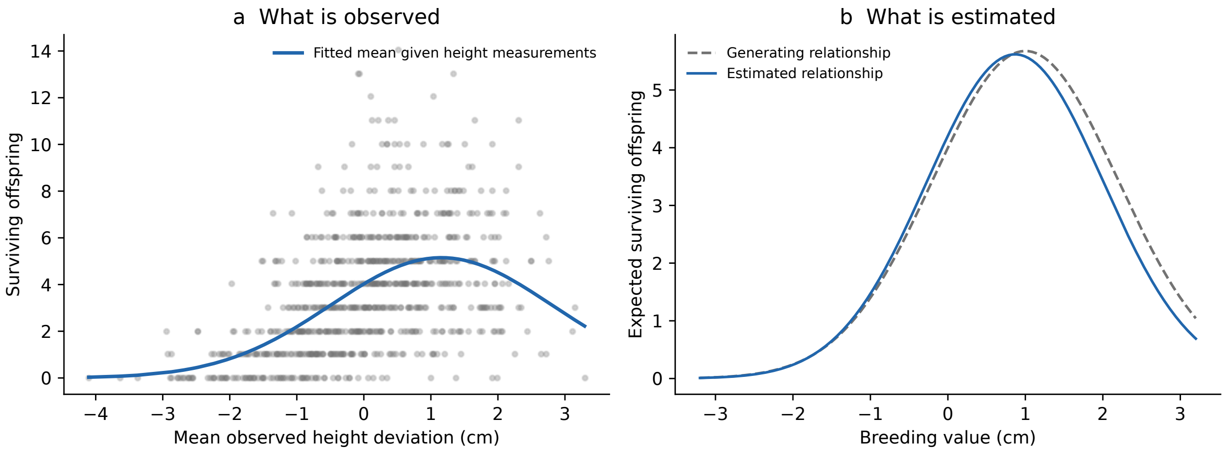


Figure 1: Observed counts and the estimated conditional fitness relationship. (a) Each point gives the mean of three height measurements and an offspring count. The curve averages expected output over the uncertain breeding value associated with those measurements. (b) The fitted breeding-value relationship is compared with the generating relationship. The latter is available only because this is a simulation.

#### 3 Steps 3 and 4: obtain slope, curvature and the variance share

The fitted log mean is $\widehat{\ell}_A(z) = \widehat{\alpha} + \widehat{b}z + \widehat{H}z^2/2$. Differentiating gives

$$\frac{d\widehat{\ell}_A}{dz} = \widehat{b} + \widehat{H}z, \qquad \frac{d^2\widehat{\ell}_A}{dz^2} = \widehat{H}. \tag{13}$$

At the estimated population mean, $z = 0$, so the fitted coefficients give the local slope and curvature directly. The factor $1/2$ in the design matrix matters: if software instead fits a coefficient $c$ multiplying $z^2$, then $H = 2c$.

Insert these coefficients and $\widehat{G}$ from Equation (9) into Equations (5)–(7). Table 1 reports all three inputs and all three outputs. The point estimate is $\widehat{\mathcal{A}_g} = 0.6107$, compared with the generating value $2/3$. A single dataset need not recover the generating parameters exactly.

| Quantity | Generating value | Estimate | 95% lower | 95% upper |
|---|---|---|---|---|
| $b$ | 0.7000 | 0.6826 | 0.6003 | 0.7789 |
| $H$ | -0.7000 | -0.7698 | -0.9167 | -0.6301 |
| $G$ | 1.0000 | 1.0021 | 0.8766 | 1.1363 |
| $V_{\text{lin}}$ | 0.4900 | 0.4669 | 0.3547 | 0.5996 |
| $V_{\text{quad}}$ | 0.2450 | 0.2976 | 0.1956 | 0.4212 |
| $\mathcal{A}_g$ | 0.6667 | 0.6107 | 0.5330 | 0.6857 |

Table 1: Generating values and estimates for seed 20261005. Intervals are 95% percentile intervals from 500 parametric bootstrap samples, conditional on the fitted quadratic model. Slope and curvature are evaluated at the estimated population mean, whose sampling uncertainty is included by recalibration in each bootstrap sample.

For comparison, the code also fits the shrunken predictions $M_i$ as exact covariates and substitutes their empirical variance for $G$. This naive analysis gives $\widehat{\mathcal{A}_g} = 0.6740$. It is closer to 2/3 in this particular sample, although both absolute variance components are underestimated. Closeness of a ratio in one dataset does not validate the procedure. Because this comparison re-estimates slope and curvature as well as changing the variance, it does not test the manuscript's narrower bias formula, which holds slope and curvature fixed.

Figure 2 shows why the two components should accompany their ratio. Their units here are squared natural-log fitness differences. They are not additive and epistatic variance components of height, nor measures of the accuracy of predicting offspring change.

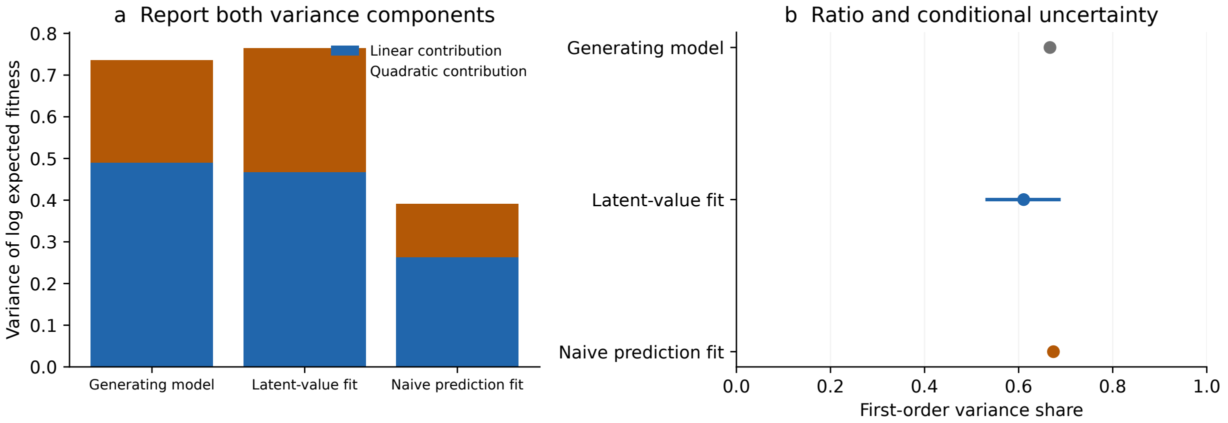


Figure 2: Variance components and their ratio. (a) The generating, latent-value and naive analyses can yield similar ratios while differing in the amount of variation assigned to each term. (b) The interval applies to the latent-value estimate under the fitted observation model. The generating value has no estimation interval because it is fixed by the simulation parameters.

### 4 Step 5: quantify uncertainty and check the model

#### 4.1 Repeat both stages of estimation

We use a parametric bootstrap with seed 20261006. For each replicate:

1. Draw new breeding values from $N(0, \widehat{G})$.
2. Draw three noisy height measurements per genotype using $\widehat{E}$ and the estimated population mean.
3. Draw new offspring counts from the fitted quadratic Poisson model.

4. Re-estimate the measurement mean, $G$ and $E$; refit the fitness relationship; recalculate the two variance components and their share.

The 2.5th and 97.5th percentiles of the estimates give the intervals in Table 1. All 500 default replicates completed; there were 0 failed fits and no primary fits at a parameter bound. The complete replicate estimates are supplied as a CSV file. All replicates completing tells us the fitting procedure ran. To measure coverage we would need to simulate many independent datasets and count how often the intervals contained the true share.

These intervals account for sampling variation and uncertainty in the breeding values and calibrated variances under the assumed model. They do not include uncertainty about whether the biological model is correct. We examine that separately. The app allows 20–1000 replicates; 20 is useful only for seeing how the calculation works. Increasing the number of bootstrap replicates reduces Monte Carlo variation in interval endpoints; it does not remove model bias.

#### 4.2 Compare fitted and observed quantities on the same scale

Group genotypes by their observed mean height. Within each group, compare mean offspring output with predictions that integrate over uncertain breeding values. Also compare the observed fraction of zeros with the fitted Poisson-mixture probability of zero. Figure 3 illustrates these checks. Plotting counts against predicted breeding values and overlaying $W_A$ directly would compare different conditional means.

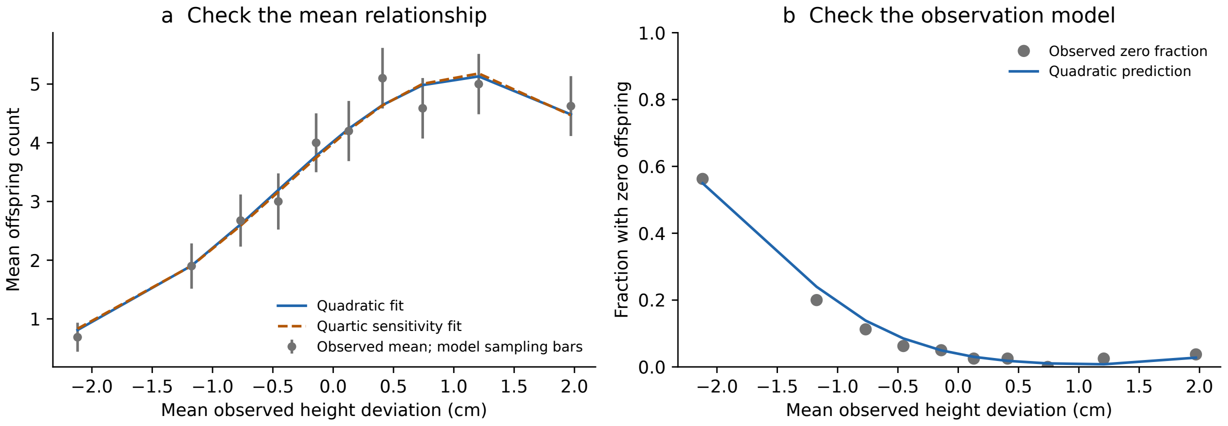


Figure 3: Checks based on observable data. (a) Observed bin means are compared with quadratic and quartic conditional predictions. Bars show approximate 95% sampling ranges around the observed means, using the sum of fitted individual predictive variances within each bin. They condition on fitted parameters and are neither simultaneous bands nor confidence intervals for the fitness surface. (b) Observed and predicted zero fractions provide a further check of the count model.

#### 4.3 Examine the quadratic assumption explicitly

The app can add a term $-\kappa a^4$ to the generating log mean, with $\kappa = 0.08$. The primary analysis still fits a quadratic. A second fit includes cubic and quartic terms, allowing for an estimated population center. Comparing the two fits shows how sensitive the result is to the assumed shape of the fitness curve. For the default quadratic data, quartic minus quadratic AIC is 3.4109; a negative difference would favor the more flexible fit. The two fits share the measurement calibration. This approximate conditional-likelihood comparison is not a formal test of the observation model, and numerical boundaries can affect its interpretation.

The fourth-order scenario demonstrates a distinction that matters for the manuscript. At $a = 0$, the added term changes neither slope nor curvature, so the generating *local quadratic share* remains 2/3. Nevertheless, it changes variation over the population. For a centered Gaussian breeding value,

$$\mathrm{Var}\{\tfrac{1}{2}HA^2 - \kappa A^4\} = \tfrac{1}{2}H^2G^2 - 12H\kappa G^3 + 96\kappa^2G^4. \tag{14}$$

This follows from $\mathrm{Var}(A^2) = 2G^2$, $\mathrm{Cov}(A^2, A^4) = 12G^3$ and $\mathrm{Var}(A^4) = 96G^4$. The linear term remains uncorrelated with these even terms. Therefore the affine $R^2$ for the complete generating log mean is

$$R^2_{\text{full}} = \frac{b^2G}{b^2G + \frac{1}{2}H^2G^2 - 12H\kappa G^3 + 96\kappa^2G^4}. \tag{15}$$

At the stated parameters this is about 0.2424, despite the local quadratic share of 0.6667. A fitted quadratic can consequently summarize the wrong amount of variation when higher-order terms matter. Its bootstrap interval does not repair that mismatch. Nor is the local quadratic share from the quartic fit the $R^2$ of the complete quartic curve.

#### 5 Using the files and app

From the repository root, run `bash run_estimation.sh` to regenerate the default dataset, estimates, 500 bootstrap samples and three figures. Run `bash run_shiny.sh` to open the companion app locally. Both commands create a separate Python environment when needed. The first installation requires an internet connection; later runs use the installed dependencies. No R installation or SLiM run is required.

The app follows the same sequence as this document. Its settings control population size, replication, genetic and environmental variance, slope, curvature, output at the mean and the random seed. Press *Generate and estimate* after changing them. Bootstrap calculations use a separate button. Generating a new dataset clears its previous intervals. The download contains the displayed data, simulation truth, parameter settings, estimates and any completed bootstrap samples.

The folder `examples/estimating_variance_share/` contains annotated Python code, this document's source and PDF, the app and the default results. `model.py` implements the simulation and estimator; `run_case.py` produces the archived example; `test_model.py` checks the calculations against Gaussian moments, a closed-form expected output, independent numerical integration and repeated simulations. The Shiny app calls these same functions. The numerical example is deterministic for the recorded seed and software versions in `results/summary.json`.

For real data, the measurement model must reflect the design. Related individuals need a relationship model; heterogeneous prediction errors need their covariance structure; repeated clonal measurements can include permanent environmental or nonadditive effects. Overdispersion or shared environments can invalidate the Poisson observation model. The worked example establishes a reproducible route through the manuscript's estimation steps under stated assumptions, rather than a general-purpose estimator for every breeding dataset.

#### References


[1] Hadfield, J. D., A. J. Wilson, D. Garant, B. C. Sheldon and L. E. B. Kruuk (2010). The misuse of BLUP in ecology and evolution. *The American Naturalist* 175:116–125. `https://doi.org/10.1086/648604`.

[2] Morrissey, M. B. and I. B. J. Goudie (2022). Analytical results for directional and quadratic selection gradients for log-linear models of fitness functions. *Evolution* 76:1378–1390. `https://doi.org/10.1111/evo.14486`.
